# Host transcriptional programs underlying lesion development in contagious bovine pleuropneumonia

**DOI:** 10.64898/2026.06.24.734280

**Authors:** Arlind B. Mara, Angela Makumi, R. Grace Ozyck, Massimo Scacchia, Hezron Wesonga, Mark Ackermann, Noah O. Okumu, Winnie Chebore, Morgan Hunte, Jeremy M. Miller, Edan R. Tulman, Steven M. Szczepanek, Elise Schieck, Steven J. Geary

**Affiliations:** The Center of Excellence for Vaccine Research, Department of Pathobiology, University of Connecticut, Storrs, Connecticut, USA; The International Livestock Research Institute, Nairobi, Kenya; Istituto Zooprofilattico Sperimentale dell’Abruzzo e del Molise G. Caporale, Teramo, Italy; Kenya Agricultural and Livestock Research Organization, Nairobi, Kenya; Department of Veterinary Pathology, College of Veterinary Medicine, Iowa State University, Ames, Iowa, USA

**Author notes:** Corresponding Authors: SJG ES. Equal Contribution (Arlind B. Mara, Angela Makumi, R. Grace Ozyck).

## Abstract

Contagious bovine pleuropneumonia (CBPP), caused by *Mycoplasma mycoides* subsp. *mycoides (Mmm)*, remains a major burden to cattle health and the agricultural industry. *Mmm* is an atypical bacterial pathogen that appears to lack classical virulence factors that cause direct tissue injury (i.e. toxins), and little is known about the mechanisms driving its pathogenicity. The host immune response is believed to be implicated in CBPP pathology, though the molecular mechanisms underlying lesion initiation, progression and chronicity are poorly defined. Classical pathology describes a continuum of lung lesions starting from early inflammation to more mature necrotic lesions and formation of fibrotic sequestra. However, the host transcriptional response driving this potentially immunopathological progression during *Mmm* infection has never been resolved *in vivo*. Here, we performed lesion-stage-resolved transcriptomic profiling of pathological lung tissue collected from experimentally infected animals and compared to healthy lung tissue collected from unchallenged controls. Differential gene expression and functional enrichment analyses were used to identify biological pathways relevant to *Mmm* infection and pathological lesion formation. Early infection was dominated by interferon-stimulated genes and cytokine-responsive pathways, creating a primarily antiviral-like response environment despite the bacterial etiology. Red hepatization showed strong induction of neutrophil chemoattractants, epithelial remodeling markers, and early matrix-remodeling enzymes. Consolidation, spanning red and grey stages, was enriched for innate immune activation, leukocyte adhesion, extracellular matrix organization, and persistent interferon signaling. Grey hepatization reflected late-stage consolidation with heightened neutrophil effector activity, oxidative and proteolytic injury, and macrophage and fibroblast-linked collagen processing. Necrosis/Sequestra lesions showed reduced inflammatory signaling, robust extracellular matrix organization, adhesion, and morphogenetic pathways consistent with encapsulation and sequestrum formation. Our data indicate that the dynamic continuum of CBPP lung pathology is initiated by interferon-primed myeloid recruitment and amplified by neutrophil-driven injury and macrophage- and fibroblast-mediated matrix remodeling. These data further substantiate the role of dysregulated immunity in the development of disease during *Mmm* infection.

## INTRODUCTION

Contagious bovine pleuropneumonia (CBPP) is a severe and economically significant respiratory disease of cattle caused by infection with the atypical bacterial pathogen *Mycoplasma mycoides* subsp. *mycoides* (*Mmm*) (**1,2**). Disease presents primarily as an acute fibrinous pneumonia and in chronic cases develops into a fibrotic pleuropneumonia followed by the development of fibrotic pulmonary sequestra and pleural adhesions that significantly impair lung function and animal productivity. CBPP is a highly contagious disease with mortality rates of up to 50% (**1**). Given its impact on animal health and livestock production, CBPP is on the list of notifiable diseases of the World Organization for Animal Health (WOAH).

While CBPP is a devastating disease*, Mmm*, its etiologic agent, appears to lack classical virulence factors (e.x. toxins), and *Mmm* virulence has been primarily attributed to components that primarily play metabolic functions or modulate the immune response. (**1,3**). Hydrogen peroxide (H_2_O_2_) production as a result of glycerol metabolism has been hypothesized to serve as a primary virulence factor of *Mmm*, as it induces cell damage *in vitro* (**1,4**). The capsular polysaccharide galactan is also considered to be a major virulence factor as it has been shown to induce vascular and cytopathic effects after intravenous infection, and to modulate the host immune response (**1,5**). Immune-mediated lung injury is also thought to be a major contributor to CBPP pathology (**1**). Indeed, gross histopathological evaluation of CBPP lung lesions follows the classical progression of an inflammatory response as it progresses from pathogen recognition and initial inflammatory cell recruitment through wound repair and fibrosis. Gross lesions of this progression, prominent in lungs of affected animals, include red hepatization, consolidation, grey hepatization, necrosis, fibrosis, necrosis, and sequestra (**1**,**6**). Despite the recognized role that host immune responses play in the pathology of mycoplasma-induced disease (**1**), we know little about the immunological pathways activated in the lung during infection with *Mmm.* An understanding of the host responses associated with *Mmm* infection is crucial, as conserved responses can inform on pathogenetic mechanisms useful for development of safe and efficacious intervention strategies.

High-throughput transcriptomics provide an opportunity to profile such host responses. While a handful of studies have conducted transcriptional analyses of whole blood or peripheral blood mononuclear cells (PBMCs) collected from infected animals or following *ex vivo* exposures of PBMCs and lung explants to *M. mycoides* species (**7–9**), there are no reports examining host responses at the local site of respiratory infection and primary disease *in vivo*. Given that interactions of recruited leukocytes with the resident cell populations at the site of infection are critical for shaping the host response and disease outcome, it is critically important that these interactions are studied within an *in vivo* system. Our group has previously utilized RNAseq to probe *in vivo* host-pathogen interactions of *Mycoplasma gallisepticum* within tracheas of infected chickens (**10–12**). Here, we utilize the same concept to understand host-pathogen interactions of *Mmm* with the bovine lung in the context of *in vivo* infection and specifically of differential pathologies produced as lung lesions mature. We performed RNAseq on lung tissues with specific pathological features (Healthy-appearing (**B_HlthI**), Red Hepatization (**C_RedH**), Consolidation (**E_Cons**), Grey Hepatization (**D_GreyH**), and Necrosis/Sequestra (**F_NecSeq**)) collected from infected animals and compared transcriptional profiles to healthy lung tissue from uninfected animals (**A_HlthU**) (See **Supplementary Table S1** for sample metadata).

## RESULTS

### Quality control, global sample structure, summary of differentially expressed genes (DEGs) and summary of Gene Ontology enriched pathways

After processing and alignment to the *Bos taurus* ARS-UCD1.2 genome, the majority of samples (63/81 | 78%) passed QC thresholds with unique alignment >50% to the reference genome. QC distributions and per condition alignment rates are shown in **Supplementary Figures S1** and **S2**. Principal component analysis on variance-stabilized counts revealed a separation of samples by infection status and pathological condition consistent with an evolving lesion landscape, with different lesions forming their own clusters but with some expected overlap apparent between red hepatization, consolidation, and grey hepatization samples (**Supplementary Figure S3**). Sample-to-sample Euclidean distance heatmaps corroborated clustering by condition and infection status (**Supplementary Figure S4**), supporting the validity of downstream differential expression and gene set enrichment analyses. Differential gene expression analysis revealed several thousand genes that were differentially regulated across different different types of lesions (**Figure 1**). Compared to Healthy Uninfected tissue, a total of **162** genes were differentially expressed in Healthy Infected tissues (116 upregulated, 46 downregulated); **3160** genes in Red Hepatization (1607 upregulated, 1553 downregulated); **5370** genes in Grey Hepatization (2315 upregulated, 3055 downregulated); **4089** genes in Consolidation (1903 upregulated, 2186 downregulated); and **10,525** genes in Necrotic/Sequestra (7458 upregulated, 3067 downregulated).

**Figure 1.**
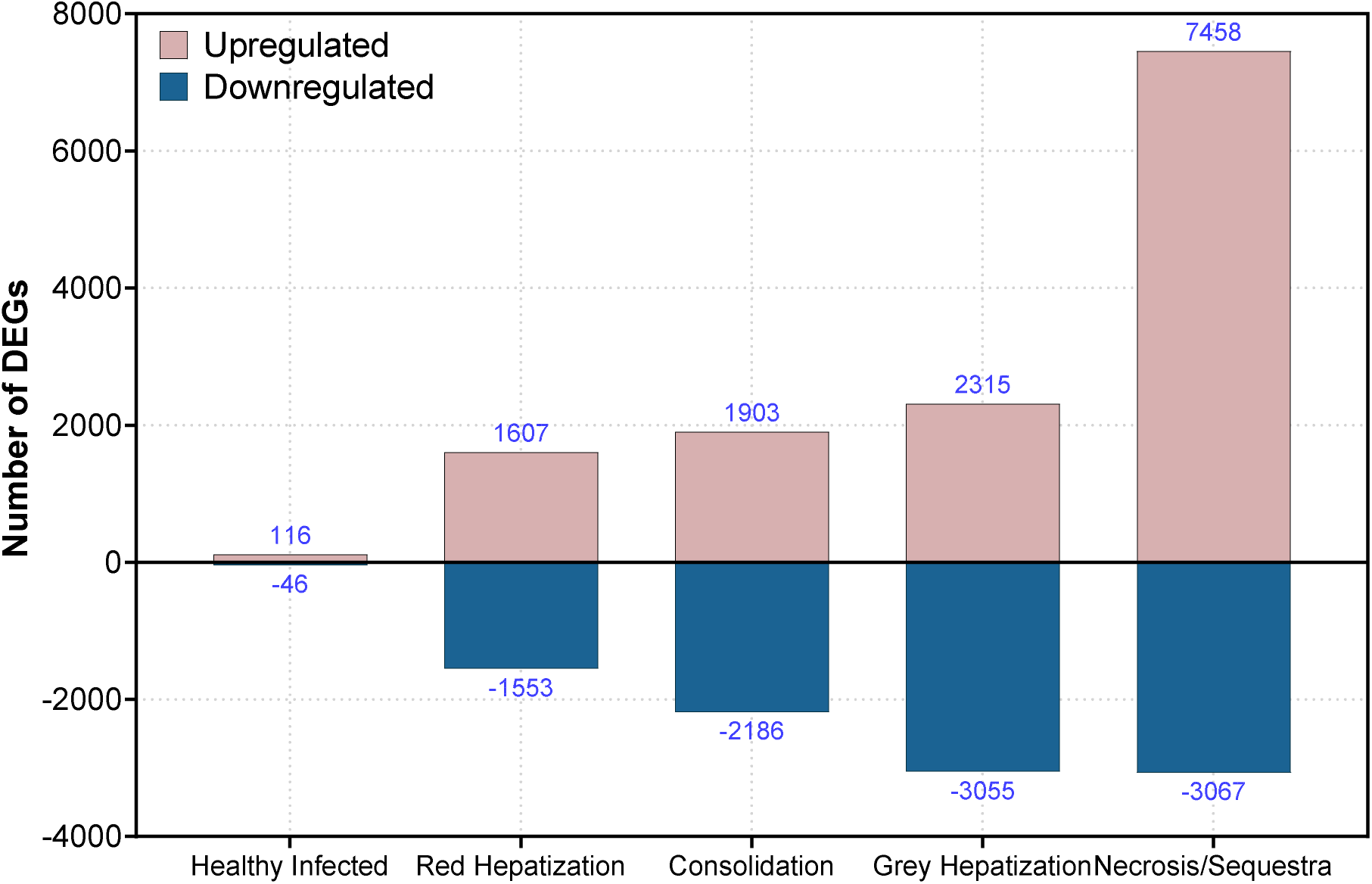
Number of significant differentially expressed genes per pathological condition as compared to healthy uninfected lung tissue.

Across all stage resolved comparisons, gene ontology analysis revealed a clear temporal progression in host biological responses, beginning with an initial dominant interferon mediated antiviral response signature, and culminating in structural remodeling and tissue reorganization (**Table 1**). In early lesions, enriched GO terms were overwhelmingly associated with *defense response to virus*, *response to biotic stimulus* and *regulation of viral processes*, despite the bacterial etiology of CBPP. This appears to be a consistent finding with members of the mycoides cluster, as investigations found *Mycoplasma mycoides* subsp. *capri* also induces an antiviral-like transcriptional profile when interacting with caprine leukocytes (**7**). As lesions advanced into Red Hepatization, Consolidation, and Grey Hepatization, antiviral signaling persisted but was increasingly accompanied by pathways related to *cytokine responses*, *inflammatory activation*, and *regulation of immune system processes*, consistent with the transition from acute exudation to a more regulated, yet highly active, inflammatory environment. Tissue remodeling pathways were also activated in these lesion stages. In contrast, necrotic tissue and sequestra exhibited a marked shift away from immune dominated signatures toward GO categories associated with *cell adhesion*, *actin cytoskeleton organization*, *cell-cell junction remodeling*, and broader morphogenetic processes. Although residual antiviral activity was observed, they were overshadowed by pathways reflecting fibrosis-driven structural remodeling.

**Table 1.**
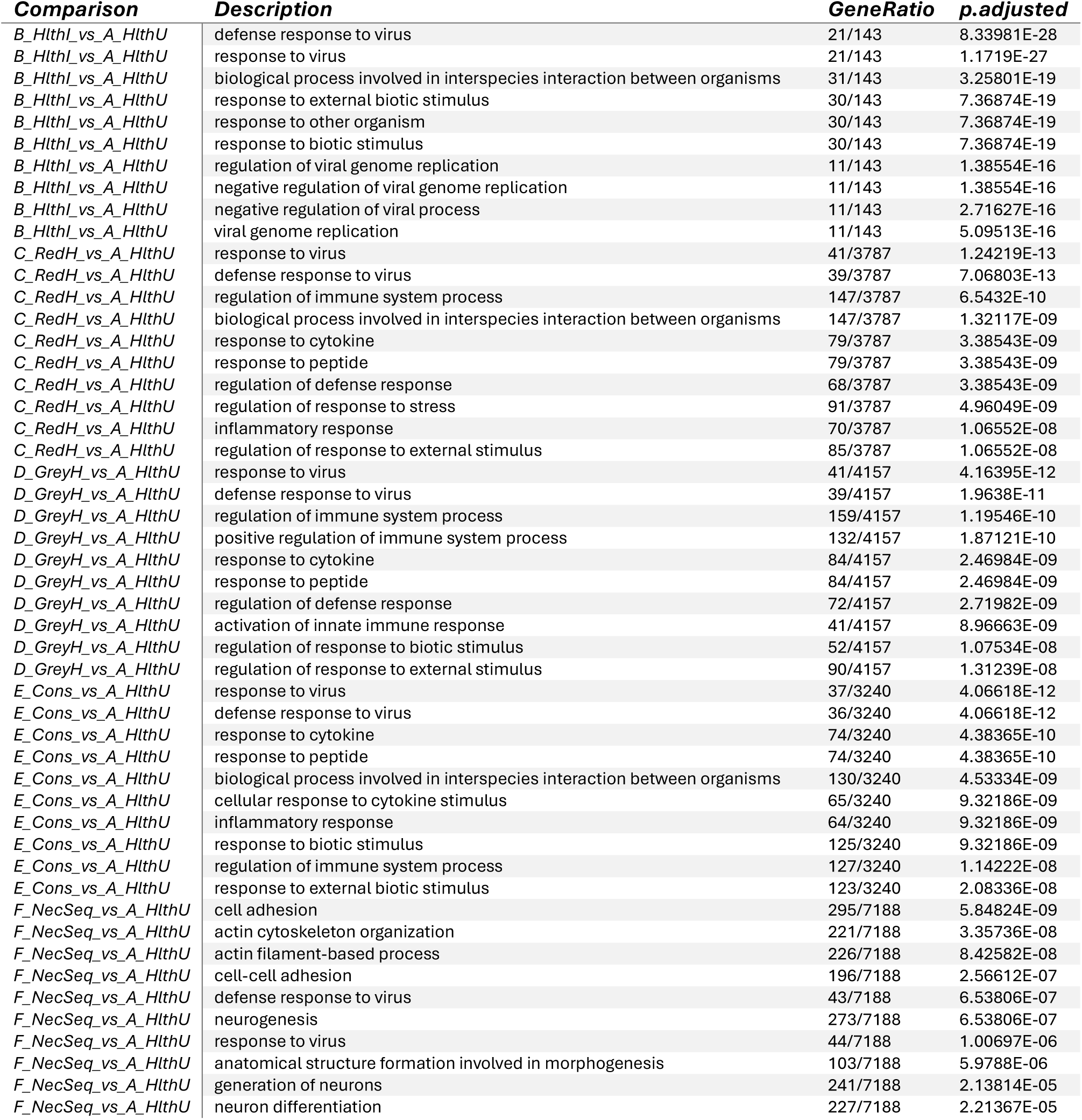
Summary of top 10 GO terms enriched by pathological condition compared to healthy uninfected control tissues. A_HlthU. : Healthy Uninfected, **B_HlthI**: Healthy Infected, **C_RedH**: Red Hepatization, **D_GreyH**: Grey Hepatization, **E_Cons**: Consolidation, **F_NecSeq**: Necrosis/Sequestra

| **Comparison** | **Description** | **GeneRatio** | **p.adjusted** |
| --- | --- | --- | --- |
| *B_HlthI_vs_A_HlthU* | defense response to virus | 21/143 | 8.33981E-28 |
| *B_HlthI_vs_A_HlthU* | response to virus | 21/143 | 1.1719E-27 |
| *B_HlthI_vs_A_HlthU* | biological process involved in interspecies interaction between organisms | 31/143 | 3.25801E-19 |
| *B_HlthI_vs_A_HlthU* | response to external biotic stimulus | 30/143 | 7.36874E-19 |
| *B_HlthI_vs_A_HlthU* | response to other organism | 30/143 | 7.36874E-19 |
| *B_HlthI_vs_A_HlthU* | response to biotic stimulus | 30/143 | 7.36874E-19 |
| *B_HlthI_vs_A_HlthU* | regulation of viral genome replication | 11/143 | 1.38554E-16 |
| *B_HlthI_vs_A_HlthU* | negative regulation of viral genome replication | 11/143 | 1.38554E-16 |
| *B_HlthI_vs_A_HlthU* | negative regulation of viral process | 11/143 | 2.71627E-16 |
| *B_HlthI_vs_A_HlthU* | viral genome replication | 11/143 | 5.09513E-16 |
| *C_RedH_vs_A_HlthU* | response to virus | 41/3787 | 1.24219E-13 |
| *C_RedH_vs_A_HlthU* | defense response to virus | 39/3787 | 7.06803E-13 |
| *C_RedH_vs_A_HlthU* | regulation of immune system process | 147/3787 | 6.5432E-10 |
| *C_RedH_vs_A_HlthU* | biological process involved in interspecies interaction between organisms | 147/3787 | 1.32117E-09 |
| *C_RedH_vs_A_HlthU* | response to cytokine | 79/3787 | 3.38543E-09 |
| *C_RedH_vs_A_HlthU* | response to peptide | 79/3787 | 3.38543E-09 |
| *C_RedH_vs_A_HlthU* | regulation of defense response | 68/3787 | 3.38543E-09 |
| *C_RedH_vs_A_HlthU* | regulation of response to stress | 91/3787 | 4.96049E-09 |
| *C_RedH_vs_A_HlthU* | inflammatory response | 70/3787 | 1.06552E-08 |
| *C_RedH_vs_A_HlthU* | regulation of response to external stimulus | 85/3787 | 1.06552E-08 |
| *D_GreyH_vs_A_HlthU* | response to virus | 41/4157 | 4.16395E-12 |
| *D_GreyH_vs_A_HlthU* | defense response to virus | 39/4157 | 1.9638E-11 |
| *D_GreyH_vs_A_HlthU* | regulation of immune system process | 159/4157 | 1.19546E-10 |
| *D_GreyH_vs_A_HlthU* | positive regulation of immune system process | 132/4157 | 1.87121E-10 |
| *D_GreyH_vs_A_HlthU* | response to cytokine | 84/4157 | 2.46984E-09 |
| *D_GreyH_vs_A_HlthU* | response to peptide | 84/4157 | 2.46984E-09 |
| *D_GreyH_vs_A_HlthU* | regulation of defense response | 72/4157 | 2.71982E-09 |
| *D_GreyH_vs_A_HlthU* | activation of innate immune response | 41/4157 | 8.96663E-09 |
| *D_GreyH_vs_A_HlthU* | regulation of response to biotic stimulus | 52/4157 | 1.07534E-08 |
| *D_GreyH_vs_A_HlthU* | regulation of response to external stimulus | 90/4157 | 1.31239E-08 |
| *E_Cons_vs_A_HlthU* | response to virus | 37/3240 | 4.06618E-12 |
| *E_Cons_vs_A_HlthU* | defense response to virus | 36/3240 | 4.06618E-12 |
| *E_Cons_vs_A_HlthU* | response to cytokine | 74/3240 | 4.38365E-10 |
| *E_Cons_vs_A_HlthU* | response to peptide | 74/3240 | 4.38365E-10 |
| *E_Cons_vs_A_HlthU* | biological process involved in interspecies interaction between organisms | 130/3240 | 4.53334E-09 |
| *E_Cons_vs_A_HlthU* | cellular response to cytokine stimulus | 65/3240 | 9.32186E-09 |
| *E_Cons_vs_A_HlthU* | inflammatory response | 64/3240 | 9.32186E-09 |
| *E_Cons_vs_A_HlthU* | response to biotic stimulus | 125/3240 | 9.32186E-09 |
| *E_Cons_vs_A_HlthU* | regulation of immune system process | 127/3240 | 1.14222E-08 |
| *E_Cons_vs_A_HlthU* | response to external biotic stimulus | 123/3240 | 2.08336E-08 |
| *F_NecSeq_vs_A_HlthU* | cell adhesion | 295/7188 | 5.84824E-09 |
| *F_NecSeq_vs_A_HlthU* | actin cytoskeleton organization | 221/7188 | 3.35736E-08 |
| *F_NecSeq_vs_A_HlthU* | actin filament-based process | 226/7188 | 8.42582E-08 |
| *F_NecSeq_vs_A_HlthU* | cell-cell adhesion | 196/7188 | 2.56612E-07 |
| *F_NecSeq_vs_A_HlthU* | defense response to virus | 43/7188 | 6.53806E-07 |
| *F_NecSeq_vs_A_HlthU* | neurogenesis | 273/7188 | 6.53806E-07 |
| *F_NecSeq_vs_A_HlthU* | response to virus | 44/7188 | 1.00697E-06 |
| *F_NecSeq_vs_A_HlthU* | anatomical structure formation involved in morphogenesis | 103/7188 | 5.9788E-06 |
| *F_NecSeq_vs_A_HlthU* | generation of neurons | 241/7188 | 2.13814E-05 |
| *F_NecSeq_vs_A_HlthU* | neuron differentiation | 227/7188 | 2.21367E-05 |

Looking broadly at the top differentially regulated gene signatures across pathological conditions, we see **15** downregulated transcripts and **25** upregulated transcripts that persist throughout the temporal progression of disease as defined by the different pathological states (**Figure 2**). The downregulated signature is dominated by loss of epithelial-basement membrane and tight junction programs. Multiple transcripts associated with structural integrity of the basement membrane were downregulated, including the collagen IV α5 chain *COL4A5*. The laminin γ2 chain *LAMC2*, and the laminin anchor receptor *ITGA3*, which together form the laminin-integrin-collagen axis required for stable epithelial anchoring and basement membrane organization, were also downregulated (**13–15**). Tight junction and epithelial adhesion molecules such as claudin 7 (*CLDN7)*, the junctional adhesion molecule A (*F11R*/*JAM-A*) and the EpCAM-like tumor-associated calcium signal transducer 2 (*TACSTD2* - *EpCAM/TROP2* family) were also decreased, consistent with impaired barrier function and reduced epithelial cohesion (**16,17**).Downregulation of other ECM organizing factors such as the latent-transforming growth factor β binding protein 2 (*LTBP2)*, the laminin related netrin-4 (*NTN4)* and the cytoskeleton-membrane linker chloride intracellular channel 5 (*CLIC5)* further point toward destruction of the epithelial barrier architecture during infection (**18–20**). Altogether, these changes indicate a coordinated collapse of epithelial barrier and basement membrane integrity during *Mmm* infection.

**Figure 2.**
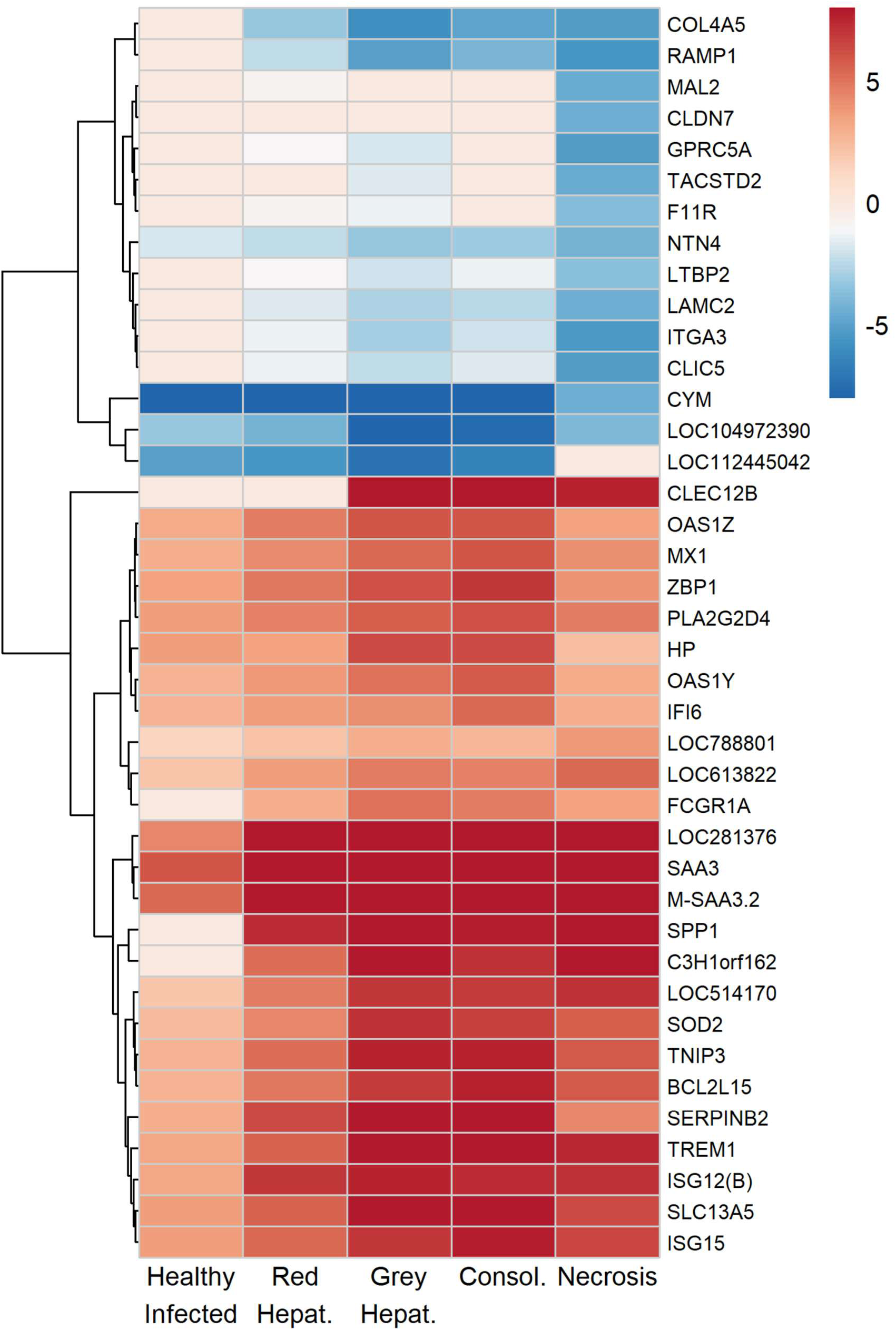
Top differentially expressed genes across pathological conditions. Heatmap reflects log2 fold-change values compared to healthy uninfected tissue from control animals.

Canonical interferon stimulated antiviral effectors such as *MX1, OAS1Z/Y, ZBP1, IFI6, ISG15* and *ISG12(B)* were strongly induced across all infected tissues, consistent with activation of the 2’-5’ oligoadenylate/Rnase L pathway, dynamin-like restriction of viral replication, and cytosolic DNA/Z-DNA sensing (**21–24**). *ZBP1* sensing of Z-nucleic acid has been shown to result in necroptotic cell death and to heighten the inflammatory response which may explains the areas of necrosis observed pathologically in the infected lungs, and the extreme inflammatory environment that is observed at the site of infection. Also upregulated by the infection were acute phase mediators such as haptoglobin (*HP*), serum amyloid A (*SAA3/M-SAA3.2*), and secreted phosphoprotein/osteopontin (*SPP1*) which are known to orchestrate hemoglobin scavenging, neutrophil recruitment, HDL remodeling and Th1-biased macrophage retention at the site of infection (**25**). Upregulation of the *FCGR1A* (CD64, high affinity Fc-gamma receptor) and the myeloid triggering receptor *TREM1* transcripts further supports the idea that the inflammatory landscape is composed of monocytes/macrophages poised for opsonophagocytosis (**26,27**). Collectively, these data indicate that as epithelial barrier integrity programs are suppressed, an interferon-driven antiviral and inflammatory myeloid axis becomes the dominant response. Overwhelming innate responses have been shown to contribute to lung pathology (**28**), and it is possible that these responses drive the severe and fibrotic lung damage that is observed in infected animals.

Superimposed on this inflammatory environment, several upregulated genes also suggest metabolic and regulatory tuning of the response. The upregulation of the mitochondrial superoxide dismutase *SOD2* is consistent with increased regulation of mitochondrial ROS production in activated cells, while *SLC13A5* upregulation indicates enhanced citrate import to support lipid and energy metabolism during activation (**29,30**). At the same time, upregulation of negative regulators such as *TNIP3* (*ABIN-3)* an A20-binding inhibitor of NF-κB, the inhibitory C-type lectin *CLEC12B*, the serine protease inhibitor *SERPINB2* (a.k.a plasminogen activator inhibitor type 2 or PAI-2), and the Bcl-2 family member *BCL2L15* indicates attempts to limit the inflammatory response (**31–34**). These transcriptional signatures should be further investigated as they may be of utility as biomarkers associated with CBPP disease severity and may inform development of downstream diagnostics.

### Lesion Stage Resolved Transcriptional Profiles: Healthy Infected versus Uninfected

Comparison of healthy lung tissue from uninfected controls and healthy appearing tissue from infected animals provides insights regarding the transcriptional programs that underpin the early host responses to infection. Surprisingly, despite the bacterial etiology of CBPP, the comparison between healthy infected and healthy uninfected lungs revealed strong enrichment of antiviral and innate immune responses, with the most significantly enriched Gene Ontology categories including **GO:0051607** - *defense response to virus*, and **GO:0009615** - *response to virus*, driven by a large cluster of interferon-stimulate gene transcripts (ISGs) (**Figure 3, Supplementary Table S2)** Processes related to *viral genome replication* and its regulation (**GO:0045069, GO:0045071, GO:0019079**) were also enriched. Interferon pathways were prominent, with enrichment of *interferon-mediated signaling pathway* (**GO:0140888**), *type I interferon-mediated signaling pathway* (**GO:0060337**) and *cellular response to type I interferon* (**GO:0071357**), underscoring the activation of canonical antiviral defense cascades. Other GO pathways such as *response to external biotic stimulus* (**GO:0043207**) and *response to other organism* (**GO:0051707**) reflect recognition of foreign microbial components, indicative of early responses. Further underscoring early responses was the upregulation of lipid mediator and acute phase pathways such as eicosanoid secretion (**GO:0032309**) and arachidonate transport (**GO:1903G63**).

**Figure 3.**
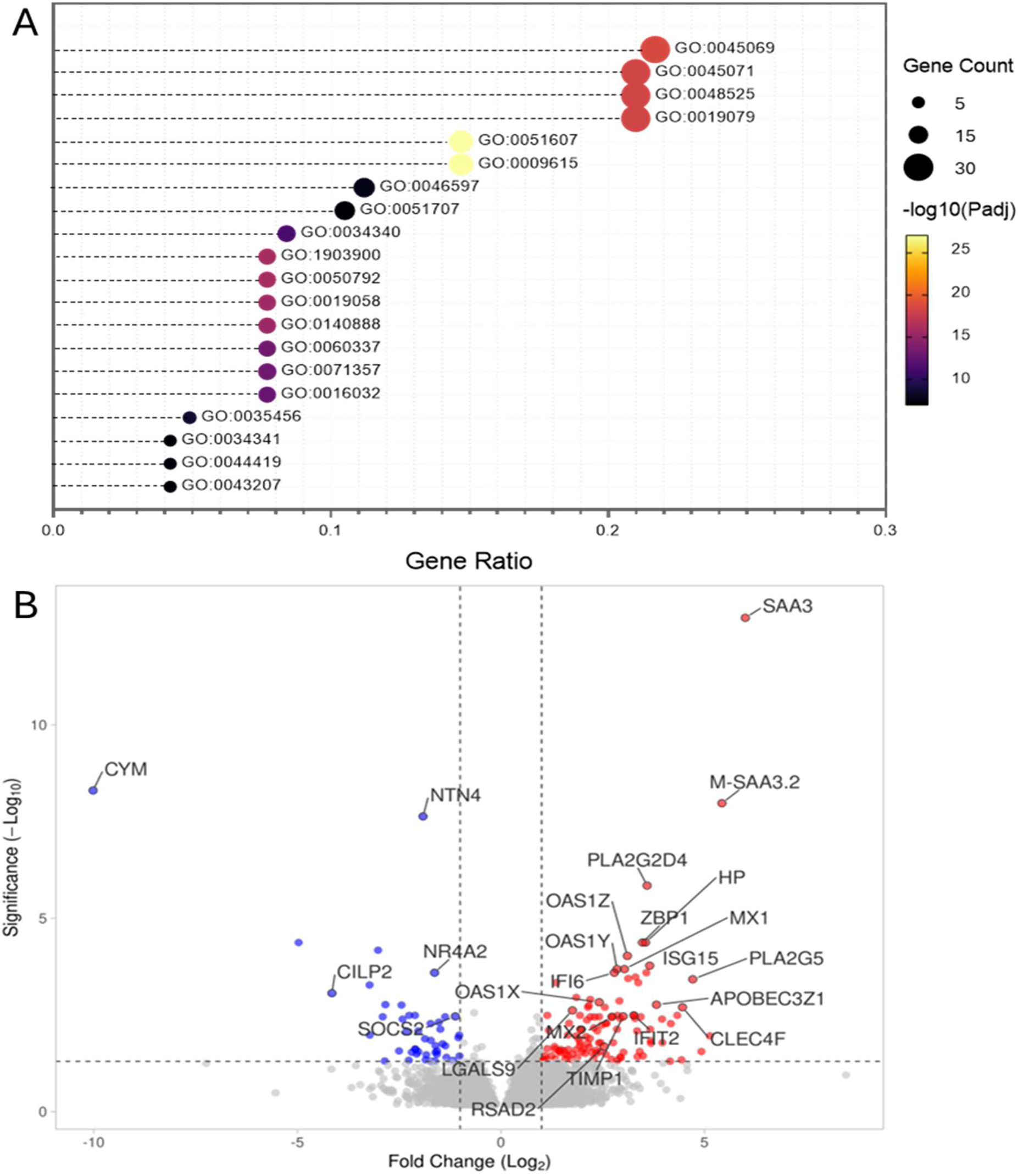
Transcriptional landscape of Healthy Infected tissues. **(A)** Top 20 enriched GO terms and **(B)** volcano plot highlighting differentially expressed genes with red marking significantly upregulated genes and blue marking significantly downregulated genes.

Within these enriched processes, the acute phase protein transcripts *SAA3*, *M-SAA3.2*, and haptoglobin (*HP*) were strongly upregulated, consistent with the rapid phase response to proinflammatory cytokines in bovine epithelia, reinforcing their role as biomarkers of bovine respiratory disease (Supplementary **Table S3**,) (**25,35–37**). Furthermore, upregulation of phospholipases *PLA2G5*and *PLA2G2D4* is indicative of activation of arachidonic acid metabolite secretion which points to early remodeling of lipid signaling that drive early inflammatory responses through prostaglandin and leukotriene production (**38**). These factors serve not only as markers of inflammation but also contribute to iron sequestration and neutrophil and macrophage modulation, creating an environment hostile to pathogens but also potentially contributing to immunopathology and bystander injury to lung tissue. The significant upregulation of *ZBP1*, a cytosolic sensor of Z-DNA/RNA that can trigger PANoptosis (pyroptosis, apoptosis, and necroptosis) may indicate that the host immune response contributes to lung pathology during *Mmm* infection (**3G,40**). A coordinated ISG cluster was evident, including *ISG15*, *MX1*, *MX2*, *OASI1Z/Y/X*, *OAS2*, *IFI6*, *RSAD2/viperin*, *IFIT2*, and *APOBEC3Z1* genes which mediate antiviral defense and immune modulation (**22**,**41–44**). *TIMP1*, an inhibitor of matrix metalloproteinases, was upregulated, suggesting early regulation of tissue integrity (**45–46**). *CHI3L1*, a chitinase-like protein linked to fibrosis and chronic inflammation was also upregulated, while downregulation of *NTN4* and *CILP2* indicated suppression of structural homeostasis of lung tissue and loss of barrier function (**47-49**). Furthermore, *LGALS9* and *CLEC4F* were upregulated while *SOCS2, NR4A2* was downregulated, consistent with enrichment of GO categories for negative regulation of T cell proliferation and negative regulation of lymphocyte activation, which highlights the suppression of the adaptive response during this early phase of innate dominance (**50–52**).

The upregulation of ISGs aligns well with the enriched GO categories of antiviral defense, confirming that infected but grossly normal lung tissue is already engaged in an antiviral transcriptional reprogramming due to *Mmm* infection, despite the bacterial etiology of CBPP. It is possible that these responses highlight a previously unappreciated intracellular niche for *Mmm* persistence in infected cattle. Importantly, however, type I interferons are also known to suppress adaptive lymphocyte proliferation and promote exhaustion during persistent infections (**23**). During Mycoplasma infection, these responses may favor pathogen persistence and disease. Strong activation of innate immunity damages lung tissue and remodels the lung environment while also blunting or delaying adaptive immune responses, potentially indicating that *Mmm* may induce these maladaptive immune responses as a way to avoid clearance and promote chronic persistence within the infected lung.

### Lesion Stage Resolved Transcriptional Profiles: Uninfected versus Red Hepatization

Red hepatization represents the first grossly visible stage of CBPP pathology and the next temporally resolved lesion following healthy appearing but infected lung tissue. Acute phase and interferon stimulated transcripts remained elevated and the GO pathways *response to virus* (**GO:0009615**) and *defense response to virus* (**GO:0051607**) remained enriched during Red Hepatization, reflecting continued persistence and recognition of *Mmm* infection (**Figure 4 and Supplementary Table S4).** Other broader immune categories such as *regulation of immune system process* (**GO:0002682**), *inflammatory response* (**GO:0006954**), *cell activation* (**GO:0001775**), and *leukocyte activation* (**GO:0045321**) were also enriched in Red Hepatization, highlighting expansion of both innate and adaptive immune pathways. Cytokine signaling was prominent, with enrichment of *response to cytokine* (**GO:0034097**) and *cytokine-mediated signaling pathways* (**GO:0019221**), underscoring the role of cytokine networks in orchestrating lesion development. Structural remodeling of lung tissue was evidenced through enrichment of *cell adhesion* (**GO:0007155**), *actin cytoskeleton organization* (**GO:0030036**), and *cell migration* (**GO:0016477**), consistent with leukocyte infiltration and tissue breakdown. Importantly, categories such as *positive regulation of immune system process* (**GO:0002684**) co-occurred with *negative regulation of lymphocyte activation* (**GO:0051250**), indicating a complex balance of immune stimulation and adaptive suppression that may be actively influenced by *Mmm* infection.

**Figure 4.**
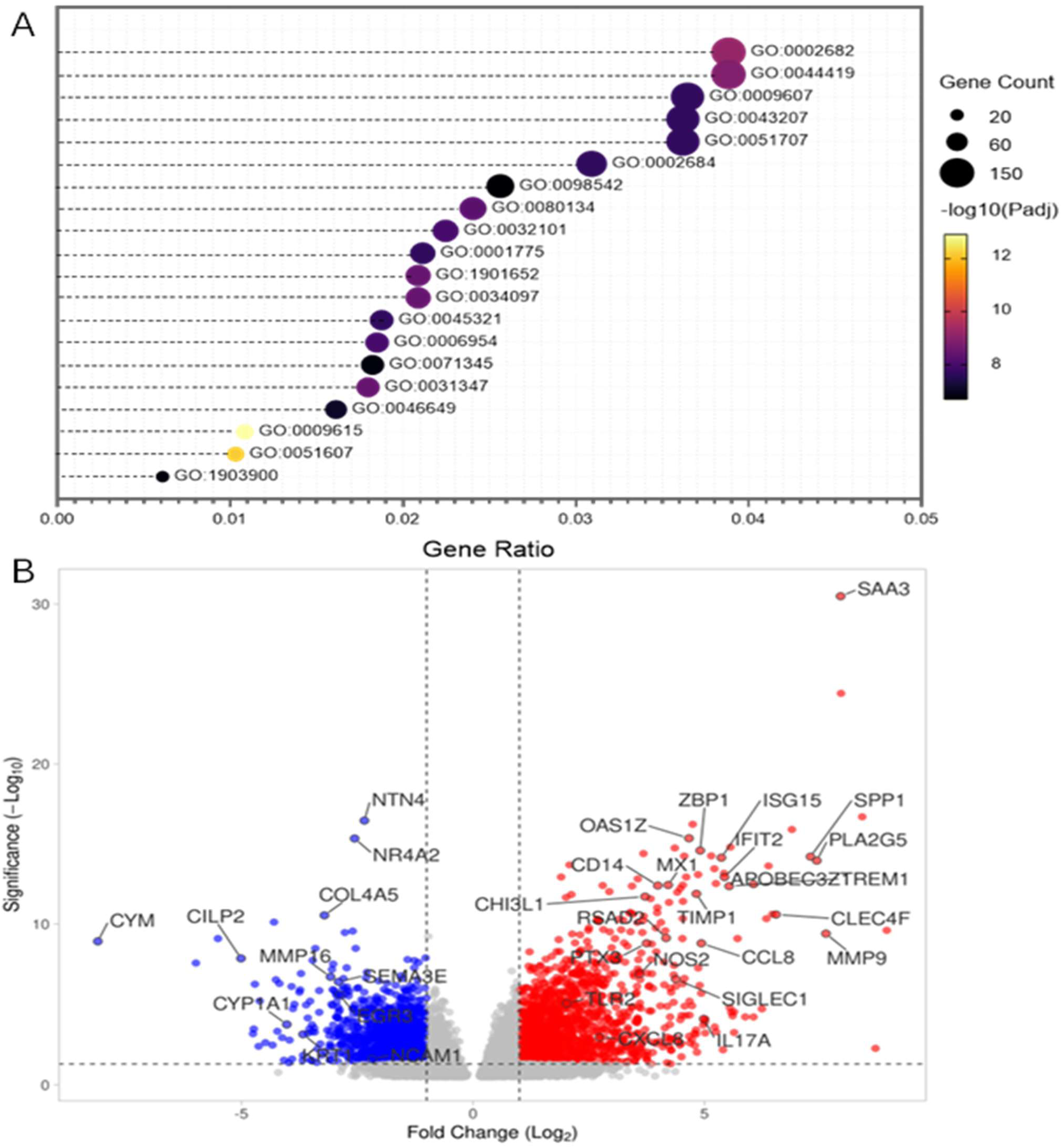
Transcriptional landscape Red Hepatized tissues. **(A)** Top 20 enriched GO terms and **(B)** volcano plot highlighting differentially expressed genes with red marking significantly upregulated genes and blue marking significantly downregulated genes

Acute phase proteins such as (*SAA3, M-SAA3.2, SAA2, SPP1*), ISGs such as (*ISG15, ISG12(B), MX1, MX2, OAS1X/Y/Z, IFIT2, RSAD2/viperin* and *ISG20*), lipid mediator signaling factors (phospholipases *PLA2G5, PLA2G2D4*) and other early transcripts such as *ZBP1, APOBEC3Z1,* and *TREM1* that were upregulated in the healthy infected tissue remained upregulated in Red Hepatization, indicating continued persistence and recognition of *Mmm* within the lesion (Supplementary **Table S5**). Inflammatory mediators such as *CCL8*, *CCL19*, *CCL20*, *CXCL2*, *CXCL5*, *CXCL8*, were upregulated, reflecting chemokine driven leukocyte recruitment, primarily of neutrophils, but also monocytes and T cells (**53**). *IL19, IL17A, IL27* and *IL21* were also upregulated, pointing to cytokine activation of Th17/Th1-associated immune pathways (**53**).

Matrix remodeling factors such as matrix metalloproteinases *MMP9*, *MMP1*, *MMP13* and *MMP25* were strongly upregulated, while structural and extracellular matrix genes such as collagen *COL4A4*, *COL4A5*, *COL4A6*, *COL15A1*, *COL21A1*, *COL25A1* and *COL2A1* were downregulated, reflecting collagen and basement membrane degradation and suppression of structural homeostasis that retains barrier integrity. *MMP15* and *MMP16* were downregulated, while *TIMP1*, a metalloproteinase inhibitor was upregulated, indicative of selective tissue remodeling (**54–56**). Chitinase-like protein transcripts *CHI3L1* and *CHI3L2* which are linked to fibrosis (**57,58**) were upregulated and can help explain the profibrotic nature of CBPP pathology. Cell adhesion, migration, and synapse formation transcripts such as *ISLR*, *SEMA3E*, *OPCML*, *NCAM1*, *NLGN1* and *KRT1* were downregulated, reflecting epithelial remodeling, and loss of normal lung function (**59-64**). Transcripts such as *SIGLEC1*, *CLEC4F*, *CD14*, *LBP*, *PTX3*, *NOS2*, *SOD2*, *TLR2*, and *TLR7*, associated with enhanced pathogen recognition, opsonization, reactive oxygen species and neutrophil effector function were upregulated, while metabolic pathway transcripts involved in xenobiotic metabolism such as *CYP1A1, CYP2A13, CYP2C90* and *CYP3A24* were downregulated, indicating loss of normal lung function and a shift toward inflammatory processes (**65–72**). Despite upregulation of chemokines involved in the recruitment of lymphocytes, several transcription factors linked to adaptive immunity were downregulated (*NR4A1*, *NR4A2*, *NR4A3*, *EGR3*), indicating suppression of T and B cell activation and downstream adaptive immunity (**73,74**), which may further support sustained persistence of *Mmm* in infected lungs. Taken together, these transcriptional signatures suggest that Red Hepatization continues to be a stage dominated by amplification of innate responses, continued restrain of adaptive immune responses, and initiation of tissue remodeling.

### Lesion Stage Resolved Transcriptional Profiles: Uninfected versus Consolidation

Transcriptionally speaking, Consolidation appeared to represent an intermediate lesion stage between Red and Grey Hepatization, where the acute inflammatory landscape begins to coexist with early adaptive immune activation and structural remodeling. *Pathogen recognition* (**GO:0002221, GO:0032496, GO:0002237**), *antiviral* (**GO:0009615, GO:0051607, GO:0045069, GO:0045071, GO:1903900**) and *cytokine/interferon mediated transcriptional signatures* (**GO:0034097 GO:0140888, GO:0060337, GO:0071357**) that were apparent in the earlier lesions remained enriched in consolidated lesions, indicating continued persistence and recognition of *Mmm (***Figure 5, Supplementary Table S6**). A pronounced ISG transcriptional signature remained evident, with strong upregulation of genes such as *ISG12(B), ISG15, ISG20, IFIT2, IFIT3, IFIT5,IFIC, IFI1C, IFI44, IFI44L, OAS1X/Y/Z, OAS2, MX1, MX2. RSAD2, USP18, XAF1, CMPK2, and EPSTI1*(**Supplementary Table S7**) (**44, 75**). Transcripts for pattern recognition receptors such as *TLR2, TLR7, NOD1, NOD2, IFIH1 (MDA5), CGAS, AIM2, NLRP1/2/3/12, ZBP1, DHX58 and NLRC4* (**76,77**) were upregulated and were accompanied by high-expression of myeloid-associated receptors such as *SIGLEC1, CLEC4D/E/F, CLEC12A/B, CLEC10A and CLEC5A*, consistent with enhanced pathogen recognition and phagocytosis (**78,79**). Chemokines and neutrophil-associated transcripts were also highly expressed, including recruiting factors such as *CXCL2, CXCL3, CXCL5, CXCL8, CXCR1, CXCR2* and activity factors such as *S100A8, S100A9, S100A12, LCN2, OLFM4, NCF1/4 and CYBB*, indicating active neutrophil recruitment and effector function (**80–82**).

**Figure 5.**
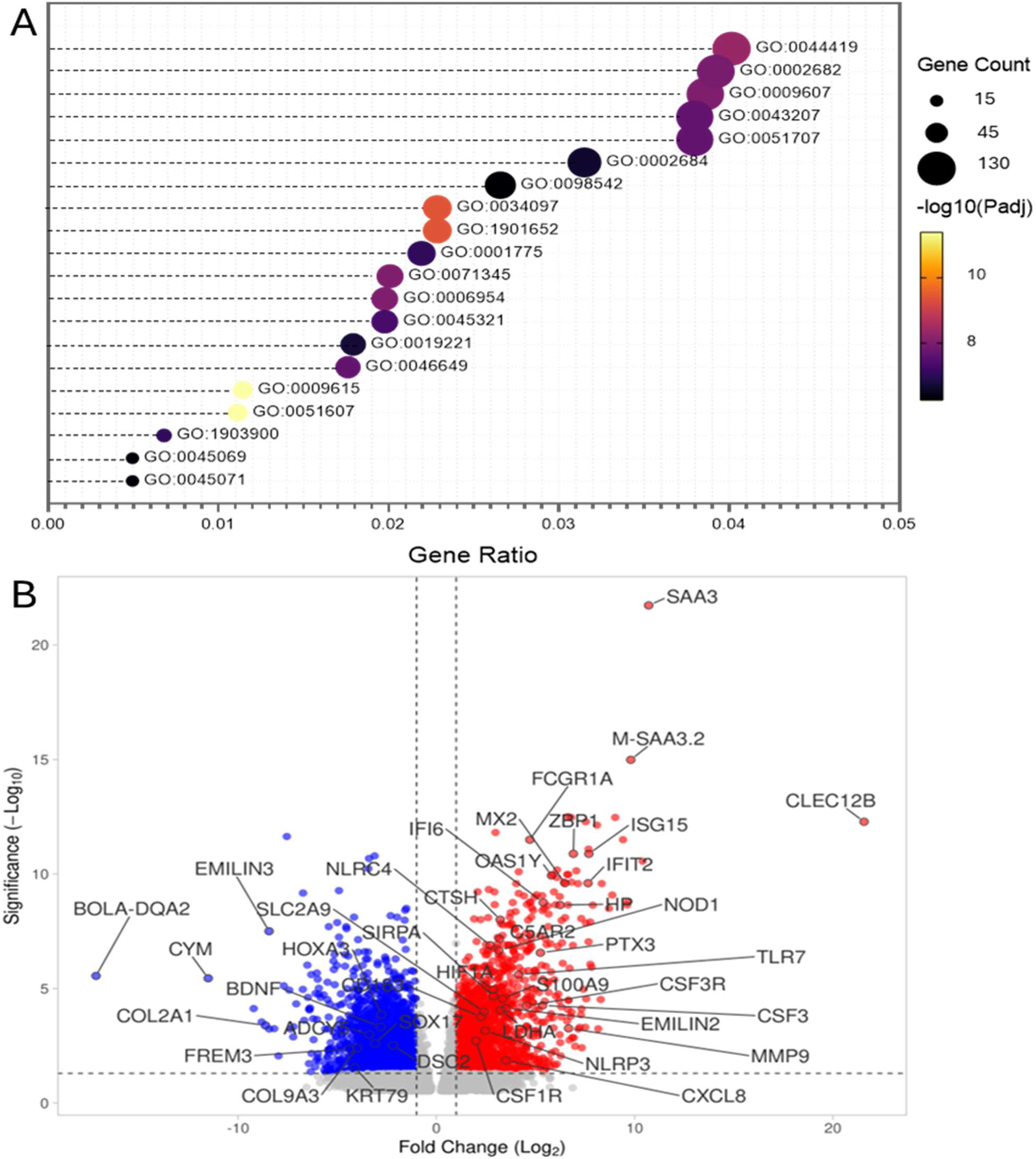
Transcriptional landscape of Consolidated tissues. (**A**) Top 20 enriched GO terms and (**B**) volcano plot highlighting differentially expressed genes with red marking significantly upregulated genes and blue marking significantly downregulated genes.

In addition to these inflammatory cues, pathways associated with cell adhesion and early structural remodeling were also enriched in Consolidation lesions (**GO:0098609** – *cell-cell adhesion*, **GO:0007159** – *leukocyte cell-cell adhesion*, **GO:0030198** – *extracellular matrix organization*, and **GO:0045229** – *external encapsulating structure organization*), indicating the beginning of the development of characteristic fibrotic lesions and sequestra that are observed with CBPP. Macrophage activation markers (*CDC8, CD1C3, CSF1R, CSF3R, CSF3, FCGR1A, FCGR2B, SIRPA, SIRPB1/2, TREM1 and TREM2*) (**83–85**) and acute-phase/complement genes (*SAA2, SAA3, HP, HPX, LBP, PTX3, C2, C5AR1, C5AR2*) (**25, 69, 86**) were upregulated, consistent with heightened phagocytic and complement activity. The increase in macrophage associated transcripts also signals the initiation of profibrotic processes and structural remodeling, which was further reflected by the upregulation of *MMP1/9/25, TIMP1, CTSL, CTSK, CTSS, VCAN, MXRA5, EMILIN2, PLOD1*, and *P4HA1*, indicating loss of lung architectural integrity (**56,87,88**). Hypoxia-linked metabolic reprogramming was also indicated by upregulated *HIF1A, EGLN3, BNIP3, LDHA, HK2, SLC2A1* and *SLC2A3*, further highlighting loss of normal lung function (**89–91**). This was further supported by loss of transcripts associated with epithelial cell identity, cilia and airway function, and structural components of healthy lung tissue such as *SFTPC, AQP5, AGER, BPIFA2A, COL2A1, COL9A1/2/3. COL4A2/3/4/5/6, HAPLN1, FREM2, FREM3, PCDH20, DSC2, KRT79* (**92–94**). Many neuronal associated transcripts (*STMN2, GRIN2B, NTRK2, BDNF, ADCY5*) and developmental regulators (*SOX7/17. FOXO4/C, HOXA3/5*) were also downregulated (**95–97**). Together, these transcriptional signatures indicate that Consolidation represents a stage characterized by intense innate immune activation, interferon signaling, neutrophil-driven inflammation and damage, macrophage recruitment, and progressing extracellular matrix and lung architecture remodeling that result in lung pathology and loss of normal organ function.

### Lesion Stage Resolved Transcriptional Profiles: Uninfected versus Grey Hepatization

Gene set enrichment analysis of Grey Hepatization revealed a transcriptional landscape that still retains the strong innate immune imprint seen in Red Hepatization and Consolidation, evidenced by the persistent enrichment of *response to virus* (**GO:0009615**), defense response to virus (**GO:0051607**), *pattern-recognition receptor signaling* (**GO:0002221**), and *inferferon mediated signaling* (**GO:0140888**, **GO:0060337**), which is expected given the chronic nature of *Mmm* infection (**Figure 6, Supplementary Table S8**). Unlike Consolidation, however, the transcriptional profile of Grey Hepatization shows a shift toward immune regulation and resolution, with pathways related to *negative regulation of immune system process* (**GO:0002683**) and *negative regulation of signaling* (**GO:0023057**) being highly enriched. Concurrently, Grey Hepatization displays a stronger emphasis on tissue remodeling and structural reorganization, reflected in the enrichment of pathways relating to *extracellular matrix organization* (**GO:0030198**), *extracellular structure organization* (**GO:0043062**), *external encapsulating structure organization* (**GO:0045229**) and *actin cytoskeleton organization* (**GO:0030036**). The enrichment of these pathways marks the transition from the exudative, neutrophil-rich environment of Red Hepatization and Consolidation, toward the macrophage and fibroblast driven restructuring the defines the fibrotic nature of CBPP pathology. In addition, pathways related to *angiogenesis* (**GO:0001525**), b*lood vessel morphogenesis* (**GO:0048514**) and *circulatory system development* (**GO:0072359**) are far more prominent in Grey Hepatization than in Consolidation, consistent with the vascular remodeling that may result in granulation tissue formation following this stage.

**Figure 6.**
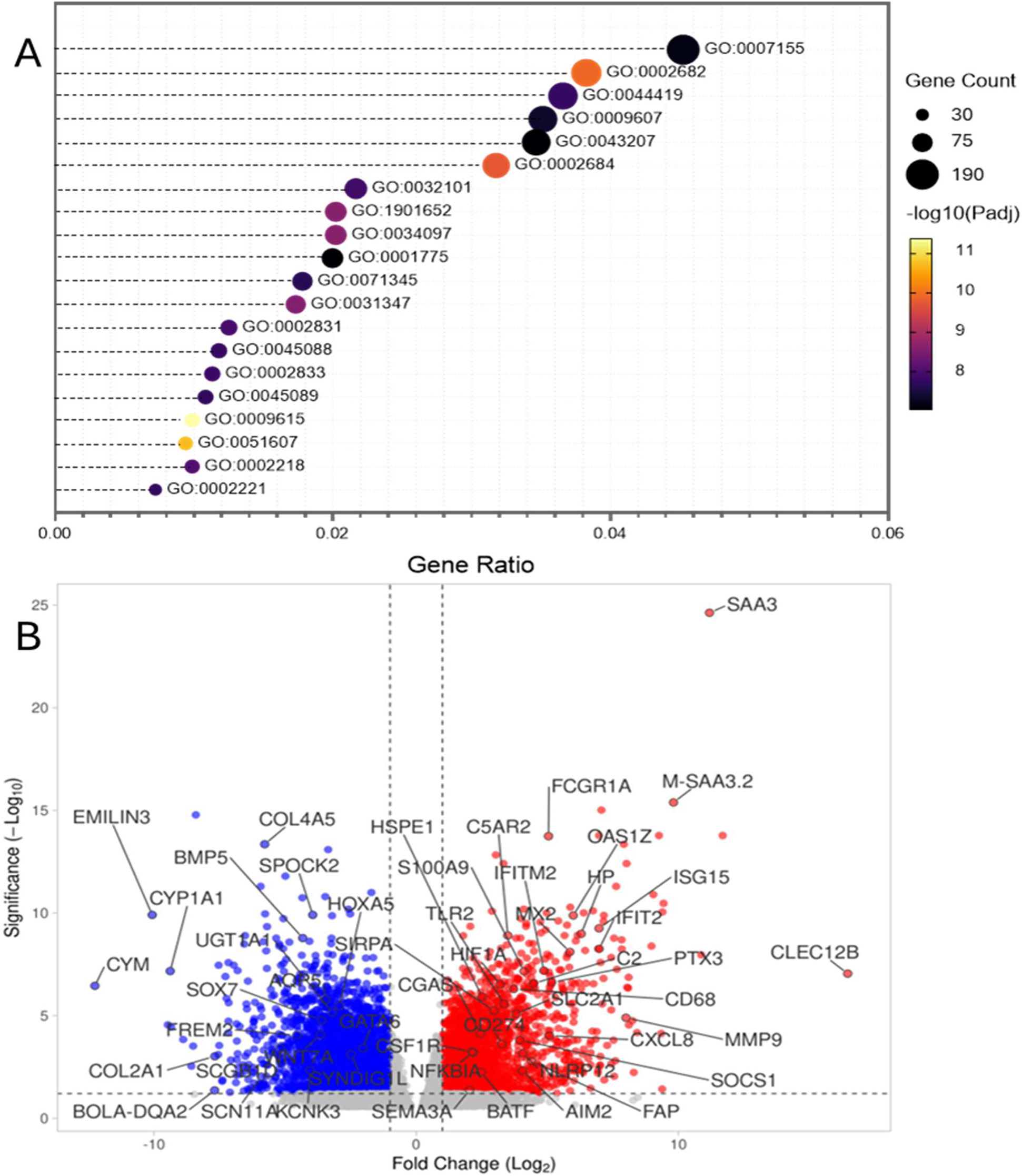
Transcriptional landscape of Grey Hepatized tissues. (**A**) Top 20 enriched GO terms and (**B**) volcano plot highlighting differentially expressed genes with red marking significantly upregulated genes and blue marking significantly downregulated genes.

At the gene level, Grey Hepatization lesions exhibited transcriptional programs that retained many of the innate immune features observed in previous stages but with a clear shift toward immune modulation, transition to a macrophage and fibroblast dominated environment and structural remodeling of lung tissue (Supplementary **Table S9**). Upregulated pattern-recognition receptors (*TLR2, TLR7, IFIH1, CGAS, AIM2, ZBP1, DHX58* and *NLRP12*) and ISGs (*ISG15, ISG12, IFIT2, IFIT3, IFITM1/2/10, MX1, MX2, OAS1X/Y/Z, OAS2, RSAD2, USP18, XAF1, CMPK2*, and *EPSTI1*) indicate that pathogen sensing and interferon mediated pathways remain active, indicative of persistent infection despite an overwhelming inflammatory response (**77,42–44**). However, the concurrent upregulation of regulatory mediators including *SOCS1/3, TNFAIP3, TNIP1/3, NFKBIA, DDIT4* and *IKBKE* suggest that the inflammatory pathways are increasingly counterbalanced by negative feedback circuits characteristic of resolution of inflammatory responses (**98–102**). However neutrophil-associated transcripts (*CXCL2, CXCL3, CXCL5, CXCL8, CXCR1, CXCR2, S100A8, S100A9, S100A12, LCN2, OLFM4, NCF1/4* and *CYBB)* remained elevated, indicating that neutrophil activity persists into this mature pathological lesion (**80–82**). Macrophage activation markers such as *CDC8, CD1C3, TREM1, TREM2, CSF1R, CSF3R, FCGR1A, FCGR2A/B, SIRPA, SIRPB1, TYROBP* and *OSMR* are strongly upregulated, reflecting the transition toward a macrophage-rich environment that orchestrates tissue remodeling and clearance of debris (**83–85**). Acute phase and complement opsonization transcripts (*SAA2, SAA3, HP, HPX, LBP, PTX3, C2, C9, C5AR1* and *C5AR2*) remained elevated, indicative of ongoing complement activation and phagocytic clearance (**25,69,86**).

Extracellular matrix (ECM) remodeling and stromal activation pathways were prominent in Grey Hepatization, with strong upregulation of *MMP1, MMP9, MMP13, MMP14, MMP25, CTSL, CTSK CTSH, CTSZ, TIMP1, VCAN, MXRA5, EMILIN2, CTHRC1, PLOD1, PLOD2, P4HA1*, COL-related regulators (**87–89**), and fibroblast associated transcripts such as *FAP*, *TGFBI*, *SULF1*, and *MEDAG* (**103–106**). These changes align with the enrichment for GO pathways for *extracellular matrix organization*, *extracellular structure organization*, and *external encapsulating structure organization*, marking the transition toward dense, airless, structurally reorganized fibrotic tissue that is characteristic of this lesion stage, and eventually leads to encapsulated sequestra.

Vascular remodeling signatures were also prominent. Upregulated angiogenesis-associated transcripts included *SEMA3A, SEMA4D, PROK2, FGF23, IL11, IL21R, GDF15, GUCY2D, VCAM1, NID2, THBS2, CREB3L1, RASA4B* and *ETV3L,* which indicates neovascular remodeling as the lung transitions from exudative inflammation toward structural reorganization and fibrotic remodeling that impairs normal lung function and is characteristic of CBPP pathology (**107–111**). Consistent with these changes, metabolic and hypoxia-associated genes such as *HIF1A, EGLN33, BNIP3, LDHA, HK2, ENO1, ALDOC, SLC2A1* and *SLC2A3* were upregulated, indicating persistent metabolic reprogramming within hypoxic damaged lung tissue (**112–115**). Additional stress-response and survival genes such as *DDIT3 (CHOP)*, *GDF10*, *SESN2*, *HSPH1*, *HSPA1A* and *HSPE1* further indicate that the lung microenvironment is undergoing damage from oxidative stress, nutrient deprivation and cellular turnover (**116–119**).

Grey hepatization was also characterized by a sweeping downregulation of transcripts associated with alveolar epithelial identity, surfactant biology, metabolic specialization, structural integrity, and neuro-epithelial signaling, which reflected profound architectural collapse and extreme loss of functional integrity of affected lung tissue. Downregulation of transcripts such as *SFTPC, AQP5, SCNN1B, CLDN18, CLDN5, AGER, SLC34A2, SLC22A17, SLC22A31, SLC9A4, SLC5AS, SLC1CA11*, and *SLC45A1* indicates loss of pneumocyte maintenance programs and destruction or dedifferentiation of epithelial cells and replacement of alveoli with dense proteinaceous and fibrinous scar tissue (**120–126**). Downregulation of *BPIFA2A, SCGB1D, SCGB2A2, MUC4*, and *SPINK5* further supports loss of secretory and barrier forming epithelial cells (**127–131**). Downregulation of components that mediate maintenance of extracellular matrix and basement membrane architecture such as *COL2A1, COL4A1/3/4/5, COLCAC, COLSA1/2/3. COL11A2, COL13A1, COL15A1, COL21A1, COL24A1, COL25A1, COL26A1, COL27A1, COL5A3, COLQ, FREM1/2/3. HMCN1/2, SPOCK1/2, WFDC1, WFIKKN2, PRELP, EMID1, EMCN* and *MXRA7* further reflect degradation of the alveolar scaffold and replacement with disorganized fibrotic matrix (**132–138**), a hallmark of CBPP pathology.

A suppression of developmental and morphogenetic regulators also indicates the collapse of epithelial-mesenchymal signaling networks that maintain tissue architecture with transcripts such as *SOX4/7/13/17, HOXA2/3/5, HOXB3/5/6, TBX1/2/3/4, TCF21, PRDMC, PRDM1C, GATA3/5/6, IRX1/2/3/5, NKD1/2, AXIN2, WNT2/7A/8B/9A/11, BMP5/6, BMPER, FGF1/12/20, FGFR1/2/3/4* and *NOTUM* being downregulated in grey hepatized tissue (**138–148**). Downregulation extended into neuronal, neuroendocrine, and synaptic signaling pathways with decreased expression of *STMN2, GAP43, NTM, GRIN1/2B/3A, CHRNB3/4, CNR1/2, ADCY5, ADCYAP1R1, PDYN, CRHR2, GALR3, GPR37/55/ASP1/ASP2, SYT7/8/13/25B, RIMS1/3, SCGN* and *SYNDIG1L* (**95**).

Further supporting loss of lung function is the downregulation of transcripts encoding factors involved in surfactant metabolism, xenobiotic detoxification, lipid handling, with *UGT1A6/2A1*, *HPGD*, *ABCC6*, *ABCB1*, *CYP1A1/1A2/2A13/2C87/2C90/4A22/4B1/17A1*, *FM01/5, GSTA1/2/3, GSTT2* and *GSTM3*, which reflect the collapse of epithelial metabolic capacity and the inability of damaged alveoli to maintain surfactant composition, lipid turnover, or oxidative homeostasis (**72**). Furthermore, downregulation of ion channels, transporters, and signaling receptors (*SCN11A/3B, KCNK3/B2/D3/H2*, *CNGA2*, *TRPV4/M4/C3*, *P2RX1/5*, *ASIC2/3*. *AQP1/5, SLC1CAS/5AS/24A3/6A1/8A3,22A31*) reflects the functional shutdown of epithelial transport capacity and the loss of barrier physiology (**149–156**).

In the immune axis, numerous immune modulatory and lymphocyte associated transcripts were differentially regulated. Transcripts like *IL7R, IL23R, CCR1, CCR2*, *CCR5*. *CXCL10*, *LAG3*, *CTLA4*, *CDCS*, *CD274* (PD-L1), *BATF, SPI1, POU2F2*, and *PRDM1* reflect a shift toward adaptive immune engagement and regulation, aligning with the enriched GO pathways for T cell activation, B cell activation, lymphocyte proliferation and regulation of lymphocyte activation (**53,157–164**). This indicates that adaptive immunity becomes more prominent during Grey Hepatization as innate neutrophilic responses begin to resolve. Downregulation of genes associated with antigen presentation such as *BOLA-DQA2*/*DQA5*, *CD1A/B* indicates suppressed cycling of antigen presentation machinery which supports adaptive immune activation (**165–167**).

Taken together, the transcriptional profile of Grey Hepatization reveals a lung environment undergoing epithelial collapse, loss of specialized metabolic and structural integrity programs, loss of neuronal and developmental processes, and dismantling of normal extracellular matrix architecture, underscoring a transition toward a structurally reorganized, fibrinous, and functionally impaired tissue state.

### Lesion Stage Resolved Transcriptional Profiles: Uninfected versus Necrosis

Necrotic tissue from fibrotic enclosed sequestra represents the most mature, chronic lesion produced by Mmm infection. A large proportion of the most strongly upregulated transcripts in this lesion corresponded to unannotated or partially annotated loci (*LOC*-). Because limited functional annotation exists for these transcripts, their biological role in the context of *Mmm* infection is difficult to interpret. However, their high fold-change values and consistent enrichment among the top differentially expressed genes may indicate that substantial components of the sequestrum-associated transcriptional response remain uncharacterized at the gene level. As such, these loci may represent important targets for future annotation and functional investigation, especially in the context of chronic tissue remodeling and immune modulation.

Pathway analysis on annotated DEGs from necrotic tissue displayed a transcriptional profile enriched by *cell adhesion*, *actin cytoskeleton organization* and *cell-cell junction pathways* (**GO:0007155, GO:0030036, GO:0098609**), consistent with the formation of dense, fibrotic capsules that physically isolate necrotic lung tissue (ie. Sequestra) (**Figure 7, Supplementary Table S10**). Upregulated transcripts included multiple profibrotic and matrix modifying enzymes (*MMP1/3/9/13/12/17/19/25*) (**54–56**) as well as adhesion molecules (*CDH9/18, CDHR2, PCDH8/15, CNTN2* and *CNTNAP2)* (**168,169**) (Supplementary **Table S11**). Enrichment of morphogenetic processes such as *anatomical structure formation*, *tube morphogenesis*, *vasculature development*, and *regulation of anatomical structure morphogenesis* (**GO: 0048646, GO:0035239, GO001944, GO:0022603**) is indicative for activation of fibroblast driven tissue remodeling programs that culminate in sequestrum development, and reorganize peripheral architecture. This was further supported by enrichment of *cell morphogenesis*, *cell projection organization*, and *actin filament dynamics* (**GO:0000902, GO:0031344, GO:0030041/42**), indicating ongoing cytoskeletal reorganization within fibroblasts, endothelial cells and residual leukocytes as the lesion matures. Transcription factors such as *HOXA10/1,1 HOXC8/9, HOXD10/13, PAX1/2/3/C/8, POU3F1/2/3/, DLX1/3/4/5*, and *BARX1* were upregulated alongside development-linked genes (*WNTC/SB/10A, RSPO4, BMP1, GDF15*) (**140,145,146**), indicating that certain repair processes that may retain tissue function are also induced. Pathways related to *angiogenesis* and *blood vessel morphogenesis* (**GO:0001525, GO:0048514**) further support the development of new vasculature at the sequestrum periphery where granulation tissue persists, further indicating an active effort to restore normal function. This is evidenced by transcripts such as *ANGPTL6*, *VEGF*-responsive channels (*KCNK9, KCNH4*), *ECM1*, *COL22A1, COL19A1, COL23A1*, and *MXRA5* (**87–89, 150**) being upregulated. Newly evident in these mature lesions was the upregulation of numerous transcripts associated with cell cycle progression, DNA replication, and cell proliferation (*CCNB1, CCNB2, CDC20, CDC6, MCM10, BIRC5, AURKA/B, PLK1, KIF20A/18B*), which indicates ongoing turnover of stromal and immune cells within the lesion associated with healing and repair processes exemplified by the enrichment of *regulation of multicellular organismal processes* (**GO: 0051239**) and *regulation of developmental growth* (**GO:0048638**) (**170,171**).

**Figure 7.**
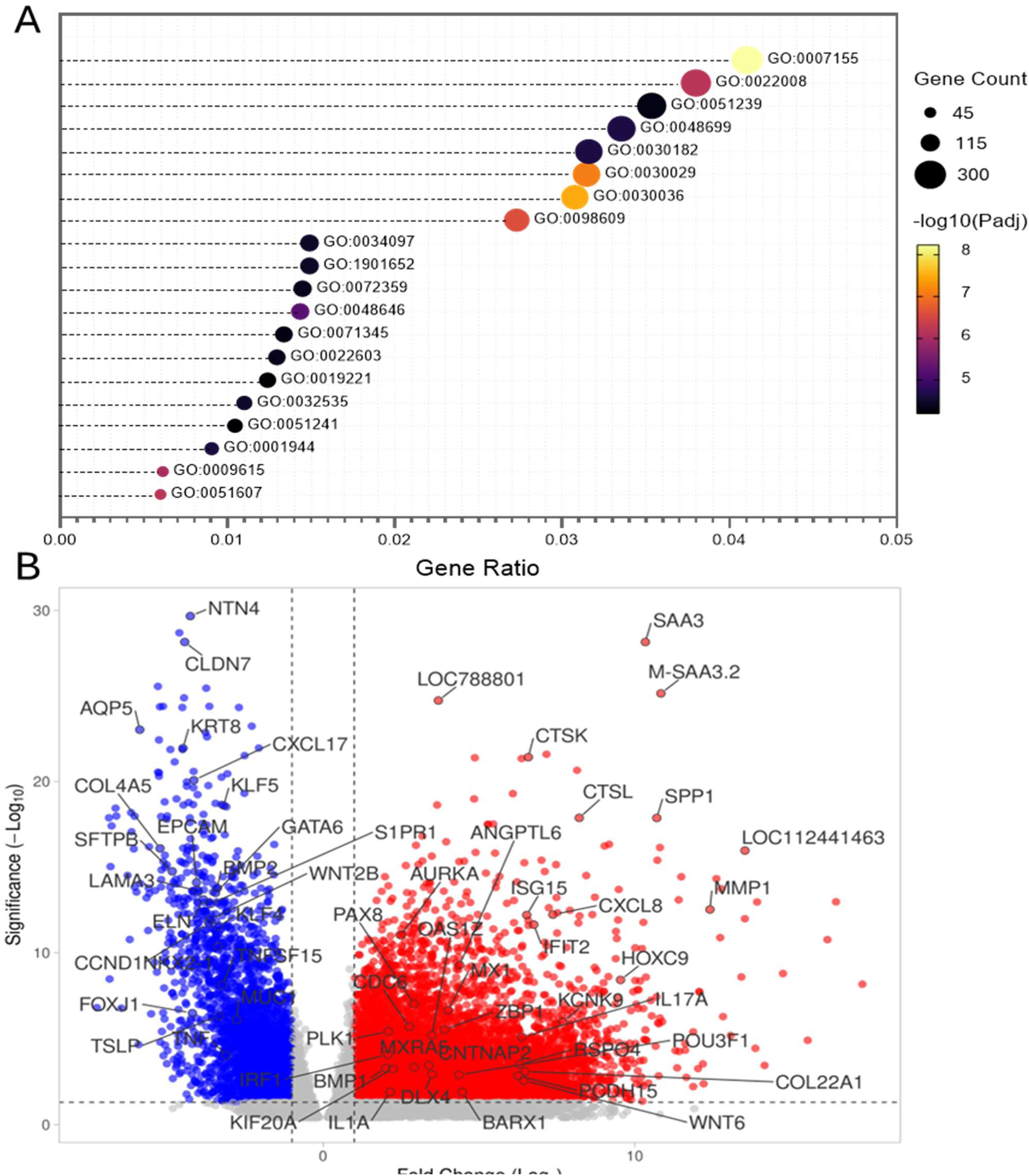
Transcriptional landscape of sequestra and necrotic tissues. (**A**) Top 20 enriched GO terms and (**B**) volcano plot highlighting differentially expressed genes with red marking significantly upregulated genes and blue marking significantly downregulated genes.

**Figure 8.**
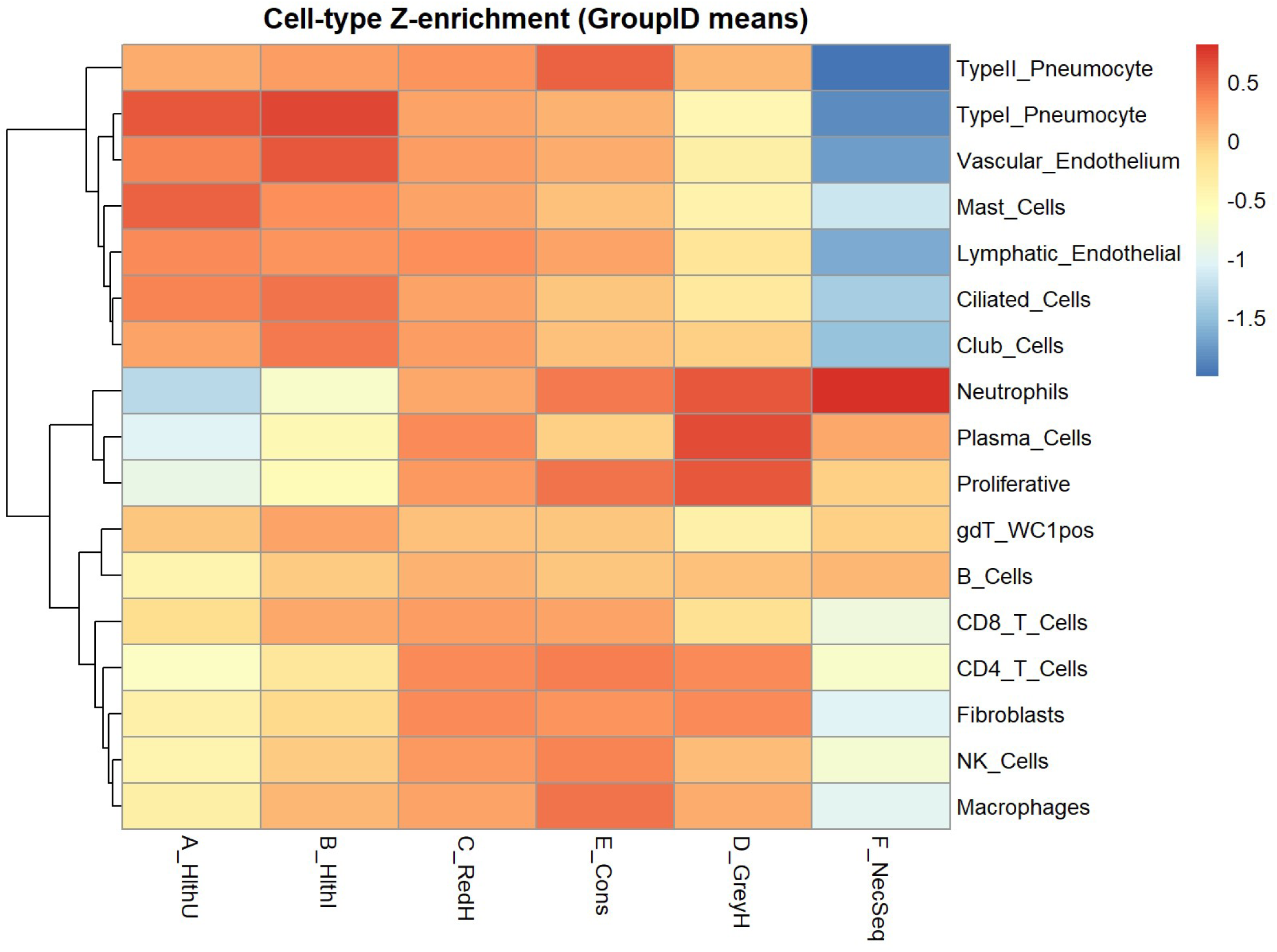
Z-score Signature Enrichment across CBPP lesion stages. The heatmap shows Z-score normalized enrichment values for curated bovine lung cell-type gene signatures (Macrophages, B cells, Neutrophils, Plasma cells, Dendritic Cells, Fibroblasts, T cells/ILCs, NK cells, Endothelial cells, Epithelial cells, etc).

On the other hand, downregulated transcripts indicated a profound loss of epithelial cell identity, with transcripts encoding hallmark airway and alveolar structural components such as surfactants, secretoglobins, keratins, epithelial adhesion factors (*SFTPA1/B/C/D, SCGB1A1/B1D/B3A2/B2A2,AQP5 EPCAM, KRT7/8/18/19, AGER, S1PR1*) and tight-junction constituents (*CLDN1/3/4/5/7/18* and *OCLN*) being reduced compared to healthy tissue controls (**120–126**). Downregulation of *NKX2-1, FOXA1, GATA6* and *HOPX*, and epithelial cell renewal transcripts *CCND1*, *KLF4/5*, *TEAD1/3/4* and *YAP1* further indicates collapse of the transcriptional network that maintains lung epithelial lineage identity **(G2-G4**). This was further supported by downregulation in developmental genes involved in Wnt, BMP and Notch signaling (*DKK2/3, WNT2B/7A/11, BMP2/3/4/6, NOTCH3* and *JAG2*) (**146,148**). Genes associated with ciliary assembly and motility were downregulated, including *FOXJ1, DNAH5/11, DNAL1/4, CFAP43/47/52, CFAP2SS, IFT57*, and *TEKT1/3* (**172**). The downregulation of *MUC1/15* and *PIGR* further support the collapse of the mucosal barrier function (**130**). Multiple collagen IV isoforms (*COL4A1/2/3/4/5/6*), laminins (*LAMA3/5, LAMB2/3*), laminin related netrin (*NTN4*) and other matrix associated proteins (*ELN, FBLN5, HMCN1, PRELP, SPON2, ITIH5, CLIC5*) were downregulated, indicating degradation of the epithelial basement membrane and consistent with the necrotic appearance of the tissue (**55,87–89, 132**). This was accompanied by the downregulation of angiogenic and endothelial genes *CD34*, *CDH5*, *TEK*, *TIE1*, *ESAM*, *RASIP1*, *GJA5*, *PLVAP*, *AQP1* and *MMRN2*, which indicates loss of microvascular endothelium within necrotic cores and aligns with the avascular nature of sequestra and the replacement of lung tissue with fibrotic encapsulation (**173**). Furthermore, metabolic and transport pathways were also broadly suppressed with xenobiotic and lipid-metabolizing enzymes (CYP1A1/2A13/2B6/2F1/4B1, AOX1/2. FMO2/3/5, UGT1A1/1A6, UGT2A1), and solute carriers (SLC6A14/6A20/16A11/22A31/45A1) being downregulated (**72, 126**). Furthermore, several transcripts relating to epithelial-driven immune response (*CXCL17, IL17RB, IL20RB, CD83, CD209, KLRK1, TNFSF15, TNF, ILDR1/2* and *TSLP*) were downregulated, reflecting the collapse of epithelial derived cytokine networks that is likely resultant from epithelial cell death (**53, 92**).

Surprisingly, despite the obliterated normal tissue architecture in necrotic lesions, we observed persistent, though attenuated, *interferon-mediated signaling* and *response to virus pathways* (**GO:0140888**, **GO:0009615**), suggesting persistence of low level innate immune activation without the robust inflammatory response seen in earlier stages. ISG transcripts such as *ISG15, IFIT2/3, IFI6/30, IFIH1, OAS1X/Y/Z, OAS2, RSAD2, MX1, MX2, USP18, ZBP1, IRF1/4/7* and *IFI44/44L* were upregulated, indicating a persistent interferon imprinting (**21–24**). Chemokine and cytokine related genes such as *CXCL2/5/8, CCL3/8*, *IL1A/B*, *IL22/24/26/17A/17F/21/21R/23R/7R*) were also upregulated (**53**). This indirectly indicates that *Mmm* continues to persist in late-stage lesions, and has the ability to survive the overwhelming inflammatory host response. Other immune related pathways indicated largely regulatory processes, rather than effector-driven, with *negative regulation of viral processes*, *negative regulation of cytokine responses*, and *modulation of leukocyte proliferation and activation* (**GO: 0048525, GO0032101, GO:0045321**)GO terms being enriched.

Collectively, the transcriptional profile mirrors the pathological phenotype of necrotic tissue and depicts these lesions as terminally remodeled, non-viable tissue compartments in which epithelial identity, barrier integrity, ciliary and metabolic function have collapsed, while fibrotic structural remodeling and persistent interferon imprinting dominate the surviving cellular landscape. The significant suppression of airway and alveolar functional programs (such as surfactant production, mucociliary clearance, tight junction architecture), underscores the complete loss of functional parenchyma that is visually apparent in the affected lung tissue. Concurrently, the induction of matrix-modifying enzymes, angiogenic modulators, and morphogenesis-linked transcripts reflects a microenvironment focused on encapsulation, containment, and long-term stabilization of necrotic tissue rather than regeneration. This polarized transcriptional state supports sequestra as chronic, structurally isolated lesions maintained by stromal and immune programs that prioritize physical isolation of damaged/necrotic tissue over eventual restoration of normal lung architecture.

### Cell-type deconvolution analysis supports a coordinated shift from structural integrity to innate, adaptive, and chronic inflammatory programs across CBPP stages

Differential gene expression and gene ontology analysis above indicate that CBPP lung lesions result from an evolving inflammatory landscape, however traditional bulk RNA sequencing cannot distinguish whether these transcriptional programs arise from transcriptional programming within stable cell populations or from changes in cellular composition. To resolve this ambiguity, and assess whether the inferred biological processes correspond to shifts in the underlying cellular landscape, we performed cell-type deconvolution using curated cell specific gene signatures obtained from the Cattle Cell Atlas (CattleCA) (**174**). Cell type deconvolution is a computational technique that aims to infer the composition of cell populations within a heterogenous tissue using bulk RNAseq data (**175**). By refining cell specific transcripts using single cell RNAseq data from a bovine host, we hoped that this approach would allow for adequate estimation of the evolving cellular landscape in CBPP lung lesions.

Our analysis revealed a stage-dependent reorganization of the cellular landscape during CBPP progression that was supportive of the inferences made from differential gene expression and gene ontology analysis above. Healthy uninfected lungs were dominated by epithelial and endothelial signatures, consistent with intact alveolar and vascular architecture (**Figure 7**). These signatures declined steadily as the lesions progressed toward the Necrotic/Sequestra lesion stage, further indicating cell death and loss of functional and structural integrity of the lung due to progressing infection. As the lesion progressed toward Red Hepatization, we see enrichment of signatures associated with myeloid cells such as neutrophils and monocytes/macrophages consistent with recruitment of innate effectors that are characteristic of this pathological stage. Much like previous analysis indicated, Grey Hepatization was dominated by macrophage and fibroblast signatures, indicating profibrotic processes that culminate in the replacement of damaged lung tissue into fibrinous scar tissue that is a hallmark of CBPP lesions. Lastly, an enrichment of T cell and B cell signatures in late-stage lesions is indicative of the formation of germinal centers within developing tertiary lymphoid structures in the lung, which are a prominent histopathological characteristic of chronic Mycoplasma infection. The ongoing enrichment of neutrophil signatures throughout lesion maturation further indicates continued pathogen persistence in infected lungs, indicating the host response is not productive, and is unable to clear *Mmm* infection.

## DISCUSSION

Our study provides a lesion-stage-resolved transcriptomic framework for understanding the progression of lung lesions during Contagious Bovine Pleuropneumonia (CBPP). Our data reveal how *Mycoplasma mycoides* subsp. *mycoides* infection drives a coordinated sequence of inflammatory injury, immune modulation, and structural remodeling in the bovine lung that culminates in characteristic lung pathology. By assessing differential gene expression across lesions of differential maturity, we obtain a temporal look at the host response to infection, and how the inflammatory landscape in the lung evolves from the initial moments of colonization and pathogen recognition down to tissue repair following pathologic sequelae. We show that each lesion stage is defined by a molecular program that aligns closely with its pathological phenotype and reflects the hosts shifting priorities as the disease progresses (**Figure 9**). However, transcriptional programs associated with pathogen recognition, interferon stimulated pathways, and innate immune responses persist throughout lesion progression, indirectly indicating chronic persistence of *Mmm* in infected tissue, despite an overwhelming inflammatory response. This further substantiates the idea that infection-induced host responses to mycoplasma infections are maladaptive, serving to contribute to immunopathology without contributing to bacterial clearance.

**Figure 9.**
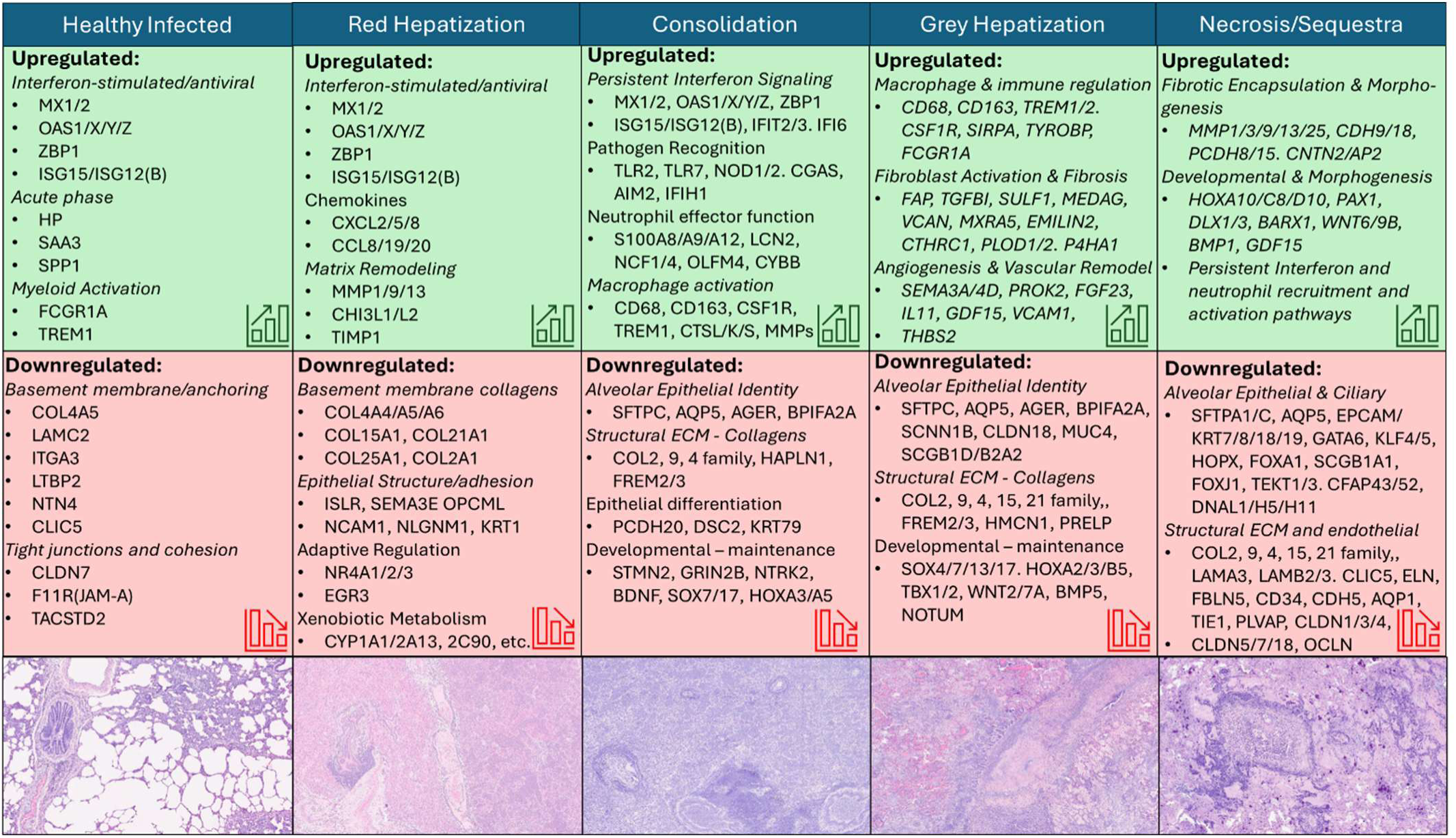
Summary of key findings per lesion stage and histopathological appearance of represented lung lesions.

Early lesions (Healthy Infected, Red Hepatization, and to an extent Consolidation) were dominated by a strong induction of innate immune pathways, including interferon stimulated gene expression, neutrophil recruitment, and inflammatory cytokines. These signatures reflect an intense but ultimately ineffective attempt to control *Mmm*, consistent with the chronic nature of disease. Concurrent upregulation of early extracellular matrix remodeling pathways aligns with the gross pathological characteristics of CBPP. Grey hepatization represented a transitional state in which inflammatory programs remained active but were increasingly accompanied by macrophage-and-fibroblast-mediated profibrotic processes, extracellular matrix deposition and early tissue remodeling. However, continued pathogen persistence was hinted by the retained antiviral response signatures and neutrophil-associated transcripts. Unlike earlier stages, where adaptive immune responses were repressed, grey hepatization revealed the upregulation of transcriptional pathways that point toward mobilization of T and B cell responses, and initiation of adaptive immunity. The concurrent persistence of pathways involved in extracellular matrix destruction with reparative morphogenetic and angiogenetic pathways indicate healing attempts with the goal of preserving tissue function. However continued pathogen persistence likely does not allow for functional tissue repair, thereby shifting the host response toward terminal tissue necrosis and sequestrum formation. The transcriptional landscape of Necrotic/Sequestra revealed a profound loss of epithelial identity of lung tissue, including suppression of surfactant genes, mucociliary machinery, epithelial tight-junctions, and epithelial lineage-defining transcription factors. These changes reflect the destruction and functional loss of airway and alveolar epithelium within the necrotic cores that form from cell death. Profibrotic processes were upregulated, driving remodeling processes associated with non-functional scar tissue formation rather than functional tissue repair. In an effort to contain infection and isolate necrotic tissue, this mature lesion reflected a transcriptional response focused on encapsulation and sequestra formation which is characteristic of chronic CBPP.

Across all stages, the prominence of unannotated LOC- transcripts, many of which were strongly differentially expressed, highlights a substantial gap in bovine genome annotation and indicates that key regulatory elements involved in lung injury and chronic remodeling remain uncharacterized. Their consistent expression patterns indicate that *Mmm* infection activates transcriptional programs that extend beyond currently annotated pathways, highlighting the need for further research in the area, as our limited knowledge about the function of these gene products limits our interpretation of the data we present herein.

Despite these limitations, beyond defining the transcriptomic progression of CBPP disease, our work provides several important contributions to the field. First, it establishes the first comprehensive stage-resolved transcriptomic atlas of CBPP to date, linking classical pathological findings with their underlying molecular programs. This framework offers a reference point for future studies seeking to understand how *Mmm* may manipulate host immunity and tissue architecture over time. While the idea that immunopathology plays a role in disease due to mycoplasma infections is not new in the field, our work significantly substantiates the hypothesis. Second, our data highlight an unexpected feature of the host response, which is the prominence of antiviral and interferon-linked pathways that serve as the dominant response to a bacterial agent, underscoring the need to revisit long held assumptions about the nature of mycoplasma infections. It is unclear whether the upregulation of these antiviral immune responses indicates a significant intracellular niche of persistence for *Mmm*, or if the pathogen is stimulating a maladaptive immune response as a means of immune evasion to promote chronic colonization and persistence in a highly inflammatory environment. More mechanistic studies are needed to address these questions, as their answers may be essential for informing the development of efficacious therapeutic or prophylactic methods to help control the disease. Furthermore, by providing a global look at the host response to *Mmm* infection, our data provide opportunity for biomarker discovery that can inform diagnostic development.

Altogether, these contributions position our work as a resource for the CBPP community and as a conceptual advance for understanding chronic respiratory mycoplasmoses. The molecular roadmap presented here provides a platform for hypothesis generation, comparative studies, and translational efforts aimed at mitigating CBPP burden.

## MATERIALS AND METHODS

### Study design and samples

Fourteen Small East African Zebu cattle were divided into two groups and challenged via nebulization with sterile growth medium (n=4) to serve as uninfected controls while others (n=10) were challenged with *Mycoplasma mycoides subsp*. *mycoides* strain *Afade*. Animals were necropsied when humane endpoint criteria were reached or at 6 weeks post-infection at the end of the study. Animals were humanely euthanized via injection of Sodium Pentobarbitone (200mg/mL) at a dosage of 100mg/kg of body weight, and death was confirmed by a veterinarian. Lung tissue samples (n = 81) were collected from necropsied cattle representing six pathological conditions: A_HlthU/Healthy Uninfected (Healthy tissue from uninfected animals), B_HlthI/Healthy Infected (Healthy looking tissue from infected animals), C_RedH/Red Hepatization), D_GreyH/Grey Hepatization), E_Cons/Consolidation and F_NecSeq/Necrotic tissue and sequestra). Pathological samples were identified by consensus of two independent, study-blind pathologists experienced with CBPP pathology. Sample metadata are provided in **Supplementary Table S1**. All animal procedures were performed under the approved institutional animal care and use (IACUC) protocol (ILRI-IACUC2024-23/1).

### RNA extraction, sequencing, and preprocessing

Collected tissues were immediately placed into TRIzol at the time of collection and frozen at -80C until all tissues from all experimental animals were collected. In preparation for RNA extraction, samples were defrosted at room temperature, tissues were minced with scissors, then placed back into the original TRIZol aliquot and refrozen overnight. The next day, samples were thawed at room temperature and RNA was extracted using a modified phenol/chloroform extraction protocol from the recommended Invitrogen TRIzol Reagent User Guide. Briefly, the TRIzol supernatant was transferred to a new sterile tube and 200uL of both chloroform and tissue culture grade water was added. Samples were vigorously mixed then incubated at room temperature for 3 minutes. Samples were centrifuged at 12000xg for 15 minutes to separate the phases. The upper, colorless aqueous phase was collected and transferred to a new tube, and the chloroform step was repeated to wash the fraction. The RNA was precipitated by adding 500uL of isopropanol, mixing vigorously, then incubating at room temperature for 10 minutes. The RNA was pelleted via centrifugation at 12000xg for 10 minutes. The pellet was resuspended in 1mL 75% ethanol, vortexed briefly, and centrifuged for 5 minutes at 7500xg. This step was repeated twice to sufficiently wash the RNA pellet, which was then air-dried for 7 minutes to evaporate the remaining ethanol. The pellets were resuspended in 50uL of tissue culture grade water and incubated in a heat block at 60C for 15 minutes. RNA concentrations and quality were measured using NanoDrop. Dual rRNA depleted RNA libraries were prepared using the Illumina Stranded Total RNA Prep with Ligation, Ribo-Zero Plus Microbiome kit following manufacturer recommended instructions, and samples were sequenced using the P4 NextSeq2000 P4 cartridges targeting 100M paired-end reads of 100-150bp in length per sample.

### Data Analyses

Following trimming of adapters and low-quality bases the raw sequence data were mapped to the *Bos taurus* reference genome to generate gene count matrices. While Small East African Zebus are a *Bos indicus* species, we observed no significant differences in alignment rates between *Bos taurus* and *Bos indicus*, and given the better annotation of the *Bos taurus* genome, decided to move forward with the *Bos taurus* alignment. As an additional quality control step, samples with unique alignment of less that 50% were removed from analysis. Differential gene expression analysis was conducted using DESeq2 version 1.50.2 using R version 4.5.2. Gene Ontology Enrichment Analysis was performed using clusterProfiler version 4.18.4 (**176–179**). Cell-type deconvolution was performed using a z-score–based approach applied to variance-stabilized transcript (VST) expression values (see **Supplementary File** for methods and R-code). Volcano plots and heatmaps were generated in R using built in plotting functions while GO Term enrichment bubble plots were generated in GraphPad Prism version 9.0.0.

## Acknowledgments

We would like to acknowledge Dalton Bioanalytics for initial sample QC, mapping to *Bos taurus* genome and GO analyses. This work was supported by funds made available by the Center of Excellence for Vaccine Research and the USDA-ARS Non-Assistance Cooperative Agreement #5030-32000-236-002S.

## Author Contributions

Conceptualization (*A.B.M., R.G.O., E.R.T., S.M.S., S.J.G.*)

Methodology (*A.B.M, R.G.O., E.R.T., S.M.S., S.J.G.*)

Software (*A.B.M.*)

Formal Analysis (*A.B.M.*)

Validation (*A.B.M., E.S., E.R.T., S.M.S., S.J.G.*)

Investigation (*A.B.M, A.M., R.G.O., M.S., H.W., M.A., N.O.O., W.C., M.L.H., J.M.M., E.S.*)

Resources (*S.J.G., E.S.*)

Data Curation (*A.B.M, A.M.*)

Writing – Original Draft (*A.B.M*)

Writing – Review C Editing (*A.B.M, A.M., R.G.O., M.S., H.W., M.A., N.O.O., W.C., M.L.H., J.M.M., E.R.T., S.M.S., E.S., S.J.G.*)

Visualization (*A.B.M.*)

Supervision (*E.S., S.J.G.*)

Project Administration (*A.B.M., A.M., E.R.T., S.M.S., E.S., S.J.G.*)

Funding Acquisition (*S.J.G., S.M.S., E.R.T*.)

