## Supplemental Information for "Host transcriptional programs underlying lesion development in contagious bovine pleuropneumonia"

**Author Affiliations:**

**Author Contributions:**

Conceptualization (*A.B.M., R.G.O., E.R.T., S.M.S., S.J.G.*)
Methodology (*A.B.M, R.G.O., E.R.T., S.M.S., S.J.G.*)
Software (*A.B.M.*)
Formal Analysis (*A.B.M.*)
Validation (*A.B.M., E.S., E.R.T., S.M.S., S.J.G.*)
Investigation (*A.B.M, A.M., R.G.O., M.S., H.W., M.A., N.O.O., W.C., M.L.H., J.M.M., E.S.*)
Resources (*S.J.G., E.S.*)
Data Curation (*A.B.M, A.M.*)
Writing – Original Draft (*A.B.M*)
Writing – Review & Editing (*A.B.M, A.M., R.G.O., M.S., H.W., M.A., N.O.O., W.C., M.L.H., J.M.M., E.R.T., S.M.S., E.S., S.J.G.*)
Vizualization (*A.B.M.*)
Supervision (*E.S., S.J.G.*)
Project Administration (*A.B.M., A.M., E.R.T., S.M.S., E.S., S.J.G.*)
Funding Acquisition (*S.J.G., S.M.S., E.R.T.*)

**Supplementary Methods: Cell-type Deconvolution**

Raw gene-level count matrices and metadata were imported into R (v4.5.2). Count matrix columns were aligned to metadata by exact matching of SampleID, and both objects were subset and reordered to ensure one-to-one correspondence. All downstream analyses were performed on this aligned dataset. Cell-type marker genes were curated from the Cattle Cell Atlas (CattleCA) for a signature list delineating: Ciliated Cells, Type I Pneumocytes, Type II Pneumocytes, Mast Cells, Plasma Cells, Proliferative Cells, Club Cells, Lymphatic and Vascular Endothelium, NK Cells, Fibroblasts, γδ T cells (WC1+), B Cells, Macrophages, CD4 T cells, CD8 T cells, and neutrophils. For each signature, only genes present in the dataset were retained. To stabilize variance across the dynamic range of expression values, raw counts were transformed using $\log_2$(count+1). For each gene, a Z-score was computed across all samples. Genes with zero variance were assigned a value of 1 to avoid undefined values. For each cell-type signature, enrichment scores were calculated as the mean Z-score of all genes belonging to that signature within each sample. To evaluate cell-type trends across disease progression, enrichment scores were aggregated by GroupID.

*R Code Used:*

```
# 0. Setup and directories

library(pheatmap)
count_file    <- "count_matrix_filtered.csv"
metadata_file <- "metadata_filtered.csv"

out_dir <- "Z_en_scRNA_Tcsplit"
if (!dir.exists(out_dir)) dir.create(out_dir, recursive = TRUE)

# 1. Load count matrix & metadata

metadata <- read.csv(metadata_file, header = TRUE, stringsAsFactors = FALSE)
counts_raw <- read.csv(count_file, header = TRUE, stringsAsFactors = FALSE, check.names = FALSE)

# Assume first column = gene IDs
rownames(counts_raw) <- counts_raw[[1]]
counts_raw <- counts_raw[, -1, drop = FALSE]

# Align samples
common_samples <- intersect(colnames(counts_raw), metadata$SampleID)
counts <- counts_raw[, common_samples, drop = FALSE]
metadata_sub <- metadata[match(common_samples, metadata$SampleID), ]
rownames(metadata_sub) <- metadata_sub$SampleID

saveRDS(counts,       file = file.path(out_dir, "counts_aligned.rds"))
saveRDS(metadata_sub, file = file.path(out_dir, "metadata_aligned.rds"))
```

### 2. Define scRNA signatures

```
celltype_signatures_raw <- list(
  Ciliated_Cells = c("CFAP299","PHGR1","PIGR","SEC14L3","TUBA1D","CETN4",
                "LRRIQ1","RARRES1","CAPS","FAM183A","TPPP3","FOXJ1","CCDC78"),
  TypeI_Pneumocyte = c("AGER","CYP4B1","GSTA2","GSTT2","CYP2B6","HSD17B13",
                "LMO7","SPADH1","KRT7","CLDN18","ADIRF","CLIC5","ICAM1"),
  Mast_Cells = c("PTI","TPSB2","LTC4S","VIPR2","FCER1A","GZMB","KIT"),
  TypeII_Pneumocyte = c("SFTPC","SFTPA1","WFDC18","S100G","SFTPB","SLC34A2",
                "SCD","SFTPD","NPC2","SFTA2","ABCA3","MUC1"),
  Plasma_Cells = c("TXNDC5","JCHAIN","MZB1"),
  Proliferative = c("TMSB10","TOP2A","CENPF","MKI67","PCLAF","UBE2C",
              "SPC24","TPX2","STMN1","BIRC5","PTTG1"),
  Club_Cells = c("SCGB1A1","SCGB2A2","SCGB1D","SCGB3A2","CLDN4","BPIFA1",
            "KRT15","BPIFB1","TFF3","AOX2"),
  Lymphatic_Endothelial = c("LYVE1","CCL21","MMRN1","PTN","RELN","S100A1",
                  "PKHD1L1","CCN3","PKDCC","TNFAIP8L3","APOLD1","PDPN"),
  Vascular_Endothelium = c("CYYR1","CALCRL","CCDC85A","SMAD6","MYZAP","LDB2",
                  "HMCN1","PRKG1","RAMP3","PRICKLE2","PECAM1","KDR","FLT1"),
  NK_Cells = c("NKG2A","DNAJC6","IDO1","KLRB1","TXK","NCR1","CX3CR1","FHIP1A","KLRK1"),
  Fibroblasts = c("DCN","COL1A1","MGP","COL3A1","COL1A2","PI16","C1S","C1R",
             "FBLN1","CFH","PDGFRA","FAP","TGFBI","SULF1","MEDAG"),
  gdT_WC1pos = c("WC1.3","RHEX","BLK","PLCL1","CD163L1","SLC16A10"),
  B_Cells = c("MS4A1","PAX5","TNFRSF13C","SOX5","BLNK","MYO1E","BANK1",
          "BCL11A","FCRL1","EBF1","MA4A1/CD19","CD21","CD79B","CD79A"),
  Macrophages = c("SGMS2","CD5L","MARCO","C1QB","C1QA","C1QC","LYZ","CD36",
             "CD68","HMOX1","MS4A7","MRC1","MSR1","CD74","CD14","CD16","FCGR3A"),
  CD4_T_Cells = c("CTLA4","CD28","LEF1","CD4","ICOS","CAMK4","SLC37A3",
             "INPP4B","MAF","CD5","IL7R","CD3E"),
  CD8_T_Cells = c("CTSW","CRTAM","CD8B","GZMM","CD96","CCL5","PRF1","THEMIS",
             "NKG7","GNLY","CD8A","CD3G","CD3D","CSF1R"),
  Neutrophils = c("HSD11B1","SDS","MEFV","GPR84","TGM3","RUBCNL","TCN1",
             "CXCR1","CXCL8","CSF3R","S100A9","S100A8","S100A12","IL1RN")
)

# Harmonize case
rownames(counts) <- toupper(rownames(counts))
celltype_signatures <- lapply(celltype_signatures_raw, toupper)

# Intersect with available genes
celltype_signatures_filtered <- lapply(celltype_signatures, function(g)
  intersect(g, rownames(counts))
)

signature_sizes <- sapply(celltype_signatures_filtered, length)
write.csv(data.frame(Signature = names(signature_sizes),
                N_genes = signature_sizes),
```

```
          file = file.path(out_dir, "signature_sizes.csv"),
          row.names = FALSE)

saveRDS(celltype_signatures_filtered,
        file = file.path(out_dir, "celltype_signatures_filtered.rds"))
```

**# 3. Compute per-gene Z-scores**

```
counts_mat <- as.matrix(counts)
mode(counts_mat) <- "numeric"

log_counts <- log2(counts_mat + 1)

gene_means <- rowMeans(log_counts)
gene_sds   <- apply(log_counts, 1, sd)
gene_sds[gene_sds == 0] <- 1

z_mat <- sweep(log_counts, 1, gene_means, "-")
z_mat <- sweep(z_mat, 1, gene_sds, "/")

saveRDS(z_mat, file = file.path(out_dir, "z_scores_gene_by_sample.rds"))
```

**# 4. Signature enrichment (mean Z)**

```
signature_names <- names(celltype_signatures_filtered)

sig_by_sample <- matrix(NA_real_,
                        nrow = length(signature_names),
                        ncol = ncol(z_mat),
                        dimnames = list(signature_names, colnames(z_mat)))

for (sig in signature_names) {
  g <- celltype_signatures_filtered[[sig]]
  if (length(g) > 0)
    sig_by_sample[sig, ] <- colMeans(z_mat[g, , drop = FALSE])
}

sig_by_sample_df <- as.data.frame(sig_by_sample)

write.csv(sig_by_sample_df,
          file = file.path(out_dir, "signature_enrichment_by_sample.csv"))
saveRDS(sig_by_sample_df,
        file = file.path(out_dir, "signature_enrichment_by_sample.rds"))
```

**# 5. GroupID aggregation**

```
group_order <- c("A_HlthU","B_HlthI","C_RedH","E_Cons","D_GreyH","F_NecSeq")
```

```
group_vec <- metadata_sub$GroupID
names(group_vec) <- metadata_sub$SampleID

sig_by_group <- matrix(NA_real_,
                       nrow = nrow(sig_by_sample_df),
                       ncol = length(group_order),
                       dimnames = list(rownames(sig_by_sample_df), group_order))

for (g in group_order) {
  samples <- names(group_vec)[group_vec == g]
  if (length(samples) > 0)
    sig_by_group[, g] <- rowMeans(sig_by_sample_df[, samples, drop = FALSE])
}

sig_by_group_df <- as.data.frame(sig_by_group)

write.csv(sig_by_group_df,
          file = file.path(out_dir, "signature_enrichment_by_GroupID.csv"))
saveRDS(sig_by_group_df,
        file = file.path(out_dir, "signature_enrichment_by_GroupID.rds"))
```

**# 6. Heatmaps**

```
# Sample-level ordering
sample_order <- order(factor(metadata_sub$GroupID, levels = group_order),
                      metadata_sub$Time,
                      metadata_sub$AnimalID)

sig_by_sample_ord <- sig_by_sample_df[, sample_order, drop = FALSE]
metadata_ord      <- metadata_sub[sample_order, ]

annotation_col <- data.frame(
  GroupID = factor(metadata_ord$GroupID, levels = group_order),
  Time    = metadata_ord$Time,
  Batch   = metadata_ord$Sequencing_Batch
)
rownames(annotation_col) <- metadata_ord$SampleID

# Sample-level heatmap
pdf(file.path(out_dir, "heatmap_signature_by_sample.pdf"), width = 10, height = 6)
pheatmap(sig_by_sample_ord,
         cluster_rows = TRUE,
         cluster_cols = FALSE,
         annotation_col = annotation_col,
         show_colnames = FALSE,
         main = "Cell-type Z-enrichment (Sample-level)")
```

```
dev.off()

# Group-level heatmap
pdf(file.path(out_dir, "heatmap_signature_by_GroupID.pdf"), width = 6, height = 6)
pheatmap(sig_by_group_df[, group_order, drop = FALSE],
         cluster_rows = TRUE,
         cluster_cols = FALSE,
         main = "Cell-type Z-enrichment (GroupID means)")
dev.off()
```

**Supplementary Table S1.** Sample metadata identifying information on the animal from which the sample was collected , infection status, pathological condition, and sequencing run.

| *SampleID* | *GroupID* | *Pathological_Condition* | *Infection_Status* | *AnimalID* | *Sequencing_Batch* |
|---|---|---|---|---|---|
| *Tissue_1_S1* | A_HlthU | Healthy_Section_A | Uninfected | BV075 | ThirdRun |
| *Tissue_2_S2* | A_HlthU | Healthy_Section_A | Uninfected | BV075 | ThirdRun |
| *Tissue_3_S1* | A_HlthU | Healthy_Section_A | Uninfected | BV084 | FirstRun |
| *Tissue_4_S3* | A_HlthU | Healthy_Section_A | Uninfected | BV084 | ThirdRun |
| *Tissue_5_S4* | A_HlthU | Healthy_Section_A | Uninfected | BV084 | ThirdRun |
| *Tissue_6_S5* | A_HlthU | Healthy_Section_A | Uninfected | BV090 | ThirdRun |
| *Tissue_8_S7* | A_HlthU | Healthy_Section_A | Uninfected | BV064 | ThirdRun |
| *Tissue_9_S2* | A_HlthU | Healthy_Section_A | Uninfected | BV064 | FirstRun |
| *Tissue_10_S3* | A_HlthU | Healthy_Section_A | Uninfected | BV075 | FirstRun |
| *Tissue_11_S8* | A_HlthU | Healthy_Section_A | Uninfected | BV090 | ThirdRun |
| *Tissue_12_S4* | A_HlthU | Healthy_Section_A | Uninfected | BV090 | FirstRun |
| *Tissue_15_S7* | B_HlthI | Healthy_Section_B | Infected | BV062 | FirstRun |
| *Tissue_16_S8* | B_HlthI | Healthy_Section_B | Infected | BV074 | FirstRun |
| *Tissue_17_S9* | B_HlthI | Healthy_Section_B | Infected | BV088 | FirstRun |
| *Tissue_18_S10* | B_HlthI | Healthy_Section_B | Infected | BV088 | FirstRun |
| *Tissue_19_S11* | B_HlthI | Healthy_Section_B | Infected | BV088 | FirstRun |
| *Tissue_20_S12* | B_HlthI | Healthy_Section_B | Infected | BV095 | FirstRun |
| *Tissue_21_S13* | B_HlthI | Healthy_Section_B | Infected | BV095 | FirstRun |
| *Tissue_22_S14* | B_HlthI | Healthy_Section_B | Infected | BV099 | FirstRun |
| *Tissue_23_S15* | C_RedH | Red Hepatization | Infected | BV062 | FirstRun |
| *Tissue_24_S16* | C_RedH | Red Hepatization | Infected | BV062 | FirstRun |
| *Tissue_25_S17* | C_RedH | Red Hepatization | Infected | BV062 | FirstRun |
| *Tissue_26_S18* | C_RedH | Red Hepatization | Infected | BV074 | FirstRun |
| *Tissue_27_S19* | C_RedH | Red Hepatization | Infected | BV074 | FirstRun |
| *Tissue_28_S20* | C_RedH | Red Hepatization | Infected | BV074 | FirstRun |
| *Tissue_29_S21* | C_RedH | Red Hepatization | Infected | BV087 | FirstRun |
| *Tissue_30_S22* | C_RedH | Red Hepatization | Infected | BV087 | FirstRun |
| *Tissue_31_S23* | C_RedH | Red Hepatization | Infected | BV087 | FirstRun |
| *Tissue_32_S24* | C_RedH | Red Hepatization | Infected | BV088 | FirstRun |
| *Tissue_33_S25* | C_RedH | Red Hepatization | Infected | BV088 | FirstRun |
| *Tissue_34_S26* | C_RedH | Red Hepatization | Infected | BV088 | FirstRun |
| *Tissue_35_S27* | C_RedH | Red Hepatization | Infected | BV088 | FirstRun |
| *Tissue_36_S9* | C_RedH | Red Hepatization | Infected | BV095 | ThirdRun |
| *Tissue_37_S28* | C_RedH | Red Hepatization | Infected | BV095 | FirstRun |
| *Tissue_38_S29* | C_RedH | Red Hepatization | Infected | BV095 | FirstRun |
| *Tissue_39_S30* | C_RedH | Red Hepatization | Infected | BV099 | FirstRun |
| *Tissue_40_S10* | C_RedH | Red Hepatization | Infected | BV099 | ThirdRun |
| *Tissue_41_S1* | C_RedH | Red Hepatization | Infected | BV099 | SecondRun |
| *Tissue_42_S2* | D_GreyH | Grey Hepatization | Infected | BV062 | SecondRun |
| *Tissue_43_S3* | D_GreyH | Grey Hepatization | Infected | BV074 | SecondRun |
| *Tissue_44_S4* | D_GreyH | Grey Hepatization | Infected | BV087 | SecondRun |
| *Tissue_45_S5* | D_GreyH | Grey Hepatization | Infected | BV087 | SecondRun |
| *Tissue_46_S6* | D_GreyH | Grey Hepatization | Infected | BV087 | SecondRun |
| *Tissue_47_S7* | D_GreyH | Grey Hepatization | Infected | BV088 | SecondRun |
| *Tissue_48_S8* | D_GreyH | Grey Hepatization | Infected | BV095 | SecondRun |
| *Tissue_49_S9* | D_GreyH | Grey Hepatization | Infected | BV095 | SecondRun |
| *Tissue_50_S10* | D_GreyH | Grey Hepatization | Infected | BV095 | SecondRun |
| *Tissue_51_S11* | D_GreyH | Grey Hepatization | Infected | BV099 | SecondRun |
| *Tissue_52_S12* | D_GreyH | Grey Hepatization | Infected | BV099 | SecondRun |
| *Tissue_54_S14* | E_Cons | Consolidation | Infected | BV074 | SecondRun |
| *Tissue_55_S15* | E_Cons | Consolidation | Infected | BV088 | SecondRun |
| *Tissue_56_S16* | E_Cons | Consolidation | Infected | BV088 | SecondRun |
| *Tissue_57_S17* | E_Cons | Consolidation | Infected | BV088 | SecondRun |
| *Tissue_58_S18* | E_Cons | Consolidation | Infected | BV088 | SecondRun |
| *Tissue_59_S19* | E_Cons | Consolidation | Infected | BV088 | SecondRun |
| *Tissue_60_S20* | E_Cons | Consolidation | Infected | BV088 | SecondRun |
| *Tissue_61_S21* | E_Cons | Consolidation | Infected | BV088 | SecondRun |
| *Tissue_64_S24* | B_HlthI | Healthy_Section_B | Infected | BV065 | SecondRun |
| *Tissue_65_S25* | B_HlthI | Healthy_Section_B | Infected | BV069 | SecondRun |
| *Tissue_66_S26* | B_HlthI | Healthy_Section_B | Infected | BV069 | SecondRun |
| *Tissue_67_S27* | B_HlthI | Healthy_Section_B | Infected | BV070 | SecondRun |
| *Tissue_68_S28* | B_HlthI | Healthy_Section_B | Infected | BV072 | SecondRun |
| *Tissue_69_S29* | B_HlthI | Healthy_Section_B | Infected | BV072 | SecondRun |
| *Tissue_70_S30* | B_HlthI | Healthy_Section_B | Infected | BV091 | SecondRun |
| *Tissue_71_S11* | B_HlthI | Healthy_Section_B | Infected | BV065 | ThirdRun |

| | | | | | |
|---|---|---|---|---|---|
| *Tissue_72_S12* | B_Hlthl | Healthy_Section_B | Infected | BV091 | ThirdRun |
| *Tissue_73_S13* | B_Hlthl | Healthy_Section_B | Infected | BV091 | ThirdRun |
| *Tissue_74_S14* | C_RedH | Red Hepatization | Infected | BV072 | ThirdRun |
| *Tissue_75_S15* | C_RedH | Red Hepatization | Infected | BV072 | ThirdRun |
| *Tissue_76_S16* | C_RedH | Red Hepatization | Infected | BV091 | ThirdRun |
| *Tissue_77_S17* | D_GreyH | Grey Hepatization | Infected | BV091 | ThirdRun |
| *Tissue_78_S18* | F_NecSeq | Necrotic-Sequestra | Infected | BV072 | ThirdRun |
| *Tissue_79_S19* | F_NecSeq | Necrotic-Sequestra | Infected | BV072 | ThirdRun |
| *Tissue_80_S20* | F_NecSeq | Necrotic-Sequestra | Infected | BV065 | ThirdRun |
| *Tissue_81_S21* | F_NecSeq | Necrotic-Sequestra | Infected | BV065 | ThirdRun |
| *Tissue_82_S22* | F_NecSeq | Necrotic-Sequestra | Infected | BV065 | ThirdRun |
| *Tissue_83_S23* | F_NecSeq | Necrotic-Sequestra | Infected | BV072 | ThirdRun |
| *Tissue_84_S24* | F_NecSeq | Necrotic-Sequestra | Infected | BV072 | ThirdRun |
| *Tissue_85_S25* | F_NecSeq | Necrotic-Sequestra | Infected | BV072 | ThirdRun |
| *Tissue_86_S26* | F_NecSeq | Necrotic-Sequestra | Infected | BV091 | ThirdRun |

**Supplementary Figure S1.** All sample Raw Read Count distribution indicating % unique alignment to the *Bos taurus* genome.

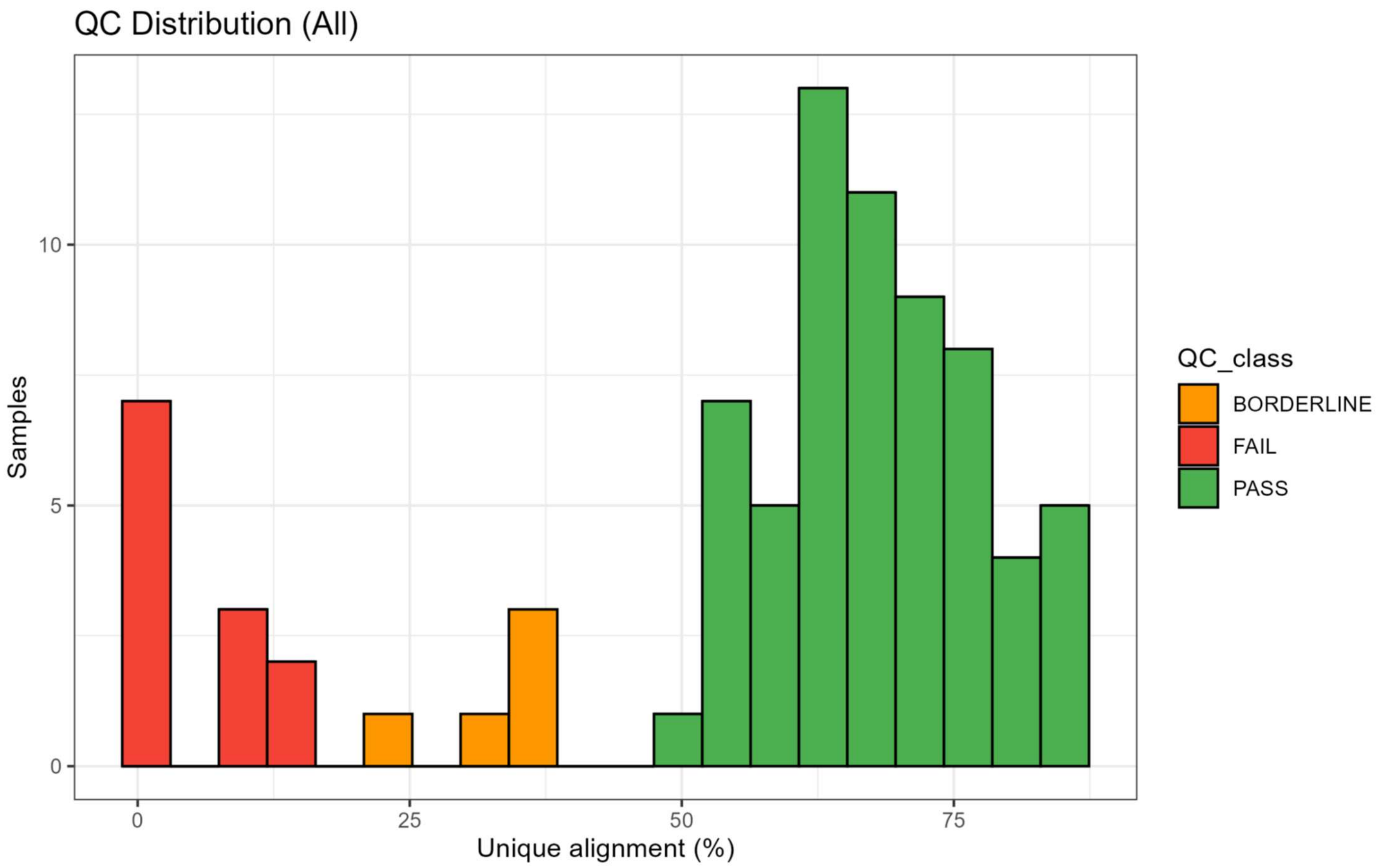

**Supplementary Figure S2.** QC filtered sample Read Count distribution indicating % unique alignment to the *Bos taurus* genome separated out by pathological condition.

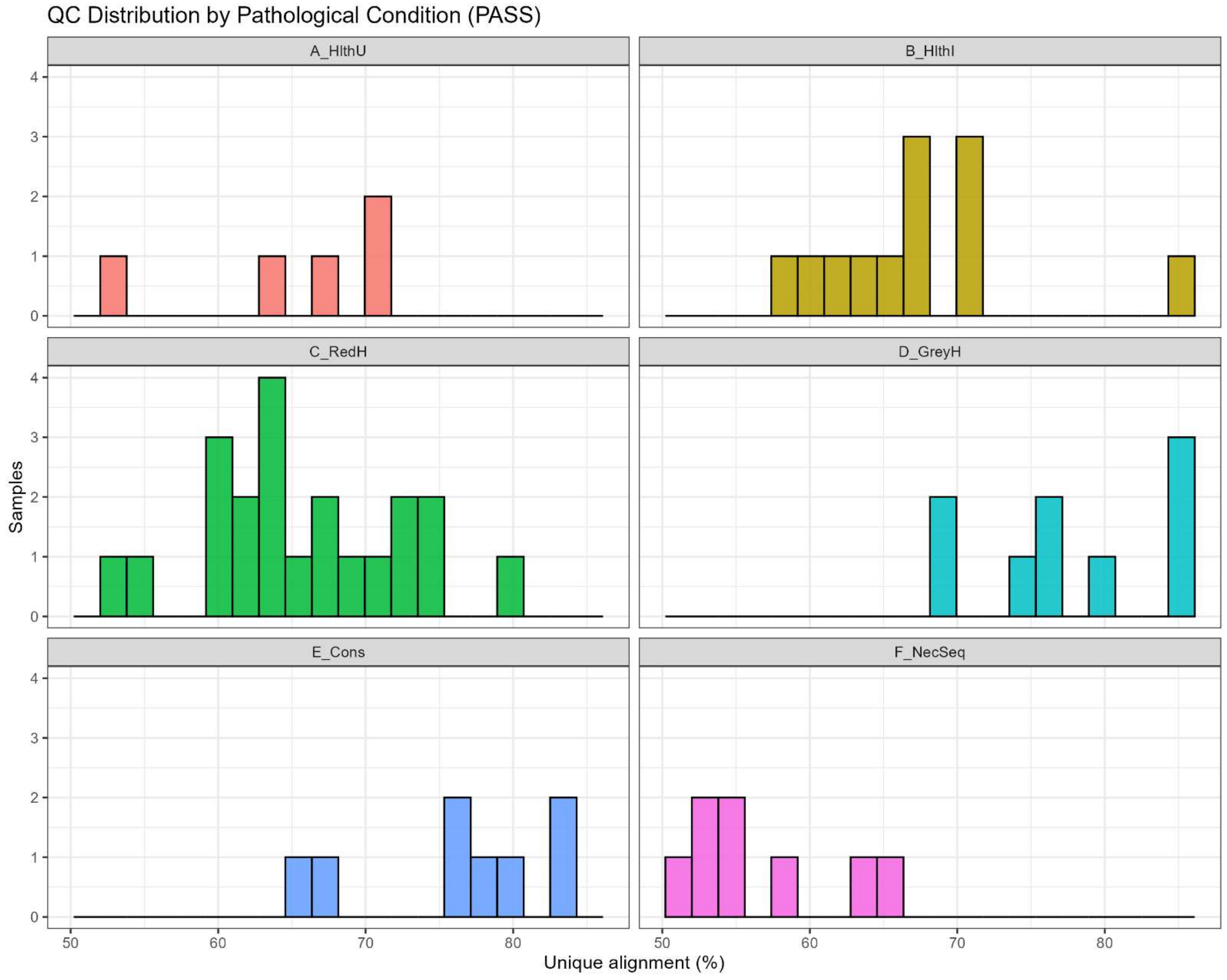

**Supplementary Figure S3.** Principal Component Analysis indicating sample distribution by infection status (top) and pathological condition (bottom).

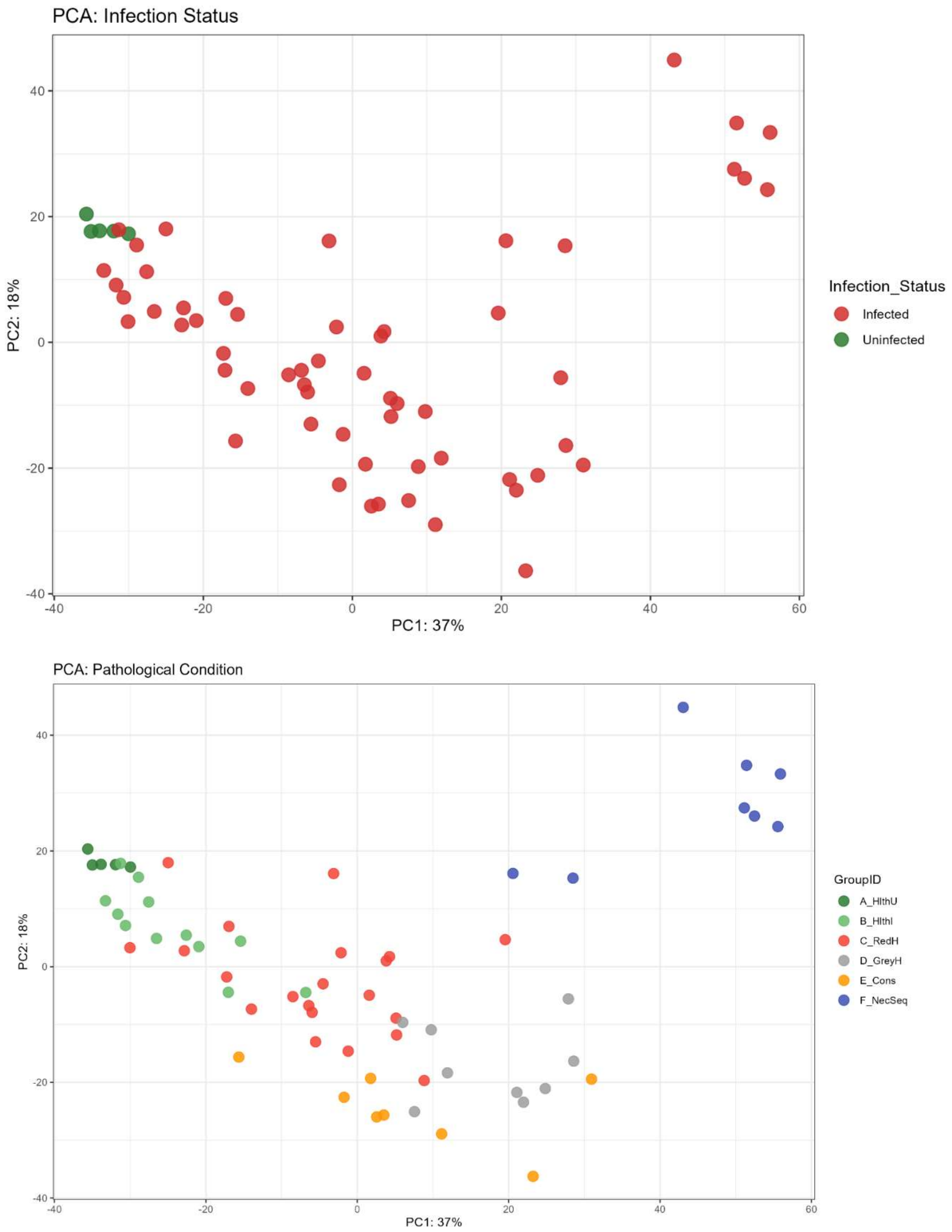

**Supplementary Figure S4.** Sample-to-Sample Distance Heatmap

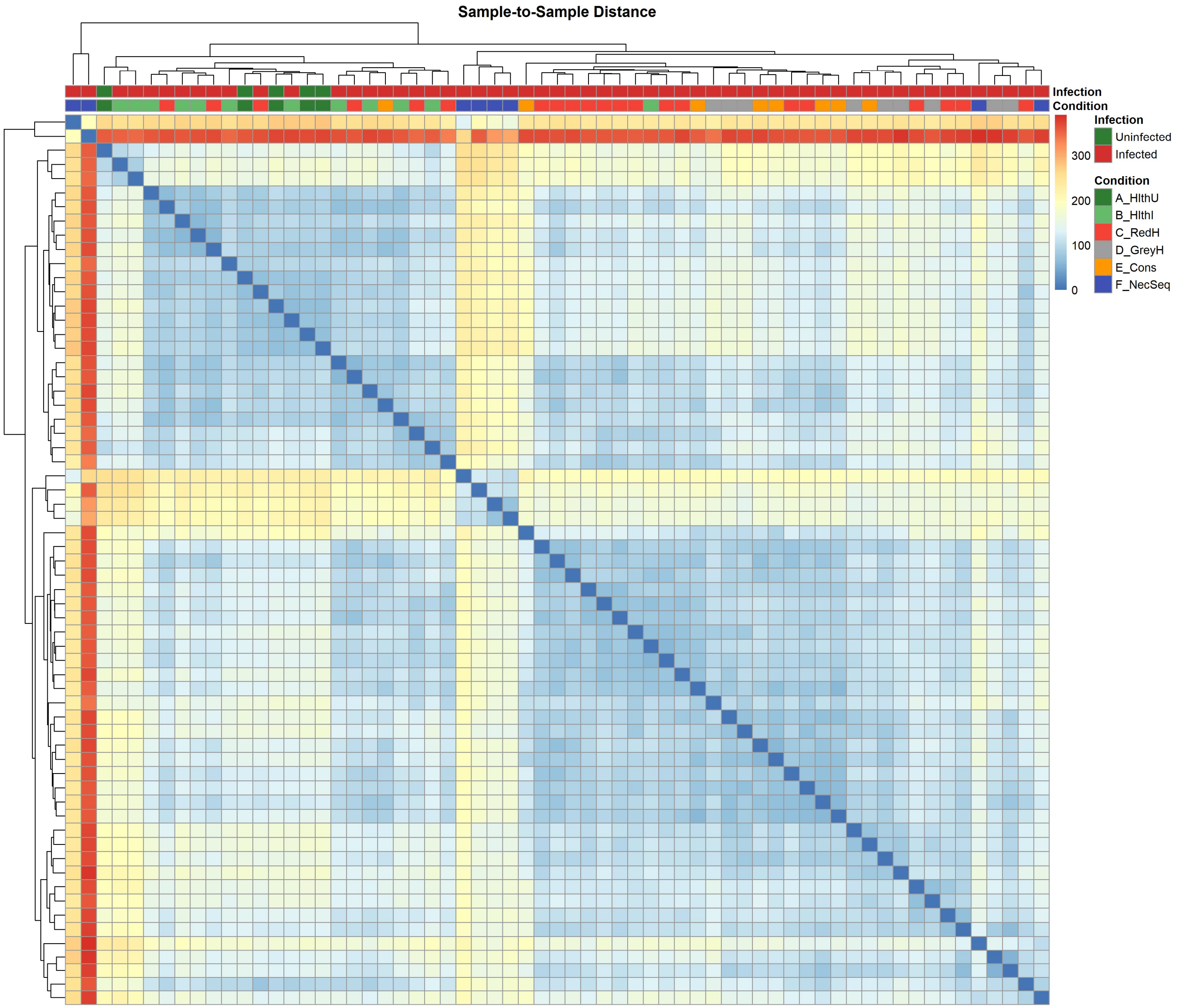

**Supplementary Table S2**. *Enriched GO Terms for Healthy Infected versus Healthy Uninfected Comparisons*.

| *ID* | *Description* | *GeneRatio* | *p.adjust* |
|---|---|---|---|
| *GO:0051607* | defense response to virus | 21/143 | 8.33981E-28 |
| *GO:0009615* | response to virus | 21/143 | 1.1719E-27 |
| *GO:0044419* | biological process involved in interspecies interaction between organisms | 31/143 | 3.25801E-19 |
| *GO:0043207* | response to external biotic stimulus | 30/143 | 7.36874E-19 |
| *GO:0051707* | response to other organism | 30/143 | 7.36874E-19 |
| *GO:0009607* | response to biotic stimulus | 30/143 | 7.36874E-19 |
| *GO:0045069* | regulation of viral genome replication | 11/143 | 1.38554E-16 |
| *GO:0045071* | negative regulation of viral genome replication | 11/143 | 1.38554E-16 |
| *GO:0048525* | negative regulation of viral process | 11/143 | 2.71627E-16 |
| *GO:0019079* | viral genome replication | 11/143 | 5.09513E-16 |
| *GO:1903900* | regulation of viral life cycle | 11/143 | 6.7023E-14 |
| *GO:0019058* | viral life cycle | 12/143 | 9.39566E-14 |
| *GO:0050792* | regulation of viral process | 11/143 | 1.9642E-13 |
| *GO:0016032* | viral process | 12/143 | 4.59937E-12 |
| *GO:0140888* | interferon-mediated signaling pathway | 7/143 | 2.2727E-09 |
| *GO:0098542* | defense response to other organism | 16/143 | 1.81324E-08 |
| *GO:0060337* | type I interferon-mediated signaling pathway | 6/143 | 5.88557E-08 |
| *GO:0071357* | cellular response to type I interferon | 6/143 | 5.88557E-08 |
| *GO:0034340* | response to type I interferon | 6/143 | 9.22522E-08 |
| *GO:0140546* | defense response to symbiont | 15/143 | 1.08848E-07 |
| *GO:0034341* | response to type II interferon | 6/143 | 1.32572E-07 |
| *GO:0045087* | innate immune response | 14/143 | 2.36069E-07 |
| *GO:0035456* | response to interferon-beta | 5/143 | 8.04066E-07 |
| *GO:0046597* | host-mediated suppression of symbiont invasion | 5/143 | 8.04066E-07 |
| *GO:0051851* | host-mediated perturbation of symbiont process | 5/143 | 1.31381E-06 |
| *GO:0042130* | negative regulation of T cell proliferation | 5/143 | 3.14785E-06 |
| *GO:0035821* | modulation of process of another organism | 5/143 | 4.51444E-06 |
| *GO:0032945* | negative regulation of mononuclear cell proliferation | 5/143 | 7.07414E-06 |
| *GO:0050672* | negative regulation of lymphocyte proliferation | 5/143 | 7.07414E-06 |
| *GO:0070664* | negative regulation of leukocyte proliferation | 5/143 | 7.07414E-06 |
| *GO:0042098* | T cell proliferation | 6/143 | 7.07414E-06 |
| *GO:0042129* | regulation of T cell proliferation | 6/143 | 7.07414E-06 |
| *GO:0008285* | negative regulation of cell population proliferation | 7/143 | 7.07414E-06 |
| *GO:0034097* | response to cytokine | 11/143 | 7.07414E-06 |
| *GO:1901652* | response to peptide | 11/143 | 7.07414E-06 |
| *GO:0019221* | cytokine-mediated signaling pathway | 10/143 | 8.02724E-06 |
| *GO:0071345* | cellular response to cytokine stimulus | 10/143 | 1.72033E-05 |
| *GO:0002682* | regulation of immune system process | 15/143 | 2.12386E-05 |
| *GO:0032944* | regulation of mononuclear cell proliferation | 6/143 | 4.24354E-05 |
| *GO:0050670* | regulation of lymphocyte proliferation | 6/143 | 4.24354E-05 |
| *GO:0070663* | regulation of leukocyte proliferation | 6/143 | 4.24354E-05 |
| *GO:0050868* | negative regulation of T cell activation | 5/143 | 4.66871E-05 |
| *GO:1903038* | negative regulation of leukocyte cell-cell adhesion | 5/143 | 4.66871E-05 |
| *GO:0032943* | mononuclear cell proliferation | 6/143 | 6.33237E-05 |
| *GO:0046651* | lymphocyte proliferation | 6/143 | 6.33237E-05 |
| *GO:0070661* | leukocyte proliferation | 6/143 | 6.33237E-05 |
| *GO:0042110* | T cell activation | 7/143 | 6.33237E-05 |
| *GO:0022408* | negative regulation of cell-cell adhesion | 5/143 | 7.57238E-05 |
| *GO:0051250* | negative regulation of lymphocyte activation | 5/143 | 0.000106032 |
| *GO:0050863* | regulation of T cell activation | 6/143 | 0.000108817 |
| *GO:0002695* | negative regulation of leukocyte activation | 5/143 | 0.000189597 |
| *GO:0050866* | negative regulation of cell activation | 5/143 | 0.000189597 |
| *GO:0007162* | negative regulation of cell adhesion | 5/143 | 0.000285229 |
| *GO:0032309* | icosanoid secretion | 4/143 | 0.000314865 |
| *GO:0050482* | arachidonate secretion | 4/143 | 0.000314865 |
| *GO:1903963* | arachidonate transport | 4/143 | 0.000314865 |
| *GO:0002683* | negative regulation of immune system process | 6/143 | 0.000352274 |
| *GO:0001775* | cell activation | 9/143 | 0.00040335 |
| *GO:0002694* | regulation of leukocyte activation | 7/143 | 0.000406834 |
| *GO:0071715* | icosanoid transport | 4/143 | 0.000447548 |
| *GO:0050865* | regulation of cell activation | 7/143 | 0.000508487 |
| *GO:1903037* | regulation of leukocyte cell-cell adhesion | 5/143 | 0.000645498 |
| *GO:0045321* | leukocyte activation | 8/143 | 0.000778729 |
| *GO:0051249* | regulation of lymphocyte activation | 6/143 | 0.001175348 |
| *GO:0007159* | leukocyte cell-cell adhesion | 5/143 | 0.001494776 |

| | | | |
|---|---|---|---|
| *GO:0046649* | lymphocyte activation | 7/143 | 0.001608389 |
| *GO:0022407* | regulation of cell-cell adhesion | 5/143 | 0.001879425 |
| *GO:0015909* | long-chain fatty acid transport | 4/143 | 0.002199281 |
| *GO:0052547* | regulation of peptidase activity | 5/143 | 0.003653752 |
| *GO:0032496* | response to lipopolysaccharide | 4/143 | 0.00420803 |
| *GO:0042127* | regulation of cell population proliferation | 9/143 | 0.005176789 |
| *GO:0051241* | negative regulation of multicellular organismal process | 6/143 | 0.005215816 |
| *GO:0002237* | response to molecule of bacterial origin | 4/143 | 0.005422903 |
| *GO:0008283* | cell population proliferation | 9/143 | 0.007721213 |
| *GO:0030162* | regulation of proteolysis | 6/143 | 0.011590389 |
| *GO:0015908* | fatty acid transport | 4/143 | 0.012000958 |
| *GO:0006869* | lipid transport | 7/143 | 0.01415157 |
| *GO:0030155* | regulation of cell adhesion | 5/143 | 0.015523121 |
| *GO:0010876* | lipid localization | 7/143 | 0.018680887 |
| *GO:0010466* | negative regulation of peptidase activity | 4/143 | 0.019501774 |
| *GO:0050776* | regulation of immune response | 8/143 | 0.019501774 |
| *GO:0002252* | immune effector process | 6/143 | 0.020685542 |
| *GO:0009617* | response to bacterium | 6/143 | 0.02187603 |
| *GO:0006915* | apoptotic process | 10/143 | 0.022544334 |
| *GO:0051239* | regulation of multicellular organismal process | 11/143 | 0.023085756 |
| *GO:0046942* | carboxylic acid transport | 6/143 | 0.023334218 |
| *GO:0015849* | organic acid transport | 6/143 | 0.023835247 |
| *GO:0008219* | cell death | 10/143 | 0.024705376 |
| *GO:0012501* | programmed cell death | 10/143 | 0.024705376 |
| *GO:0071222* | cellular response to lipopolysaccharide | 3/143 | 0.026174042 |
| *GO:0045861* | negative regulation of proteolysis | 4/143 | 0.026174042 |
| *GO:0046634* | regulation of alpha-beta T cell activation | 2/143 | 0.030562963 |
| *GO:0006879* | intracellular iron ion homeostasis | 3/143 | 0.030562963 |
| *GO:0071219* | cellular response to molecule of bacterial origin | 3/143 | 0.032793219 |
| *GO:0050778* | positive regulation of immune response | 7/143 | 0.032993569 |
| *GO:0001959* | regulation of cytokine-mediated signaling pathway | 2/143 | 0.034538069 |
| *GO:0003254* | regulation of membrane depolarization | 2/143 | 0.034538069 |
| *GO:0060759* | regulation of response to cytokine stimulus | 2/143 | 0.034538069 |
| *GO:0045089* | positive regulation of innate immune response | 4/143 | 0.034538069 |
| *GO:0006644* | phospholipid metabolic process | 7/143 | 0.034538069 |

**Supplementary Table S3.** *Relevant Differentially Expressed Genes for Healthy Infected versus Healthy Uninfected Comparisons.*

| ***Gene*** | ***Change*** | ***log2 F.C.*** | ***p.adjusted*** |
|---|---|---|---|
| *SAA3* | Increased | 6.005 | 1.72312E-13 |
| *CYM* | Decreased | -10.018 | 4.99042E-09 |
| *M-SAA3.2* | Increased | 5.43 | 1.06564E-08 |
| *NTN4* | Decreased | -1.912 | 2.33537E-08 |
| *PLA2G2D4* | Increased | 3.592 | 1.4319E-06 |
| *HP* | Increased | 3.553 | 4.24554E-05 |
| *ZBP1* | Increased | 3.48 | 4.24554E-05 |
| *OAS1Z* | Increased | 3.107 | 9.42156E-05 |
| *ISG15* | Increased | 3.655 | 0.000168436 |
| *MX1* | Increased | 3.035 | 0.000206667 |
| *OAS1Y* | Increased | 2.853 | 0.000206667 |
| *IFI6* | Increased | 2.787 | 0.000255557 |
| *NR4A2* | Decreased | -1.628 | 0.000255557 |
| *PLA2G5* | Increased | 4.713 | 0.000379031 |
| *CILP2* | Decreased | -4.15 | 0.000865537 |
| *OAS1X* | Increased | 2.415 | 0.001484307 |
| *APOBEC3Z1* | Increased | 3.823 | 0.001718113 |
| *CLEC4F* | Increased | 4.461 | 0.002005364 |
| *LGALS9* | Increased | 1.757 | 0.002418581 |
| *IFIT2* | Increased | 3.268 | 0.003240784 |
| *TIMP1* | Increased | 3.004 | 0.003444879 |
| *SOCS2* | Decreased | -1.118 | 0.003444879 |
| *MX2* | Increased | 2.729 | 0.003525088 |
| *RSAD2* | Increased | 2.538 | 0.020735251 |

**Supplementary Table S4**. *Enriched GO Terms for Red Hepatization versus Healthy Uninfected Comparisons.*

| *ID* | *Description* | *GeneRatio* | *p.adjust* |
|---|---|---|---|
| *GO:0009615* | response to virus | 41/3787 | 1.24219E-13 |
| *GO:0051607* | defense response to virus | 39/3787 | 7.06803E-13 |
| *GO:0002682* | regulation of immune system process | 147/3787 | 6.5432E-10 |
| *GO:0044419* | biological process involved in interspecies interaction between organisms | 147/3787 | 1.32117E-09 |
| *GO:0031347* | regulation of defense response | 68/3787 | 3.38543E-09 |
| *GO:0034097* | response to cytokine | 79/3787 | 3.38543E-09 |
| *GO:1901652* | response to peptide | 79/3787 | 3.38543E-09 |
| *GO:0080134* | regulation of response to stress | 91/3787 | 4.96049E-09 |
| *GO:0006954* | inflammatory response | 70/3787 | 1.06552E-08 |
| *GO:0032101* | regulation of response to external stimulus | 85/3787 | 1.06552E-08 |
| *GO:0045321* | leukocyte activation | 71/3787 | 2.07105E-08 |
| *GO:0001775* | cell activation | 80/3787 | 2.07105E-08 |
| *GO:0002684* | positive regulation of immune system process | 117/3787 | 2.07105E-08 |
| *GO:0009607* | response to biotic stimulus | 138/3787 | 2.07105E-08 |
| *GO:0043207* | response to external biotic stimulus | 137/3787 | 2.07105E-08 |
| *GO:0051707* | response to other organism | 137/3787 | 2.07105E-08 |
| *GO:0046649* | lymphocyte activation | 61/3787 | 8.30733E-08 |
| *GO:1903900* | regulation of viral life cycle | 23/3787 | 1.46369E-07 |
| *GO:0071345* | cellular response to cytokine stimulus | 68/3787 | 1.46369E-07 |
| *GO:0098542* | defense response to other organism | 97/3787 | 1.98905E-07 |
| *GO:0007155* | cell adhesion | 173/3787 | 2.19796E-07 |
| *GO:0045087* | innate immune response | 88/3787 | 4.51215E-07 |
| *GO:0140546* | defense response to symbiont | 94/3787 | 9.44504E-07 |
| *GO:0051249* | regulation of lymphocyte activation | 43/3787 | 1.31367E-06 |
| *GO:0002831* | regulation of response to biotic stimulus | 45/3787 | 1.61312E-06 |
| *GO:0034340* | response to type I interferon | 14/3787 | 1.69797E-06 |
| *GO:0050792* | regulation of viral process | 23/3787 | 2.03215E-06 |
| *GO:0050865* | regulation of cell activation | 50/3787 | 2.03215E-06 |
| *GO:0019221* | cytokine-mediated signaling pathway | 61/3787 | 2.03215E-06 |
| *GO:0098609* | cell-cell adhesion | 118/3787 | 2.03215E-06 |
| *GO:0044093* | positive regulation of molecular function | 111/3787 | 2.46925E-06 |
| *GO:0045069* | regulation of viral genome replication | 16/3787 | 2.88496E-06 |
| *GO:0045071* | negative regulation of viral genome replication | 16/3787 | 2.88496E-06 |
| *GO:0032944* | regulation of mononuclear cell proliferation | 27/3787 | 3.02256E-06 |
| *GO:0050670* | regulation of lymphocyte proliferation | 27/3787 | 3.02256E-06 |
| *GO:0070663* | regulation of leukocyte proliferation | 27/3787 | 3.02256E-06 |
| *GO:0045088* | regulation of innate immune response | 42/3787 | 3.05422E-06 |
| *GO:0002694* | regulation of leukocyte activation | 48/3787 | 3.18323E-06 |
| *GO:0060337* | type I interferon-mediated signaling pathway | 13/3787 | 4.39475E-06 |
| *GO:0071357* | cellular response to type I interferon | 13/3787 | 4.39475E-06 |
| *GO:0140888* | interferon-mediated signaling pathway | 14/3787 | 6.72967E-06 |
| *GO:0019058* | viral life cycle | 26/3787 | 6.89663E-06 |
| *GO:0032943* | mononuclear cell proliferation | 28/3787 | 6.89663E-06 |
| *GO:0046651* | lymphocyte proliferation | 28/3787 | 6.89663E-06 |
| *GO:0070661* | leukocyte proliferation | 28/3787 | 6.89663E-06 |
| *GO:0048525* | negative regulation of viral process | 16/3787 | 8.07015E-06 |
| *GO:0002252* | immune effector process | 63/3787 | 8.21965E-06 |
| *GO:0002833* | positive regulation of response to biotic stimulus | 39/3787 | 1.03577E-05 |
| *GO:0050776* | regulation of immune response | 95/3787 | 1.35586E-05 |
| *GO:0032103* | positive regulation of response to external stimulus | 49/3787 | 1.63603E-05 |
| *GO:1902531* | regulation of intracellular signal transduction | 147/3787 | 1.70094E-05 |
| *GO:0007159* | leukocyte cell-cell adhesion | 30/3787 | 1.81499E-05 |
| *GO:0045089* | positive regulation of innate immune response | 37/3787 | 1.86958E-05 |
| *GO:0051239* | regulation of multicellular organismal process | 152/3787 | 1.97153E-05 |
| *GO:0019079* | viral genome replication | 16/3787 | 2.21139E-05 |
| *GO:0031349* | positive regulation of defense response | 40/3787 | 2.24899E-05 |
| *GO:0050790* | regulation of catalytic activity | 149/3787 | 3.21827E-05 |
| *GO:0030036* | actin cytoskeleton organization | 124/3787 | 3.65667E-05 |
| *GO:0042110* | T cell activation | 36/3787 | 3.67377E-05 |
| *GO:0050778* | positive regulation of immune response | 84/3787 | 4.44938E-05 |
| *GO:0030029* | actin filament-based process | 127/3787 | 5.24665E-05 |
| *GO:0002218* | activation of innate immune response | 32/3787 | 5.7127E-05 |
| *GO:0022407* | regulation of cell-cell adhesion | 30/3787 | 6.98827E-05 |

| | | | |
|---|---|---|---|
| GO:0002263 | cell activation involved in immune response | 23/3787 | 8.28719E-05 |
| GO:0051241 | negative regulation of multicellular organismal process | 48/3787 | 9.22042E-05 |
| GO:0051250 | negative regulation of lymphocyte activation | 19/3787 | 0.000140202 |
| GO:0002221 | pattern recognition receptor signaling pathway | 23/3787 | 0.000144809 |
| GO:0002366 | leukocyte activation involved in immune response | 22/3787 | 0.000198213 |
| GO:0043408 | regulation of MAPK cascade | 54/3787 | 0.000227485 |
| GO:1903706 | regulation of hemopoiesis | 19/3787 | 0.000266072 |
| GO:0048585 | negative regulation of response to stimulus | 114/3787 | 0.000283372 |
| GO:0002695 | negative regulation of leukocyte activation | 20/3787 | 0.000349668 |
| GO:0050866 | negative regulation of cell activation | 20/3787 | 0.000349668 |
| GO:0000165 | MAPK cascade | 59/3787 | 0.000349668 |
| GO:1903037 | regulation of leukocyte cell-cell adhesion | 24/3787 | 0.000463304 |
| GO:0002253 | activation of immune response | 65/3787 | 0.000463304 |
| GO:0002683 | negative regulation of immune system process | 31/3787 | 0.000541837 |
| GO:0030097 | hemopoiesis | 40/3787 | 0.000569976 |
| GO:0043085 | positive regulation of catalytic activity | 80/3787 | 0.000728807 |
| GO:0030155 | regulation of cell adhesion | 40/3787 | 0.000755416 |
| GO:0050863 | regulation of T cell activation | 26/3787 | 0.000861654 |
| GO:0002758 | innate immune response-activating signaling pathway | 28/3787 | 0.000971013 |
| GO:0042098 | T cell proliferation | 18/3787 | 0.001076651 |
| GO:0042129 | regulation of T cell proliferation | 18/3787 | 0.001076651 |
| GO:0016477 | cell migration | 117/3787 | 0.001076651 |
| GO:0050727 | regulation of inflammatory response | 25/3787 | 0.001138037 |
| GO:0051251 | positive regulation of lymphocyte activation | 25/3787 | 0.001138037 |
| GO:0044403 | biological process involved in symbiotic interaction | 12/3787 | 0.001424533 |
| GO:0002275 | myeloid cell activation involved in immune response | 11/3787 | 0.001699522 |
| GO:0016032 | viral process | 27/3787 | 0.001849519 |
| GO:0035456 | response to interferon-beta | 9/3787 | 0.001882249 |
| GO:0002696 | positive regulation of leukocyte activation | 29/3787 | 0.001882249 |
| GO:0050867 | positive regulation of cell activation | 29/3787 | 0.001882249 |
| GO:0050900 | leukocyte migration | 31/3787 | 0.001882249 |
| GO:0002250 | adaptive immune response | 32/3787 | 0.001882249 |
| GO:0009117 | nucleotide metabolic process | 70/3787 | 0.001882249 |
| GO:0050864 | regulation of B cell activation | 17/3787 | 0.00236608 |
| GO:0050868 | negative regulation of T cell activation | 15/3787 | 0.002470792 |
| GO:1903038 | negative regulation of leukocyte cell-cell adhesion | 15/3787 | 0.002470792 |
| GO:0051336 | regulation of hydrolase activity | 77/3787 | 0.002475696 |

**Supplementary Table S5.** *Relevant Differentially Expressed Genes for Red Hepatization versus Healthy Uninfected Comparisons.*

| *Gene* | *Change* | *log2 F.C.* | *p.adjusted* |
|---|---|---|---|
| *SAA3* | Increased | 7.936 | 3.24E-31 |
| *M-SAA3.2* | Increased | 7.944 | 3.86E-25 |
| *ISG12(B)* | Increased | 6.884 | 1.24E-16 |
| *OAS1Z* | Increased | 4.662 | 4.36E-16 |
| *NR4A2* | Decreased | -2.555 | 4.50E-16 |
| *SOD2* | Increased | 4.356 | 1.77E-15 |
| *ZBP1* | Increased | 4.904 | 2.55E-15 |
| *SPP1* | Increased | 7.286 | 6.10E-15 |
| *ISG15* | Increased | 5.361 | 7.11E-15 |
| *MX2* | Increased | 4.602 | 1.17E-13 |
| *IFIT2* | Increased | 5.43 | 1.17E-13 |
| *APOBEC3Z1* | Increased | 6.055 | 3.25E-13 |
| *MX1* | Increased | 4.213 | 3.70E-13 |
| *CD14* | Increased | 3.989 | 3.97E-13 |
| *TREM1* | Increased | 5.53 | 4.33E-13 |
| *CHI3L1* | Increased | 3.714 | 1.95E-12 |
| *OAS1Y* | Increased | 3.833 | 2.64E-12 |
| *ISG20* | Increased | 4.232 | 4.50E-12 |
| *OAS1X* | Increased | 3.462 | 2.38E-11 |
| *CLEC4F* | Increased | 6.541 | 2.57E-11 |
| *COL4A5* | Decreased | -3.207 | 2.88E-11 |
| *MMP9* | Increased | 7.619 | 4.00E-10 |
| *RSAD2* | Increased | 4.167 | 7.39E-10 |
| *CYM* | Decreased | -8.098 | 1.19E-09 |
| *PTX3* | Increased | 3.748 | 1.57E-09 |
| *CCL8* | Increased | 4.929 | 1.57E-09 |
| *COL25A1* | Decreased | -3.404 | 3.30E-09 |
| *MMP25* | Increased | 3.57 | 1.67E-08 |
| *NR4A1* | Decreased | -3.461 | 1.82E-08 |
| *COL4A6* | Decreased | -2.727 | 5.27E-08 |
| *CHI3L2* | Increased | 4.86 | 5.41E-08 |
| *NOS2* | Increased | 3.595 | 1.29E-07 |
| *SIGLEC1* | Increased | 4.377 | 2.68E-07 |
| *SEMA3E* | Decreased | -2.903 | 3.89E-07 |
| *NR4A3* | Decreased | -3.029 | 1.20E-06 |
| *TLR7* | Increased | 2.618 | 1.33E-06 |
| *IL27* | Increased | 3.249 | 1.44E-06 |
| *EGR3* | Decreased | -2.879 | 2.47E-06 |
| *COL4A4* | Decreased | -2.233 | 4.81E-06 |
| *TLR2* | Increased | 2.023 | 8.60E-06 |
| *OPCML* | Decreased | -3.603 | 1.08E-05 |
| *MMP13* | Increased | 3.849 | 2.54E-05 |
| *ISLR* | Decreased | -2.902 | 3.07E-05 |
| *CCL19* | Increased | 3.335 | 5.83E-05 |
| *MMP1* | Increased | 5.878 | 5.97E-05 |
| *IL19* | Increased | 6.054 | 6.03E-05 |
| *IL17A* | Increased | 4.98 | 7.98E-05 |
| *CXCL2* | Increased | 3.033 | 1.73E-04 |
| *CYP1A1* | Decreased | -4.02 | 1.74E-04 |
| *CYP2C90* | Decreased | -3.017 | 1.90E-04 |
| *COL21A1* | Decreased | -2.077 | 2.40E-04 |
| *COL15A1* | Decreased | -2.029 | 2.43E-04 |
| *CCL20* | Increased | 3.715 | 3.55E-04 |
| *CXCL5* | Increased | 2.19 | 3.80E-04 |
| *CYP2A13* | Decreased | -3.144 | 3.98E-04 |
| *SAA2* | Increased | 5.011 | 5.90E-04 |
| *KRT1* | Decreased | -3.668 | 7.16E-04 |
| *CXCL8* | Increased | 2.733 | 1.10E-03 |
| *IL21* | Increased | 2.749 | 2.74E-03 |
| *CYP3A24* | Decreased | -2.062 | 7.99E-03 |
| *COL2A1* | Decreased | -3.666 | 1.45E-02 |
| *NCAM1* | Decreased | -2.177 | 2.50E-02 |
| *NLGN1* | Decreased | -2.146 | 4.34E-02 |

| | | | |
|---|---|---|---|
| *PLA2G2D4* | Increased | 4.559 | 2.55E-15 |
| *PLA2G5* | Increased | 7.421 | 1.09E-14 |
| *NTN4* | Decreased | -2.345 | 3.43E-17 |
| *CILP2* | Decreased | -5.01 | 1.38E-08 |
| *TIMP1* | Increased | 4.82 | 1.32E-12 |
| *MMP15* | Decreased | -2.02 | 7.41E-06 |
| *MMP16* | Decreased | -3.08 | 1.91E-07 |

**Supplementary Table S6**. *Enriched GO Terms for Consolidation versus Healthy Uninfected Comparisons.*

| ***ID*** | ***Description*** | ***GeneRatio*** | ***p.adjust*** |
|---|---|---|---|
| *GO:0009615* | response to virus | 37/3240 | 4.06618E-12 |
| *GO:0051607* | defense response to virus | 36/3240 | 4.06618E-12 |
| *GO:0034097* | response to cytokine | 74/3240 | 4.38365E-10 |
| *GO:1901652* | response to peptide | 74/3240 | 4.38365E-10 |
| *GO:0044419* | biological process involved in interspecies interaction between organisms | 130/3240 | 4.53334E-09 |
| *GO:0071345* | cellular response to cytokine stimulus | 65/3240 | 9.32186E-09 |
| *GO:0006954* | inflammatory response | 64/3240 | 9.32186E-09 |
| *GO:0009607* | response to biotic stimulus | 125/3240 | 9.32186E-09 |
| *GO:0002682* | regulation of immune system process | 127/3240 | 1.14222E-08 |
| *GO:0043207* | response to external biotic stimulus | 123/3240 | 2.08336E-08 |
| *GO:0051707* | response to other organism | 123/3240 | 2.08336E-08 |
| *GO:0046649* | lymphocyte activation | 57/3240 | 2.08336E-08 |
| *GO:0045321* | leukocyte activation | 64/3240 | 4.60256E-08 |
| *GO:1903900* | regulation of viral life cycle | 22/3240 | 8.33545E-08 |
| *GO:0001775* | cell activation | 71/3240 | 8.88109E-08 |
| *GO:0019221* | cytokine-mediated signaling pathway | 58/3240 | 1.85703E-07 |
| *GO:0002684* | positive regulation of immune system process | 102/3240 | 2.24113E-07 |
| *GO:0045069* | regulation of viral genome replication | 16/3240 | 4.57965E-07 |
| *GO:0045071* | negative regulation of viral genome replication | 16/3240 | 4.57965E-07 |
| *GO:0098542* | defense response to other organism | 86/3240 | 5.54524E-07 |
| *GO:0045087* | innate immune response | 79/3240 | 5.60973E-07 |
| *GO:0051249* | regulation of lymphocyte activation | 40/3240 | 6.15234E-07 |
| *GO:0050792* | regulation of viral process | 22/3240 | 8.88937E-07 |
| *GO:0031347* | regulation of defense response | 56/3240 | 1.10142E-06 |
| *GO:0042110* | T cell activation | 36/3240 | 1.30791E-06 |
| *GO:0050865* | regulation of cell activation | 46/3240 | 1.30791E-06 |
| *GO:0048525* | negative regulation of viral process | 16/3240 | 1.30791E-06 |
| *GO:0032101* | regulation of response to external stimulus | 71/3240 | 1.3111E-06 |
| *GO:0140546* | defense response to symbiont | 83/3240 | 2.32734E-06 |
| *GO:0002694* | regulation of leukocyte activation | 44/3240 | 2.82908E-06 |
| *GO:0007159* | leukocyte cell-cell adhesion | 29/3240 | 3.60493E-06 |
| *GO:0019079* | viral genome replication | 16/3240 | 3.7628E-06 |
| *GO:0002221* | pattern recognition receptor signaling pathway | 23/3240 | 1.58929E-05 |
| *GO:0080134* | regulation of response to stress | 72/3240 | 1.61404E-05 |
| *GO:1902531* | regulation of intracellular signal transduction | 130/3240 | 2.75012E-05 |
| *GO:0032943* | mononuclear cell proliferation | 25/3240 | 3.15823E-05 |
| *GO:0046651* | lymphocyte proliferation | 25/3240 | 3.15823E-05 |
| *GO:0070661* | leukocyte proliferation | 25/3240 | 3.15823E-05 |
| *GO:0032103* | positive regulation of response to external stimulus | 44/3240 | 3.27825E-05 |
| *GO:0051604* | protein maturation | 89/3240 | 3.5489E-05 |
| *GO:0019058* | viral life cycle | 23/3240 | 4.14812E-05 |
| *GO:0002831* | regulation of response to biotic stimulus | 38/3240 | 4.14812E-05 |
| *GO:0050900* | leukocyte migration | 32/3240 | 4.40037E-05 |
| *GO:0098609* | cell-cell adhesion | 100/3240 | 4.5953E-05 |
| *GO:0030198* | extracellular matrix organization | 49/3240 | 4.76214E-05 |
| *GO:0045229* | external encapsulating structure organization | 49/3240 | 4.76214E-05 |
| *GO:0034340* | response to type I interferon | 12/3240 | 5.43141E-05 |
| *GO:0032944* | regulation of mononuclear cell proliferation | 23/3240 | 6.02584E-05 |
| *GO:0050670* | regulation of lymphocyte proliferation | 23/3240 | 6.02584E-05 |
| *GO:0070663* | regulation of leukocyte proliferation | 23/3240 | 6.02584E-05 |
| *GO:0002833* | positive regulation of response to biotic stimulus | 34/3240 | 6.9578E-05 |
| *GO:0043062* | extracellular structure organization | 49/3240 | 7.22688E-05 |
| *GO:0002218* | activation of innate immune response | 29/3240 | 9.35286E-05 |
| *GO:0009117* | nucleotide metabolic process | 67/3240 | 0.000117233 |
| *GO:0045088* | regulation of innate immune response | 35/3240 | 0.000118688 |
| *GO:0022407* | regulation of cell-cell adhesion | 27/3240 | 0.000144284 |

| | | | |
|---|---|---|---|
| *GO:0140888* | interferon-mediated signaling pathway | 12/3240 | 0.000144284 |
| *GO:1903037* | regulation of leukocyte cell-cell adhesion | 23/3240 | 0.000144284 |
| *GO:0006457* | protein folding | 45/3240 | 0.000144284 |
| *GO:0045089* | positive regulation of innate immune response | 32/3240 | 0.000144496 |
| *GO:0050776* | regulation of immune response | 81/3240 | 0.000159544 |
| *GO:0060337* | type I interferon-mediated signaling pathway | 11/3240 | 0.000162201 |
| *GO:0071357* | cellular response to type I interferon | 11/3240 | 0.000162201 |
| *GO:0031663* | lipopolysaccharide-mediated signaling pathway | 9/3240 | 0.000163774 |
| *GO:0051851* | host-mediated perturbation of symbiont process | 10/3240 | 0.00017446 |
| *GO:0050863* | regulation of T cell activation | 25/3240 | 0.000203262 |
| *GO:0031349* | positive regulation of defense response | 34/3240 | 0.000290458 |
| *GO:0032496* | response to lipopolysaccharide | 20/3240 | 0.000302487 |
| *GO:0002237* | response to molecule of bacterial origin | 21/3240 | 0.000311297 |
| *GO:0050778* | positive regulation of immune response | 72/3240 | 0.000329125 |
| *GO:0007155* | cell adhesion | 138/3240 | 0.000356704 |
| *GO:0051250* | negative regulation of lymphocyte activation | 17/3240 | 0.00037093 |
| *GO:0002263* | cell activation involved in immune response | 20/3240 | 0.000469163 |
| *GO:0035456* | response to interferon-beta | 9/3240 | 0.00062404 |
| *GO:0046597* | host-mediated suppression of symbiont invasion | 9/3240 | 0.00062404 |
| *GO:0042098* | T cell proliferation | 17/3240 | 0.00062404 |
| *GO:0042129* | regulation of T cell proliferation | 17/3240 | 0.00062404 |
| *GO:0002683* | negative regulation of immune system process | 28/3240 | 0.000652741 |
| *GO:0002695* | negative regulation of leukocyte activation | 18/3240 | 0.000652741 |
| *GO:0050866* | negative regulation of cell activation | 18/3240 | 0.000652741 |
| *GO:0071216* | cellular response to biotic stimulus | 20/3240 | 0.00068133 |
| *GO:0044093* | positive regulation of molecular function | 89/3240 | 0.00068133 |
| *GO:0030155* | regulation of cell adhesion | 36/3240 | 0.000793313 |
| *GO:0019693* | ribose phosphate metabolic process | 48/3240 | 0.000913754 |
| *GO:0002253* | activation of immune response | 57/3240 | 0.000921153 |
| *GO:0071222* | cellular response to lipopolysaccharide | 17/3240 | 0.00096274 |
| *GO:0009259* | ribonucleotide metabolic process | 46/3240 | 0.00096274 |
| *GO:0071219* | cellular response to molecule of bacterial origin | 18/3240 | 0.000992347 |
| *GO:0002366* | leukocyte activation involved in immune response | 19/3240 | 0.000997206 |
| *GO:0048585* | negative regulation of response to stimulus | 98/3240 | 0.001107309 |
| *GO:0006163* | purine nucleotide metabolic process | 54/3240 | 0.001368296 |
| *GO:0050864* | regulation of B cell activation | 16/3240 | 0.001446125 |
| *GO:0002758* | innate immune response-activating signaling pathway | 25/3240 | 0.001487243 |
| *GO:0002224* | toll-like receptor signaling pathway | 13/3240 | 0.001737891 |
| *GO:0030595* | leukocyte chemotaxis | 22/3240 | 0.001871794 |
| *GO:0008219* | cell death | 110/3240 | 0.001871794 |
| *GO:0012501* | programmed cell death | 110/3240 | 0.001871794 |
| *GO:0032945* | negative regulation of mononuclear cell proliferation | 11/3240 | 0.001871794 |
| *GO:0050672* | negative regulation of lymphocyte proliferation | 11/3240 | 0.001871794 |
| *GO:0070664* | negative regulation of leukocyte proliferation | 11/3240 | 0.001871794 |

**Supplementary Table S7.** *Relevant Differentially Expressed Genes for Consolidation versus Healthy Uninfected Comparisons.*

| ***Gene*** | ***Change*** | ***log2 F.C.*** | ***p.adjusted*** |
|---|---|---|---|
| *CLEC12B* | Increased | 21.566 | 5.21539E-13 |
| *ZBP1* | Increased | 6.906 | 1.31062E-11 |
| *ISG15* | Increased | 7.689 | 1.31062E-11 |
| *OAS1Y* | Increased | 5.807 | 1.16066E-10 |
| *OAS1Z* | Increased | 6.076 | 1.41796E-10 |
| *IFIT2* | Increased | 7.649 | 2.49755E-10 |
| *MX2* | Increased | 6.481 | 2.49755E-10 |
| *MX1* | Increased | 5.933 | 6.02953E-10 |
| *EPSTI1* | Increased | 4.148 | 6.99509E-10 |
| *IFI44* | Increased | 5.052 | 8.10501E-10 |
| *XAF1* | Increased | 4.417 | 9.03662E-10 |
| *CLEC4F* | Increased | 9.637 | 1.6537E-09 |
| *NCF1* | Increased | 4.716 | 1.6537E-09 |
| *IFI6* | Increased | 5.388 | 1.75785E-09 |
| *OAS2* | Increased | 5.441 | 3.15718E-09 |
| *DHX58* | Increased | 5.657 | 5.59446E-09 |
| *ISG12(B)* | Increased | 7.308 | 8.74925E-09 |
| *RSAD2* | Increased | 6.312 | 1.06298E-08 |
| *NCF4* | Increased | 3.622 | 1.69935E-08 |
| *ISG20* | Increased | 5.638 | 1.98881E-08 |
| *IFI44L* | Increased | 4.755 | 4.2613E-08 |
| *USP18* | Increased | 5.497 | 6.70977E-08 |
| *OAS1X* | Increased | 4.592 | 9.09734E-08 |
| *IFIT3* | Increased | 5.756 | 1.09699E-07 |
| *IFIT5* | Increased | 3.417 | 1.19461E-07 |
| *IFIH1* | Increased | 3.862 | 1.25588E-07 |
| *NLRC4* | Increased | 2.709 | 1.2832E-07 |
| *NOD1* | Increased | 3.151 | 2.19512E-07 |
| *CGAS* | Increased | 3.15 | 4.6792E-07 |
| *SIGLEC1* | Increased | 6.715 | 1.29322E-06 |
| *IFI16* | Increased | 2.89 | 1.74899E-06 |
| *TLR7* | Increased | 4.156 | 2.44759E-06 |
| *CMPK2* | Increased | 4.467 | 4.30565E-06 |
| *CLEC12A* | Increased | 3.669 | 5.7764E-06 |
| *CYBB* | Increased | 2.366 | 9.85023E-06 |
| *NOD2* | Increased | 2.392 | 2.19314E-05 |
| *S100A9* | Increased | 3.342 | 2.76486E-05 |
| *LCN2* | Increased | 5.463 | 3.5332E-05 |
| *TLR2* | Increased | 3.002 | 5.49039E-05 |
| *CXCL3* | Increased | 3.409 | 8.00006E-05 |
| *CXCL2* | Increased | 5.162 | 8.00006E-05 |
| *CLEC4E* | Increased | 1.984 | 0.000247279 |
| *CXCR2* | Increased | 3.748 | 0.000657483 |
| *NLRP3* | Increased | 2.461 | 0.000689706 |
| *CXCR1* | Increased | 4.021 | 0.000876188 |
| *CLEC4D* | Increased | 2.887 | 0.001065884 |
| *CXCL5* | Increased | 3.272 | 0.00126845 |
| *S100A8* | Increased | 2.593 | 0.001472722 |
| *CLEC10A* | Increased | 3.633 | 0.001567044 |
| *NLRP1* | Increased | 2.471 | 0.002773415 |
| *OLFM4* | Increased | 5.375 | 0.010356347 |
| *CLEC4A* | Increased | 1.721 | 0.011527559 |
| *AIM2* | Increased | 3.748 | 0.013355297 |
| *CXCL8* | Increased | 3.497 | 0.013589504 |
| *S100A12* | Increased | 2.659 | 0.018020758 |
| *NLRP12* | Increased | 2.895 | 0.026741223 |
| *SAA3* | Increased | 10.704 | 1.87637E-22 |
| *M-SAA3.2* | Increased | 9.803 | 1.03104E-15 |
| *TREM1* | Increased | 9.018 | 3.38886E-13 |
| *FCGR1A* | Increased | 4.712 | 3.1779E-12 |
| *COL4A5* | Decreased | -4.913 | 5.30708E-10 |
| *HP* | Increased | 6.242 | 2.24687E-09 |
| *TIMP1* | Increased | 6.493 | 8.01636E-09 |
| *COL4A3* | Decreased | -5 | 1.31671E-08 |
| *COL4A6* | Decreased | -4.55 | 1.97969E-08 |

| | | | |
|---|---|---|---|
| *C2* | Increased | 4.794 | 5.98867E-08 |
| *C5AR2* | Increased | 3.179 | 6.63531E-08 |
| *CTSL* | Increased | 6.119 | 1.43901E-07 |
| *PTX3* | Increased | 5.245 | 2.73194E-07 |
| *COL4A4* | Decreased | -3.623 | 5.30579E-06 |
| *CD68* | Increased | 3.403 | 9.62397E-06 |
| *HIF1A* | Increased | 2.858 | 1.0976E-05 |
| *MMP25* | Increased | 4.558 | 1.2949E-05 |
| *SIRPA* | Increased | 2.874 | 2.17603E-05 |
| *C5AR1* | Increased | 3.082 | 3.20748E-05 |
| *SIRPB1* | Increased | 2.764 | 4.22016E-05 |
| *CSF3* | Increased | 5.349 | 5.47754E-05 |
| *CSF3R* | Increased | 4.56 | 5.6399E-05 |
| *EMILIN2* | Increased | 3.506 | 7.15048E-05 |
| *CTSS* | Increased | 2.092 | 8.24685E-05 |
| *LDHA* | Increased | 3.245 | 8.76246E-05 |
| *AGER* | Decreased | -3.56 | 0.000102159 |
| *CD163* | Increased | 2.254 | 0.000172577 |
| *COL4A2* | Decreased | -2.801 | 0.000239818 |
| *HPX* | Increased | 5.328 | 0.000295825 |
| *CTSK* | Increased | 3.262 | 0.000340107 |
| *COL2A1* | Decreased | -8.618 | 0.000407214 |
| *PCDH20* | Decreased | -4.043 | 0.000502664 |
| *SLC2A1* | Increased | 3.111 | 0.00054469 |
| *MMP9* | Increased | 6.666 | 0.0005485 |
| *SAA2* | Increased | 7.156 | 0.000581446 |
| *BPIFA2A* | Decreased | -8.432 | 0.000586642 |
| *P4HA1* | Increased | 2.113 | 0.000837665 |
| *EGLN3* | Increased | 3 | 0.000862343 |
| *HAPLN1* | Decreased | -5.027 | 0.001344389 |
| *BNIP3* | Increased | 2.819 | 0.001394794 |
| *VCAN* | Increased | 2.838 | 0.00163004 |
| *CSF1R* | Increased | 1.999 | 0.001903709 |
| *HK2* | Increased | 1.829 | 0.002929651 |
| *COL9A3* | Decreased | -3.992 | 0.00395137 |
| *COL9A2* | Decreased | -3.972 | 0.004020591 |
| *TREM2* | Increased | 2.839 | 0.004328201 |
| *FREM3* | Decreased | -4.388 | 0.004328201 |
| *SIRPB2* | Increased | 1.998 | 0.00459037 |
| *AQP5* | Decreased | -2.185 | 0.00532828 |
| *FREM2* | Decreased | -3.424 | 0.005567491 |
| *MXRA5* | Increased | 2.961 | 0.007184133 |
| *LBP* | Increased | 4.723 | 0.008507226 |
| *SLC2A3* | Increased | 2.483 | 0.009438529 |
| *PLOD1* | Increased | 1.669 | 0.00970143 |
| *SFTPC* | Decreased | -2.745 | 0.022156718 |
| *FCGR2B* | Increased | 1.559 | 0.022255499 |
| *MMP1* | Increased | 5.189 | 0.026502823 |
| *COL9A1* | Decreased | -4.912 | 0.043960054 |
| *HOXA3* | Decreased | -2.783 | 0.000138382 |
| *BDNF* | Decreased | -2.814 | 0.00047073 |
| *FOXO4* | Decreased | -2.271 | 0.000657385 |
| *ADCY5* | Decreased | -3.38 | 0.001029021 |
| *HOXA5* | Decreased | -2.11 | 0.001617777 |
| *SOX17* | Decreased | -3.091 | 0.002494805 |
| *DSC2* | Decreased | -2.182 | 0.002974663 |
| *NTRK2* | Decreased | -3.012 | 0.003370259 |
| *SOX7* | Decreased | -2.57 | 0.004264795 |
| *FOXO6* | Decreased | -3.647 | 0.004378175 |
| *STMN2* | Decreased | -7.962 | 0.008721434 |
| *KRT79* | Decreased | -4.114 | 0.027779111 |
| *GRIN2B* | Decreased | -5.164 | 0.030238535 |
| *CTSH* | Increased | 3.217 | 9.83004E-09 |
| *ZBP1* | Increased | 6.906 | 1.31062E-11 |

**Supplementary Table 8**. *Enriched GO Terms for Grey Hepatization versus Healthy Uninfected Comparisons*.

| ID | Description | GeneRatio | p.adjust |
|---|---|---|---|
| GO:0009615 | response to virus | 41/4157 | 4.16395E-12 |
| GO:0051607 | defense response to virus | 39/4157 | 1.9638E-11 |
| GO:0002682 | regulation of immune system process | 159/4157 | 1.19546E-10 |
| GO:0002684 | positive regulation of immune system process | 132/4157 | 1.87121E-10 |
| GO:0034097 | response to cytokine | 84/4157 | 2.46984E-09 |
| GO:1901652 | response to peptide | 84/4157 | 2.46984E-09 |
| GO:0031347 | regulation of defense response | 72/4157 | 2.71982E-09 |
| GO:0002218 | activation of innate immune response | 41/4157 | 8.96663E-09 |
| GO:0002831 | regulation of response to biotic stimulus | 52/4157 | 1.07534E-08 |
| GO:0032101 | regulation of response to external stimulus | 90/4157 | 1.31239E-08 |
| GO:0002833 | positive regulation of response to biotic stimulus | 47/4157 | 1.5664E-08 |
| GO:0045088 | regulation of innate immune response | 49/4157 | 1.66673E-08 |
| GO:0044419 | biological process involved in interspecies interaction between organisms | 152/4157 | 1.66673E-08 |
| GO:0002221 | pattern recognition receptor signaling pathway | 30/4157 | 1.78787E-08 |
| GO:0045089 | positive regulation of innate immune response | 45/4157 | 1.81442E-08 |
| GO:0071345 | cellular response to cytokine stimulus | 74/4157 | 2.25527E-08 |
| GO:0009607 | response to biotic stimulus | 146/4157 | 4.40744E-08 |
| GO:0007155 | cell adhesion | 188/4157 | 6.77794E-08 |
| GO:0001775 | cell activation | 83/4157 | 9.21122E-08 |
| GO:0043207 | response to external biotic stimulus | 144/4157 | 9.21122E-08 |
| GO:0051707 | response to other organism | 144/4157 | 9.21122E-08 |
| GO:0046649 | lymphocyte activation | 64/4157 | 1.15317E-07 |
| GO:0006954 | inflammatory response | 71/4157 | 1.19086E-07 |
| GO:0002758 | innate immune response-activating signaling pathway | 37/4157 | 1.27315E-07 |
| GO:0045321 | leukocyte activation | 73/4157 | 1.27421E-07 |
| GO:0031349 | positive regulation of defense response | 47/4157 | 1.27421E-07 |
| GO:0051249 | regulation of lymphocyte activation | 47/4157 | 1.27421E-07 |
| GO:0098609 | cell-cell adhesion | 130/4157 | 1.64572E-07 |
| GO:0019221 | cytokine-mediated signaling pathway | 67/4157 | 1.94601E-07 |
| GO:0032103 | positive regulation of response to external stimulus | 56/4157 | 3.2557E-07 |
| GO:0098542 | defense response to other organism | 102/4157 | 4.06822E-07 |
| GO:0050790 | regulation of catalytic activity | 169/4157 | 4.73819E-07 |
| GO:0050865 | regulation of cell activation | 54/4157 | 5.14731E-07 |
| GO:0050776 | regulation of immune response | 106/4157 | 7.77338E-07 |
| GO:0032944 | regulation of mononuclear cell proliferation | 29/4157 | 7.77338E-07 |
| GO:0050670 | regulation of lymphocyte proliferation | 29/4157 | 7.77338E-07 |
| GO:0070663 | regulation of leukocyte proliferation | 29/4157 | 7.77338E-07 |
| GO:0002694 | regulation of leukocyte activation | 52/4157 | 7.77338E-07 |
| GO:1902531 | regulation of intracellular signal transduction | 164/4157 | 8.058E-07 |
| GO:0002253 | activation of immune response | 78/4157 | 9.44815E-07 |
| GO:0045087 | innate immune response | 92/4157 | 1.04928E-06 |
| GO:0050778 | positive regulation of immune response | 95/4157 | 1.40009E-06 |
| GO:0140546 | defense response to symbiont | 99/4157 | 1.4061E-06 |
| GO:0051239 | regulation of multicellular organismal process | 168/4157 | 2.20367E-06 |
| GO:0032943 | mononuclear cell proliferation | 30/4157 | 2.35123E-06 |
| GO:0046651 | lymphocyte proliferation | 30/4157 | 2.35123E-06 |
| GO:0070661 | leukocyte proliferation | 30/4157 | 2.35123E-06 |
| GO:0001525 | angiogenesis | 41/4157 | 2.96666E-06 |
| GO:0042110 | T cell activation | 40/4157 | 3.80225E-06 |
| GO:0002757 | immune response-activating signaling pathway | 61/4157 | 3.89419E-06 |
| GO:0080134 | regulation of response to stress | 87/4157 | 4.06168E-06 |
| GO:0030036 | actin cytoskeleton organization | 137/4157 | 4.48066E-06 |
| GO:0002764 | immune response-regulating signaling pathway | 62/4157 | 5.60663E-06 |
| GO:0007159 | leukocyte cell-cell adhesion | 32/4157 | 9.16773E-06 |
| GO:0030029 | actin filament-based process | 139/4157 | 1.46832E-05 |
| GO:0140888 | interferon-mediated signaling pathway | 14/4157 | 1.70639E-05 |
| GO:0048585 | negative regulation of response to stimulus | 128/4157 | 1.70639E-05 |
| GO:0048514 | blood vessel morphogenesis | 42/4157 | 1.99679E-05 |
| GO:0051604 | protein maturation | 107/4157 | 2.44116E-05 |
| GO:0030198 | extracellular matrix organization | 58/4157 | 2.8375E-05 |
| GO:0045229 | external encapsulating structure organization | 58/4157 | 2.8375E-05 |
| GO:0001944 | vasculature development | 45/4157 | 2.8375E-05 |
| GO:0002683 | negative regulation of immune system process | 35/4157 | 5.06381E-05 |
| GO:0034340 | response to type I interferon | 13/4157 | 5.06381E-05 |
| GO:0043062 | extracellular structure organization | 58/4157 | 5.06381E-05 |

| | | | |
|---|---|---|---|
| *GO:0051251* | positive regulation of lymphocyte activation | 29/4157 | 5.18742E-05 |
| *GO:0043086* | negative regulation of catalytic activity | 61/4157 | 5.48447E-05 |
| *GO:0072359* | circulatory system development | 69/4157 | 6.35089E-05 |
| *GO:0045069* | regulation of viral genome replication | 15/4157 | 6.56184E-05 |
| *GO:0045071* | negative regulation of viral genome replication | 15/4157 | 6.56184E-05 |
| *GO:0019220* | regulation of phosphate metabolic process | 104/4157 | 7.34791E-05 |
| *GO:0051174* | regulation of phosphorus metabolic process | 104/4157 | 7.34791E-05 |
| *GO:0001568* | blood vessel development | 42/4157 | 8.30277E-05 |
| *GO:0043549* | regulation of kinase activity | 71/4157 | 9.96805E-05 |
| *GO:0042325* | regulation of phosphorylation | 97/4157 | 0.000104308 |
| *GO:0009968* | negative regulation of signal transduction | 101/4157 | 0.000136928 |
| *GO:0060337* | type I interferon-mediated signaling pathway | 12/4157 | 0.00014147 |
| *GO:0071357* | cellular response to type I interferon | 12/4157 | 0.00014147 |
| *GO:0009967* | positive regulation of signal transduction | 112/4157 | 0.000160422 |
| *GO:0035239* | tube morphogenesis | 42/4157 | 0.000161788 |
| *GO:0048525* | negative regulation of viral process | 15/4157 | 0.000172317 |
| *GO:0002696* | positive regulation of leukocyte activation | 33/4157 | 0.000172871 |
| *GO:0050867* | positive regulation of cell activation | 33/4157 | 0.000172871 |
| *GO:0030155* | regulation of cell adhesion | 44/4157 | 0.000178951 |
| *GO:0032496* | response to lipopolysaccharide | 23/4157 | 0.000178951 |
| *GO:0048646* | anatomical structure formation involved in morphogenesis | 65/4157 | 0.000178951 |
| *GO:0042098* | T cell proliferation | 20/4157 | 0.000178951 |
| *GO:0042129* | regulation of T cell proliferation | 20/4157 | 0.000178951 |
| *GO:0051338* | regulation of transferase activity | 72/4157 | 0.000220185 |
| *GO:0002237* | response to molecule of bacterial origin | 24/4157 | 0.000246709 |
| *GO:0010648* | negative regulation of cell communication | 101/4157 | 0.000352347 |
| *GO:0022407* | regulation of cell-cell adhesion | 30/4157 | 0.000361914 |
| *GO:0050863* | regulation of T cell activation | 28/4157 | 0.00039082 |
| *GO:0062207* | regulation of pattern recognition receptor signaling pathway | 11/4157 | 0.00039082 |
| *GO:0019079* | viral genome replication | 15/4157 | 0.00039082 |
| *GO:0023057* | negative regulation of signaling | 101/4157 | 0.000399757 |
| *GO:1903900* | regulation of viral life cycle | 19/4157 | 0.00041081 |
| *GO:0035295* | tube development | 44/4157 | 0.000426143 |
| *GO:0007169* | cell surface receptor protein tyrosine kinase signaling pathway | 89/4157 | 0.00048274 |
| *GO:1903037* | regulation of leukocyte cell-cell adhesion | 25/4157 | 0.000505712 |

**Supplementary Table S9.** *Discussion Relevant Differentially Expressed Genes for Grey Hepatization versus Healthy Uninfected Comparisons.*

| ***Gene*** | ***Change*** | ***log2 F.C.*** | ***p.adjusted*** |
|---|---|---|---|
| *OAS1Z* | Increased | 5.992 | 1.33E-10 |
| *ZBP1* | Increased | 6.156 | 1.24E-09 |
| *ISG12(B)* | Increased | 7.544 | 1.70E-09 |
| *IFIT2* | Increased | 6.979 | 5.62E-09 |
| *MX2* | Increased | 5.862 | 7.97E-09 |
| *OAS1Y* | Increased | 5.095 | 1.29E-08 |
| *MX1* | Increased | 5.313 | 2.32E-08 |
| *IFITM2* | Increased | 4.869 | 6.47E-08 |
| *OAS2* | Increased | 4.91 | 7.03E-08 |
| *EPSTI1* | Increased | 3.576 | 8.82E-08 |
| *DHX58* | Increased | 5.103 | 1.17E-07 |
| *XAF1* | Increased | 3.772 | 1.47E-07 |
| *IFITM1* | Increased | 4.512 | 2.55E-07 |
| *TLR7* | Increased | 4.36 | 3.50E-07 |
| *OAS1X* | Increased | 4.246 | 5.05E-07 |
| *RSAD2* | Increased | 5.314 | 1.33E-06 |
| *IFIT3* | Increased | 5.015 | 2.66E-06 |
| *TLR2* | Increased | 3.354 | 2.68E-06 |
| *IFIH1* | Increased | 3.347 | 3.43E-06 |
| *USP18* | Increased | 4.552 | 6.22E-06 |
| *CGAS* | Increased | 2.449 | 7.61E-05 |
| *NLRP12* | Increased | 4.061 | 6.89E-04 |
| *CMPK2* | Increased | 2.942 | 2.53E-03 |
| *AIM2* | Increased | 4.053 | 4.78E-03 |
| *IFITM10* | Increased | 4.196 | 3.49E-02 |
| *TNIP3* | Increased | 7.626 | 5.09E-12 |
| *NCF1* | Increased | 4.975 | 9.99E-11 |
| *LCN2* | Increased | 7.034 | 2.59E-08 |
| *NCF4* | Increased | 3.479 | 4.34E-08 |
| *S100A9* | Increased | 4.113 | 6.96E-08 |
| *CYBB* | Increased | 2.672 | 2.16E-07 |
| *DDIT4* | Increased | 3.771 | 2.15E-06 |
| *CXCL2* | Increased | 5.889 | 2.72E-06 |
| *CXCR2* | Increased | 4.646 | 8.43E-06 |
| *S100A8* | Increased | 3.386 | 1.04E-05 |
| *CXCL5* | Increased | 4.114 | 1.70E-05 |
| *OLFM4* | Increased | 8.247 | 1.77E-05 |
| *TNIP1* | Increased | 2.798 | 2.71E-05 |
| *CXCR1* | Increased | 4.742 | 3.50E-05 |
| *S100A12* | Increased | 4.151 | 4.90E-05 |
| *CXCL8* | Increased | 5.08 | 9.46E-05 |
| *SOCS1* | Increased | 3.958 | 1.58E-04 |
| *NFKBIA* | Increased | 2.174 | 6.34E-04 |
| *TNFAIP3* | Increased | 2.301 | 9.42E-04 |
| *SOCS3* | Increased | 2.705 | 1.12E-03 |
| *IKBKE* | Increased | 2.283 | 9.74E-03 |
| *CXCL3* | Increased | 2.063 | 1.94E-02 |
| *SAA3* | Increased | 11.188 | 2.39E-25 |
| *M-SAA3.2* | Increased | 9.814 | 4.16E-16 |
| *TREM1* | Increased | 9.261 | 1.70E-14 |
| *FCGR1A* | Increased | 5.047 | 1.83E-14 |
| *OSMR* | Increased | 3.04 | 1.49E-13 |
| *HP* | Increased | 6.293 | 1.04E-09 |
| *C5AR2* | Increased | 3.494 | 1.26E-09 |
| *TYROBP* | Increased | 2.785 | 1.39E-07 |
| *SIRPB1* | Increased | 3.352 | 2.23E-07 |
| *PTX3* | Increased | 5.164 | 2.42E-07 |
| *C2* | Increased | 4.478 | 2.76E-07 |
| *CSF3R* | Increased | 5.561 | 2.94E-07 |
| *C5AR1* | Increased | 3.572 | 5.05E-07 |
| *CD68* | Increased | 3.718 | 5.06E-07 |
| *SIRPA* | Increased | 2.962 | 5.72E-06 |
| *HPX* | Increased | 5.694 | 5.54E-05 |

| | | | |
|---|---|---|---|
| *TREM2* | Increased | 3.546 | 1.43E-04 |
| *CD163* | Increased | 2.192 | 1.57E-04 |
| *CSF1R* | Increased | 2.118 | 5.77E-04 |
| *LBP* | Increased | 5.748 | 6.20E-04 |
| *FCGR2B* | Increased | 2.051 | 1.12E-03 |
| *SAA2* | Increased | 6.559 | 1.24E-03 |
| *FCGR2A* | Increased | 2.032 | 2.61E-03 |
| *C9* | Increased | 8.299 | 5.90E-03 |
| *CTSL* | Increased | 7.145 | 2.60E-10 |
| *CTSH* | Increased | 3.233 | 5.28E-09 |
| *TIMP1* | Increased | 6.445 | 6.74E-09 |
| *MMP25* | Increased | 5.137 | 3.30E-07 |
| *P4HA1* | Increased | 2.897 | 1.18E-06 |
| *CTSK* | Increased | 3.959 | 4.78E-06 |
| *EMILIN2* | Increased | 3.846 | 5.77E-06 |
| *MMP9* | Increased | 8.001 | 1.28E-05 |
| *MEDAG* | Increased | 2.802 | 1.69E-05 |
| *VCAN* | Increased | 3.409 | 6.21E-05 |
| *MMP1* | Increased | 8.425 | 6.79E-05 |
| *MMP14* | Increased | 2.747 | 6.82E-05 |
| *SULF1* | Increased | 3.974 | 1.02E-04 |
| *CTSZ* | Increased | 2.613 | 1.66E-04 |
| *PLOD2* | Increased | 4.192 | 1.69E-04 |
| *PLOD1* | Increased | 2.037 | 7.28E-04 |
| *MXRA5* | Increased | 3.487 | 7.42E-04 |
| *FAP* | Increased | 4.404 | 1.66E-03 |
| *MMP13* | Increased | 4.25 | 2.69E-03 |
| *TGFBI* | Increased | 2.065 | 3.19E-03 |
| *CTHRC1* | Increased | 3.466 | 4.49E-02 |
| *LDHA* | Increased | 4.288 | 5.51E-08 |
| *HIF1A* | Increased | 3.206 | 3.04E-07 |
| *HSPE1* | Increased | 2.51 | 1.19E-06 |
| *SEMA4D* | Increased | 3.937 | 1.53E-06 |
| *BNIP3* | Increased | 3.852 | 3.30E-06 |
| *VCAM1* | Increased | 3.031 | 7.41E-06 |
| *SLC2A1* | Increased | 3.809 | 8.21E-06 |
| *HSPH1* | Increased | 3.163 | 9.66E-06 |
| *PROK2* | Increased | 5.412 | 1.04E-05 |
| *EGLN3* | Increased | 3.721 | 1.24E-05 |
| *GDF10* | Decreased | -4.657 | 1.63E-05 |
| *ENO1* | Increased | 2.859 | 2.82E-05 |
| *NID2* | Increased | 2.351 | 3.95E-05 |
| *SESN2* | Increased | 3.265 | 5.54E-05 |
| *SLC2A3* | Increased | 3.522 | 6.50E-05 |
| *HK2* | Increased | 2.227 | 1.18E-04 |
| *IL21R* | Increased | 3.452 | 1.87E-04 |
| *RASA4B* | Increased | 3.284 | 1.99E-04 |
| *DDIT3* | Increased | 1.854 | 3.37E-04 |
| *FGF23* | Increased | 6.081 | 5.41E-04 |
| *ALDOC* | Increased | 2.55 | 5.54E-04 |
| *CREB3L1* | Increased | 3.292 | 1.00E-03 |
| *GDF15* | Increased | 3.769 | 1.01E-02 |
| *THBS2* | Increased | 3.307 | 1.95E-02 |
| *HSPA1A* | Increased | 2.264 | 2.11E-02 |
| *ETV3L* | Increased | 4.382 | 3.29E-02 |
| *SEMA3A* | Increased | 2.038 | 4.24E-02 |
| *IL11* | Increased | 2.806 | 4.55E-02 |
| *GUCY2D* | Increased | 3.768 | 4.60E-02 |
| *AGER* | Decreased | -5.074 | 5.42E-09 |
| *AQP5* | Decreased | -3.465 | 1.69E-06 |
| *CLDN5* | Decreased | -4.135 | 1.98E-06 |
| *SLC22A31* | Decreased | -3.977 | 2.07E-06 |
| *SLC16A11* | Decreased | -2.347 | 5.99E-05 |
| *SLC45A1* | Decreased | -5.154 | 9.20E-05 |
| *SFTPC* | Decreased | -4.157 | 1.43E-04 |
| *BPIFA2A* | Decreased | -8.509 | 3.06E-04 |
| *SCGB1D* | Decreased | -5.189 | 3.98E-04 |
| *SCNN1B* | Decreased | -3.461 | 1.31E-03 |
| *SPINK5* | Decreased | -4.55 | 6.98E-03 |
| *CLDN18* | Decreased | -2.185 | 7.08E-03 |

| | | | |
|---|---|---|---|
| *SLC34A2* | Decreased | -2.143 | 8.49E-03 |
| *SLC5A9* | Decreased | -2.598 | 9.38E-03 |
| *SLC22A17* | Decreased | -2.075 | 1.43E-02 |
| *SLC9A4* | Decreased | -4.186 | 2.66E-02 |
| *MUC4* | Decreased | -4.982 | 3.09E-02 |
| *SCGB2A2* | Unchanged | -1.544 | 1.80E-01 |
| *COL4A5* | Decreased | -5.778 | 4.50E-14 |
| *COL4A3* | Decreased | -5.932 | 5.09E-12 |
| *SPOCK2* | Decreased | -3.947 | 1.25E-10 |
| *MXRA7* | Decreased | -3.944 | 5.45E-08 |
| *COL25A1* | Decreased | -4.557 | 1.43E-06 |
| *COL27A1* | Decreased | -3.284 | 3.74E-06 |
| *COL15A1* | Decreased | -3.785 | 1.12E-05 |
| *COL21A1* | Decreased | -3.781 | 1.83E-05 |
| *WFDC1* | Decreased | -3.144 | 2.76E-05 |
| *COL13A1* | Decreased | -4.508 | 3.27E-05 |
| *WFIKKN2* | Decreased | -5.761 | 6.34E-05 |
| *FREM2* | Decreased | -4.564 | 9.12E-05 |
| *EMID1* | Decreased | -2.819 | 2.23E-04 |
| *COL26A1* | Decreased | -3.718 | 2.42E-04 |
| *COL6A6* | Decreased | -4.642 | 2.67E-04 |
| *HMCN2* | Decreased | -4.3 | 5.22E-04 |
| *COL24A1* | Decreased | -4.247 | 9.23E-04 |
| *COL2A1* | Decreased | -7.68 | 9.69E-04 |
| *HMCN1* | Decreased | -2.878 | 1.90E-03 |
| *FREM1* | Decreased | -2.532 | 2.53E-03 |
| *SPOCK1* | Decreased | -2.823 | 2.81E-03 |
| *COL9A3* | Decreased | -3.958 | 2.91E-03 |
| *COL9A2* | Decreased | -3.907 | 3.21E-03 |
| *COL9A1* | Decreased | -6.78 | 3.26E-03 |
| *COL4A1* | Decreased | -2.082 | 3.35E-03 |
| *COL5A3* | Decreased | -2.233 | 4.17E-03 |
| *FREM3* | Decreased | -4.175 | 4.76E-03 |
| *COL11A2* | Decreased | -3.53 | 7.39E-03 |
| *COLQ* | Decreased | -3.964 | 2.07E-02 |
| *PRELP* | Decreased | -2.073 | 4.52E-02 |
| *BMP5* | Decreased | -4.328 | 1.67E-09 |
| *BMP6* | Decreased | -4.018 | 9.37E-07 |
| *TBX2* | Decreased | -3.446 | 9.73E-07 |
| *FGFR3* | Decreased | -4.063 | 1.61E-06 |
| *HOXA5* | Decreased | -2.927 | 3.34E-06 |
| *PRDM6* | Decreased | -3.152 | 5.72E-06 |
| *FGFR4* | Decreased | -4.553 | 1.22E-05 |
| *SOX7* | Decreased | -3.604 | 1.67E-05 |
| *SOX17* | Decreased | -3.978 | 3.97E-05 |
| *HOXA3* | Decreased | -2.883 | 4.13E-05 |
| *NKD1* | Decreased | -3.248 | 7.96E-05 |
| *SOX4* | Decreased | -2.626 | 8.00E-05 |
| *WNT7A* | Decreased | -3.654 | 8.64E-05 |
| *TCF21* | Decreased | -3.15 | 8.77E-05 |
| *WNT9A* | Decreased | -3.238 | 1.72E-04 |
| *WNT11* | Decreased | -3.838 | 1.79E-04 |
| *FGFR1* | Decreased | -2.113 | 1.85E-04 |
| *AXIN2* | Decreased | -3.191 | 1.87E-04 |
| *GATA6* | Decreased | -2.029 | 3.95E-04 |
| *HOXB3* | Decreased | -2.35 | 4.60E-04 |
| *TBX3* | Decreased | -2.46 | 5.37E-04 |
| *IRX3* | Decreased | -2.107 | 6.81E-04 |
| *IRX5* | Decreased | -2.118 | 8.68E-04 |
| *GATA5* | Decreased | -3.266 | 8.81E-04 |
| *FGFR2* | Decreased | -2.311 | 9.49E-04 |
| *GATA3* | Decreased | -2.269 | 1.01E-03 |
| *TBX4* | Decreased | -2.365 | 1.14E-03 |
| *BMPER* | Decreased | -2.426 | 1.63E-03 |
| *IRX2* | Decreased | -2.046 | 1.81E-03 |
| *HOXB5* | Decreased | -2.559 | 2.18E-03 |
| *IRX1* | Decreased | -2.45 | 2.31E-03 |
| *PRDM16* | Decreased | -2.605 | 2.33E-03 |
| *NKD2* | Decreased | -2.561 | 2.36E-03 |
| *FGF1* | Decreased | -2.108 | 4.86E-03 |

| | | | |
|---|---|---|---|
| *HOXA2* | Decreased | -2.552 | 6.50E-03 |
| *FGF12* | Decreased | -2.402 | 7.67E-03 |
| *SOX13* | Decreased | -2.322 | 8.51E-03 |
| *HOXB6* | Decreased | -2.214 | 9.08E-03 |
| *WNT2* | Decreased | -2.163 | 1.27E-02 |
| *WNT8B* | Decreased | -3.35 | 2.32E-02 |
| *NOTUM* | Decreased | -3.725 | 2.52E-02 |
| *TBX1* | Decreased | -2.099 | 3.33E-02 |
| *FGF20* | Decreased | -3.244 | 3.95E-02 |
| *GPRASP1* | Decreased | -3.365 | 9.88E-09 |
| *GPRASP2* | Decreased | -2.492 | 1.29E-08 |
| *CNR1* | Decreased | -7.277 | 1.25E-07 |
| *RIMS3* | Decreased | -3.683 | 7.66E-07 |
| *ADCY5* | Decreased | -4.772 | 1.98E-06 |
| *GRIN3A* | Decreased | -4.358 | 1.09E-05 |
| *CNR2* | Decreased | -3.023 | 3.60E-05 |
| *GPR37* | Decreased | -4.5 | 5.80E-05 |
| *PDYN* | Decreased | -4.913 | 5.54E-04 |
| *CHRNB3* | Decreased | -5.183 | 6.73E-04 |
| *SYNDIG1L* | Decreased | -2.49 | 7.87E-04 |
| *ADCYAP1R1* | Decreased | -3.727 | 8.00E-04 |
| *GPR55* | Decreased | -3.381 | 8.98E-04 |
| *GAP43* | Decreased | -6.838 | 1.40E-03 |
| *STMN2* | Decreased | -8.574 | 2.92E-03 |
| *RIMS1* | Decreased | -3.918 | 3.05E-03 |
| *SYT13* | Decreased | -6.516 | 5.17E-03 |
| *GRIN1* | Decreased | -2.519 | 8.72E-03 |
| *NTM* | Decreased | -7.39 | 1.38E-02 |
| *GRIN2B* | Decreased | -5.462 | 1.55E-02 |
| *SYT7* | Decreased | -2.243 | 2.29E-02 |
| *SYT8* | Decreased | -4.376 | 2.53E-02 |
| *GALR3* | Decreased | -2.885 | 3.23E-02 |
| *CRHR2* | Decreased | -3.895 | 3.28E-02 |
| *SCGN* | Decreased | -4.119 | 3.63E-02 |
| *CHRNB4* | Decreased | -4.041 | 4.99E-02 |
| *GSTA3* | Decreased | -7.536 | 3.54E-08 |
| *CYP1A1* | Decreased | -9.384 | 6.84E-08 |
| *GSTT2* | Decreased | -4.433 | 1.44E-07 |
| *ABCC6* | Decreased | -4.355 | 4.87E-07 |
| *TRPV4* | Decreased | -3.006 | 9.87E-07 |
| *AQP5* | Decreased | -3.465 | 1.69E-06 |
| *SLC22A31* | Decreased | -3.977 | 2.07E-06 |
| *CYP2A13* | Decreased | -6.476 | 2.75E-06 |
| *TRPC3* | Decreased | -3.529 | 1.06E-05 |
| *CNGA2* | Decreased | -5.899 | 1.17E-05 |
| *HPGD* | Decreased | -4.564 | 1.69E-05 |
| *UGT1A6* | Decreased | -7.163 | 1.89E-05 |
| *TRPM4* | Decreased | -3.236 | 3.10E-05 |
| *CYP4B1* | Decreased | -3.892 | 4.72E-05 |
| *GSTA2* | Decreased | -4.298 | 1.06E-04 |
| *ABCB1* | Decreased | -3.11 | 2.77E-04 |
| *CYP2C90* | Decreased | -4.468 | 5.50E-04 |
| *SLC16A9* | Decreased | -3.301 | 8.26E-04 |
| *GSTA1* | Decreased | -3.876 | 9.13E-04 |
| *P2RX1* | Decreased | -2.708 | 1.18E-03 |
| *KCNB2* | Decreased | -3.437 | 1.48E-03 |
| *KCNK3* | Decreased | -4.047 | 4.61E-03 |
| *KCND3* | Decreased | -3.373 | 6.57E-03 |
| *P2RX5* | Decreased | -1.484 | 7.34E-03 |
| *AQP1* | Decreased | -2.087 | 8.38E-03 |
| *UGT2A1* | Decreased | -3.721 | 8.84E-03 |
| *GSTM3* | Decreased | -2.206 | 9.05E-03 |
| *SLC5A9* | Decreased | -2.598 | 9.38E-03 |
| *SCN11A* | Decreased | -5.913 | 9.50E-03 |
| *KCNH2* | Decreased | -2.998 | 9.71E-03 |
| *CYP4A22* | Decreased | -6.401 | 1.31E-02 |
| *CYP17A1* | Decreased | -2.331 | 1.38E-02 |
| *SLC24A3* | Decreased | -1.91 | 1.80E-02 |
| *SCN3B* | Decreased | -3.378 | 1.95E-02 |
| *SLC6A1* | Decreased | -2.808 | 2.05E-02 |

| | | | |
|---|---|---|---|
| *FMO5* | Decreased | -1.259 | 2.31E-02 |
| *SLC8A3* | Decreased | -2.804 | 2.40E-02 |
| *ASIC2* | Decreased | -3.12 | 3.25E-02 |
| *CYP2C87* | Decreased | -4.292 | 3.47E-02 |
| *ASIC3* | Decreased | -3.276 | 4.36E-02 |
| *CYP1A2* | Decreased | -4.603 | 5.94E-02 |
| *CCR1* | Increased | 4.995 | 1.04E-08 |
| *SPI1* | Increased | 3.439 | 6.00E-08 |
| *IL7R* | Increased | 5.065 | 7.74E-08 |
| *CD69* | Increased | 4.314 | 1.60E-05 |
| *POU2F2* | Increased | 3.219 | 2.53E-05 |
| *CCR2* | Increased | 2.785 | 3.27E-05 |
| *IL23R* | Increased | 4.004 | 3.95E-05 |
| *CCR5* | Increased | 3.006 | 6.02E-05 |
| *PRDM1* | Increased | 2.132 | 1.22E-04 |
| *CD274* | Increased | 3.27 | 2.43E-04 |
| *CTLA4* | Increased | 2.384 | 8.98E-04 |
| *LAG3* | Increased | 3.118 | 4.81E-03 |
| *BATF* | Increased | 2.476 | 5.66E-03 |
| *CD1B* | Decreased | -4.734 | 5.75E-03 |
| *CXCL10* | Increased | 2.351 | 1.33E-02 |
| *BOLA-DQA2* | Decreased | -7.688 | 4.40E-02 |
| *CD1A* | Decreased | -3.024 | 5.33E-02 |
| *BOLA-DQA5* | Decreased | -3.644 | 1.11E-01 |
| *EMILIN3* | Decreased | -10.069 | 1.25E-10 |
| *CYM* | Decreased | -12.256 | 3.55E-07 |
| *CLEC12B* | Increased | 16.459 | 9.12E-08 |

**Supplementary Table S10**. *Enriched GO Terms for Necrosis/Sequestra versus Healthy Uninfected Comparisons*.

| ***ID*** | ***Description*** | ***GeneRatio*** | ***p.adjust*** |
|---|---|---|---|
| *GO:0007155* | cell adhesion | 295/7188 | 5.84824E-09 |
| *GO:0030036* | actin cytoskeleton organization | 221/7188 | 3.35736E-08 |
| *GO:0030029* | actin filament-based process | 226/7188 | 8.42582E-08 |
| *GO:0098609* | cell-cell adhesion | 196/7188 | 2.56612E-07 |
| *GO:0051607* | defense response to virus | 43/7188 | 6.53806E-07 |
| *GO:0022008* | neurogenesis | 273/7188 | 6.53806E-07 |
| *GO:0009615* | response to virus | 44/7188 | 1.00697E-06 |
| *GO:0048646* | anatomical structure formation involved in morphogenesis | 103/7188 | 5.9788E-06 |
| *GO:0048699* | generation of neurons | 241/7188 | 2.13814E-05 |
| *GO:0030182* | neuron differentiation | 227/7188 | 2.21367E-05 |
| *GO:0001944* | vasculature development | 65/7188 | 2.2592E-05 |
| *GO:0032535* | regulation of cellular component size | 79/7188 | 3.1339E-05 |
| *GO:0034097* | response to cytokine | 107/7188 | 3.42487E-05 |
| *GO:1901652* | response to peptide | 107/7188 | 3.42487E-05 |
| *GO:0072359* | circulatory system development | 104/7188 | 3.8905E-05 |
| *GO:0022603* | regulation of anatomical structure morphogenesis | 93/7188 | 3.8905E-05 |
| *GO:0051239* | regulation of multicellular organismal process | 254/7188 | 4.48117E-05 |
| *GO:0071345* | cellular response to cytokine stimulus | 96/7188 | 4.51462E-05 |
| *GO:0019221* | cytokine-mediated signaling pathway | 89/7188 | 6.08328E-05 |
| *GO:0051241* | negative regulation of multicellular organismal process | 75/7188 | 6.16372E-05 |
| *GO:0000902* | cell morphogenesis | 170/7188 | 6.42577E-05 |
| *GO:0048514* | blood vessel morphogenesis | 58/7188 | 8.49715E-05 |
| *GO:0120035* | regulation of plasma membrane bounded cell projection organization | 84/7188 | 8.49715E-05 |
| *GO:0050790* | regulation of catalytic activity | 248/7188 | 0.000106328 |
| *GO:0001568* | blood vessel development | 60/7188 | 0.000110489 |
| *GO:0051128* | regulation of cellular component organization | 267/7188 | 0.00011775 |
| *GO:0001775* | cell activation | 110/7188 | 0.00011775 |
| *GO:0090066* | regulation of anatomical structure size | 97/7188 | 0.000123044 |
| *GO:0048638* | regulation of developmental growth | 32/7188 | 0.000150714 |
| *GO:0007015* | actin filament organization | 139/7188 | 0.00015188 |
| *GO:0016477* | cell migration | 203/7188 | 0.000152573 |
| *GO:0010975* | regulation of neuron projection development | 60/7188 | 0.000161376 |
| *GO:0045926* | negative regulation of growth | 26/7188 | 0.000162415 |
| *GO:0031344* | regulation of cell projection organization | 85/7188 | 0.000162415 |
| *GO:0001525* | angiogenesis | 53/7188 | 0.000162415 |
| *GO:0008154* | actin polymerization or depolymerization | 63/7188 | 0.000219426 |
| *GO:0048666* | neuron development | 180/7188 | 0.000241426 |
| *GO:0035239* | tube morphogenesis | 60/7188 | 0.000247561 |
| *GO:0048640* | negative regulation of developmental growth | 23/7188 | 0.000266566 |
| *GO:0098660* | inorganic ion transmembrane transport | 243/7188 | 0.000309814 |
| *GO:0045069* | regulation of viral genome replication | 18/7188 | 0.000309814 |
| *GO:0045071* | negative regulation of viral genome replication | 18/7188 | 0.000309814 |
| *GO:0098662* | inorganic cation transmembrane transport | 221/7188 | 0.000309814 |
| *GO:0048585* | negative regulation of response to stimulus | 190/7188 | 0.000361487 |
| *GO:1902531* | regulation of intracellular signal transduction | 238/7188 | 0.000361487 |
| *GO:0001558* | regulation of cell growth | 33/7188 | 0.00039646 |
| *GO:0032943* | mononuclear cell proliferation | 36/7188 | 0.000408748 |
| *GO:0046651* | lymphocyte proliferation | 36/7188 | 0.000408748 |
| *GO:0070661* | leukocyte proliferation | 36/7188 | 0.000408748 |
| *GO:0046649* | lymphocyte activation | 80/7188 | 0.000478347 |
| *GO:0008361* | regulation of cell size | 34/7188 | 0.000484476 |
| *GO:0040008* | regulation of growth | 40/7188 | 0.000484476 |
| *GO:0007167* | enzyme-linked receptor protein signaling pathway | 186/7188 | 0.000484476 |
| *GO:0035295* | tube development | 64/7188 | 0.000484476 |
| *GO:0098657* | import into cell | 191/7188 | 0.000484476 |
| *GO:0006954* | inflammatory response | 90/7188 | 0.000505513 |
| *GO:0045321* | leukocyte activation | 93/7188 | 0.000566567 |
| *GO:0032101* | regulation of response to external stimulus | 113/7188 | 0.000566567 |
| *GO:0030155* | regulation of cell adhesion | 62/7188 | 0.000605164 |
| *GO:0061387* | regulation of extent of cell growth | 25/7188 | 0.000605164 |
| *GO:1903900* | regulation of viral life cycle | 25/7188 | 0.000605164 |
| *GO:0030308* | negative regulation of cell growth | 23/7188 | 0.000630403 |
| *GO:0098655* | monoatomic cation transmembrane transport | 228/7188 | 0.000630403 |
| *GO:0050768* | negative regulation of neurogenesis | 21/7188 | 0.000630403 |
| *GO:0051961* | negative regulation of nervous system development | 21/7188 | 0.000630403 |

| | | | |
|---|---|---|---|
| *GO:0034220* | monoatomic ion transmembrane transport | 260/7188 | 0.000630403 |
| *GO:0032944* | regulation of mononuclear cell proliferation | 33/7188 | 0.00064824 |
| *GO:0050670* | regulation of lymphocyte proliferation | 33/7188 | 0.00064824 |
| *GO:0070663* | regulation of leukocyte proliferation | 33/7188 | 0.00064824 |
| *GO:0048870* | cell motility | 231/7188 | 0.000703822 |
| *GO:0031175* | neuron projection development | 163/7188 | 0.000720841 |
| *GO:0030042* | actin filament depolymerization | 26/7188 | 0.000828046 |
| *GO:0030041* | actin filament polymerization | 57/7188 | 0.000831033 |
| *GO:0022407* | regulation of cell-cell adhesion | 41/7188 | 0.000904688 |
| *GO:0031345* | negative regulation of cell projection organization | 24/7188 | 0.000913672 |
| *GO:0048675* | axon extension | 24/7188 | 0.000913672 |
| *GO:0048525* | negative regulation of viral process | 18/7188 | 0.000980407 |
| *GO:0010977* | negative regulation of neuron projection development | 22/7188 | 0.000980407 |
| *GO:0030516* | regulation of axon extension | 22/7188 | 0.000980407 |
| *GO:0050771* | negative regulation of axonogenesis | 20/7188 | 0.001012288 |
| *GO:0007159* | leukocyte cell-cell adhesion | 39/7188 | 0.001100967 |
| *GO:0040007* | growth | 51/7188 | 0.001100967 |
| *GO:0048589* | developmental growth | 43/7188 | 0.001151526 |
| *GO:0016049* | cell growth | 36/7188 | 0.001166944 |
| *GO:0050793* | regulation of developmental process | 210/7188 | 0.001205215 |
| *GO:0140888* | interferon-mediated signaling pathway | 15/7188 | 0.001466291 |
| *GO:0001667* | ameboidal-type cell migration | 28/7188 | 0.001470469 |
| *GO:0120039* | plasma membrane bounded cell projection morphogenesis | 117/7188 | 0.001470469 |
| *GO:0048858* | cell projection morphogenesis | 119/7188 | 0.001500439 |
| *GO:0048667* | cell morphogenesis involved in neuron differentiation | 107/7188 | 0.00162842 |
| *GO:0051336* | regulation of hydrolase activity | 128/7188 | 0.001647444 |
| *GO:0030001* | metal ion transport | 243/7188 | 0.00164779 |
| *GO:0050792* | regulation of viral process | 26/7188 | 0.00173349 |
| *GO:0060284* | regulation of cell development | 71/7188 | 0.001906396 |
| *GO:0030335* | positive regulation of cell migration | 53/7188 | 0.002010777 |
| *GO:1990138* | neuron projection extension | 27/7188 | 0.002345742 |
| *GO:0034340* | response to type I interferon | 14/7188 | 0.002739999 |
| *GO:0048812* | neuron projection morphogenesis | 114/7188 | 0.003003812 |
| *GO:0048588* | developmental cell growth | 28/7188 | 0.003003812 |
| *GO:0060560* | developmental growth involved in morphogenesis | 28/7188 | 0.003003812 |

**Table 11.** *Relevant Differentially Expressed Genes for Necrosis versus Healthy Uninfected Comparisons.*

| Gene | Change | log2 F.C. | p.adjusted |
|---|---|---|---|
| *NTN4* | Decreased | -4.263 | 2.1195E-30 |
| *AQP5* | Decreased | -4.263 | 2.1195E-30 |
| *SAA3* | Increased | 10.343 | 6.9202E-29 |
| *CLDN7* | Decreased | -4.442 | 6.9202E-29 |
| *M-SAA3.2* | Increased | 10.841 | 7.064E-26 |
| *CLIC5* | Decreased | -5.139 | 4.017E-25 |
| *CLDN4* | Decreased | -5.269 | 3.6715E-23 |
| *KRT8* | Decreased | -4.499 | 1.1097E-22 |
| *KRT19* | Decreased | -4.902 | 1.3274E-22 |
| *CTSK* | Increased | 6.584 | 3.7475E-22 |
| *CXCL17* | Decreased | -4.149 | 8.4206E-21 |
| *KRT7* | Decreased | -4.375 | 1.8601E-20 |
| *KLF5* | Decreased | -3.192 | 2.5026E-19 |
| *AGER* | Decreased | -6.164 | 6.5596E-19 |
| *CYP4B1* | Decreased | -6.638 | 1.0355E-18 |
| *CTSL* | Increased | 8.22 | 1.3294E-18 |
| *SPP1* | Increased | 10.709 | 1.3429E-18 |
| *CLDN5* | Decreased | -6.149 | 8.5965E-18 |
| *MMP13* | Increased | 9.178 | 4.6635E-17 |
| *COL4A5* | Decreased | -5.235 | 8.0345E-17 |
| *OCLN* | Decreased | -4.355 | 1.0666E-16 |
| *CLDN18* | Decreased | -5.079 | 1.9986E-16 |
| *ESAM* | Decreased | -3.873 | 2.0832E-16 |
| *DKK3* | Decreased | -2.788 | 2.5846E-16 |
| *HOPX* | Decreased | -3.548 | 4.4242E-16 |
| *UGT1A1* | Decreased | -5.017 | 7.2835E-16 |
| *SFTPB* | Decreased | -4.842 | 1.7623E-15 |
| *MMP9* | Increased | 11.494 | 3.664E-15 |
| *SFTPD* | Decreased | -4.726 | 4.9751E-15 |
| *EPCAM* | Decreased | -4.063 | 7.8579E-15 |
| *SLC6A14* | Decreased | -4.652 | 9.3967E-15 |
| *BMP2* | Decreased | -3.394 | 1.672E-14 |
| *LAMB2* | Decreased | -4.044 | 1.8756E-14 |
| *CD34* | Decreased | -3.827 | 2.1086E-14 |
| *SCGB1A1* | Decreased | -5.39 | 2.3592E-14 |
| *LAMA3* | Decreased | -3.998 | 2.5126E-14 |
| *GATA6* | Decreased | -3.454 | 3.2659E-14 |
| *ELN* | Decreased | -3.846 | 7.7832E-14 |
| *AQP1* | Decreased | -4.474 | 1.0274E-13 |
| *FMO3* | Decreased | -4.035 | 1.2015E-13 |
| *S1PR1* | Decreased | -3.622 | 1.4383E-13 |
| *MMP1* | Increased | 12.415 | 3.0352E-13 |
| *CLDN3* | Decreased | -3.875 | 3.72E-13 |
| *CDH5* | Decreased | -3.756 | 4.6676E-13 |
| *CXCL8* | Increased | 7.374 | 6.0177E-13 |
| *ISG15* | Increased | 6.533 | 6.3192E-13 |
| *WNT2B* | Decreased | -3.63 | 1.1668E-12 |
| *COL4A3* | Decreased | -4.97 | 1.2968E-12 |
| *CCND1* | Decreased | -3.671 | 1.8891E-12 |
| *IFIT2* | Increased | 6.76 | 2.3143E-12 |
| *KLF4* | Decreased | -3.367 | 2.962E-12 |
| *GJA5* | Decreased | -3.539 | 4.2606E-12 |
| *SCGB2A2* | Decreased | -5.616 | 5.1077E-12 |
| *FMO5* | Decreased | -2.852 | 7.9782E-12 |
| *AURKA* | Increased | 2.495 | 9.6536E-12 |
| *CLDN1* | Decreased | -2.964 | 1.5365E-11 |
| *KRT18* | Decreased | -3.444 | 1.6885E-11 |
| *MMP19* | Increased | 3.265 | 1.8441E-11 |
| *SLC16A11* | Decreased | -3.224 | 2.7484E-11 |
| *CYP2F1* | Decreased | -4.971 | 2.8247E-11 |
| *TEK* | Decreased | -3.648 | 3.1758E-11 |
| *BMP3* | Decreased | -3.739 | 3.4578E-11 |
| *IL7R* | Increased | 4.994 | 4.1244E-11 |
| *NKX2-1* | Decreased | -3.369 | 4.2291E-11 |
| *IFIT3* | Increased | 5.614 | 4.6976E-11 |
| *CD83* | Decreased | -2.423 | 1.9399E-10 |
| *AOX1* | Decreased | -4.322 | 2.5752E-10 |

| | | | |
|---|---|---|---|
| *SFTPC* | Decreased | -5.413 | 3.6475E-10 |
| *SFTPA1* | Decreased | -4.579 | 5.0406E-10 |
| *ANGPTL6* | Increased | 4.38 | 5.2626E-10 |
| *KCNH4* | Increased | 5.845 | 7.1929E-10 |
| *CXCL2* | Increased | 6.096 | 1.2665E-09 |
| *SCGB1D* | Decreased | -6.854 | 3.417E-09 |
| *IFT57* | Decreased | -2.17 | 4.1279E-09 |
| *CCNB2* | Increased | 2.57 | 4.3638E-09 |
| *MMP3* | Increased | 10.168 | 4.4896E-09 |
| *SLC6A20* | Decreased | -3.116 | 6.7482E-09 |
| *LAMA5* | Decreased | -3.303 | 6.8436E-09 |
| *TNFSF15* | Decreased | -3.231 | 7.6277E-09 |
| *COL4A2* | Decreased | -3.369 | 8.4221E-09 |
| *IFIH1* | Increased | 3.328 | 8.9745E-09 |
| *ILDR1* | Decreased | -3.093 | 1.1472E-08 |
| *TEAD1* | Decreased | -2.237 | 1.6429E-08 |
| *MX2* | Increased | 4.652 | 1.6636E-08 |
| *IL20RB* | Decreased | -2.529 | 1.7486E-08 |
| *COL4A6* | Decreased | -3.627 | 2.736E-08 |
| *MCM10* | Increased | 2.656 | 4.1968E-08 |
| *MMRN2* | Decreased | -3.121 | 6.6147E-08 |
| *PAX2* | Increased | 10.049 | 1.0013E-07 |
| *PIGR* | Decreased | -3.131 | 1.3026E-07 |
| *BMP4* | Decreased | -2.102 | 1.3095E-07 |
| *IFI30* | Increased | 2.508 | 1.4138E-07 |
| *UGT1A6* | Decreased | -7.259 | 1.5795E-07 |
| *IL1B* | Increased | 3.445 | 1.7086E-07 |
| *SLC22A31* | Decreased | -3.652 | 1.7533E-07 |
| *TEAD3* | Decreased | -2.321 | 1.89E-07 |
| *MX1* | Increased | 4.007 | 2.2595E-07 |
| *IL17F* | Increased | 7.041 | 2.7854E-07 |
| *FOXJ1* | Decreased | -4.195 | 3.2697E-07 |
| *ILDR2* | Decreased | -3.304 | 3.8134E-07 |
| *TSLP* | Decreased | -3.331 | 4.8064E-07 |
| *FBLN5* | Decreased | -2.84 | 4.8109E-07 |
| *LAMB3* | Decreased | -2.344 | 5.1236E-07 |
| *YAP1* | Decreased | -2.6 | 6.535E-07 |
| *FMO2* | Decreased | -3.571 | 7.1728E-07 |
| *WNT7A* | Decreased | -3.903 | 7.9332E-07 |
| *MUC1* | Decreased | -2.777 | 8.146E-07 |
| *KCNK9* | Increased | 7.695 | 1.0418E-06 |
| *USP18* | Increased | 3.969 | 1.1992E-06 |
| *AOX2* | Decreased | -4.374 | 1.5009E-06 |
| *FOXA1* | Decreased | -2.745 | 1.7079E-06 |
| *CDC6* | Increased | 2.764 | 2.0437E-06 |
| *COL4A1* | Decreased | -2.635 | 2.2001E-06 |
| *BMP6* | Decreased | -3.171 | 2.5395E-06 |
| *WNT9B* | Increased | 6.263 | 2.9532E-06 |
| *ZBP1* | Increased | 3.894 | 3.0035E-06 |
| *RASIP1* | Decreased | -3.145 | 3.0945E-06 |
| *CYP2A13* | Decreased | -5.175 | 3.8214E-06 |
| *PLK1* | Increased | 2.095 | 3.8473E-06 |
| *OAS2* | Increased | 3.439 | 4.0642E-06 |
| *OAS1Z* | Increased | 3.491 | 6.3223E-06 |
| *DKK2* | Decreased | -4.646 | 6.3952E-06 |
| *RSAD2* | Increased | 4.027 | 6.8009E-06 |
| *IL17A* | Increased | 6.368 | 8.2918E-06 |
| *OAS1X* | Increased | 3.071 | 8.9733E-06 |
| *ECM1* | Increased | 2.876 | 1.1939E-05 |
| *IRF4* | Increased | 2.973 | 1.2838E-05 |
| *MMP25* | Increased | 3.58 | 1.4884E-05 |
| *MUC15* | Decreased | -3.256 | 1.5784E-05 |
| *NOTCH3* | Decreased | -2.616 | 1.594E-05 |
| *ITIH5* | Decreased | -3.196 | 1.6088E-05 |
| *HMCN1* | Decreased | -3.15 | 1.7653E-05 |
| *IL21R* | Increased | 3.209 | 1.8073E-05 |
| *BIRC5* | Increased | 2.773 | 1.9409E-05 |
| *SPON2* | Decreased | -3.198 | 1.9524E-05 |
| *IL21* | Increased | 4.65 | 2.195E-05 |
| *AURKB* | Increased | 2.459 | 2.3075E-05 |

| | | | |
|---|---|---|---|
| *IFI44L* | Increased | 2.962 | 2.4047E-05 |
| *DNAL1* | Decreased | -1.791 | 2.5059E-05 |
| *UGT2A1* | Decreased | -4.653 | 2.8272E-05 |
| *TIE1* | Decreased | -2.44 | 2.8292E-05 |
| *OAS1Y* | Increased | 3.049 | 3.6003E-05 |
| *TEAD4* | Decreased | -2.268 | 3.7293E-05 |
| *IFI6* | Increased | 3.003 | 3.8981E-05 |
| *IFI44* | Increased | 2.765 | 3.8991E-05 |
| *MMP12* | Increased | 5.03 | 4.4509E-05 |
| *PAX1* | Increased | 8.637 | 4.9081E-05 |
| *PRELP* | Decreased | -3.114 | 7.045E-05 |
| *CCNB1* | Increased | 2.748 | 7.5391E-05 |
| *IRF1* | Increased | 2.089 | 8.4014E-05 |
| *CXCL5* | Increased | 3.06 | 9.0775E-05 |
| *TNF* | Decreased | -3.114 | 9.3236E-05 |
| *IL17RB* | Decreased | -2.822 | 0.00010792 |
| *IRF7* | Increased | 2.99 | 0.00011183 |
| *DNAH11* | Decreased | -2.77 | 0.00014858 |
| *CDC20* | Increased | 2.293 | 0.00016415 |
| *CCL8* | Increased | 3.982 | 0.00025613 |
| *TEKT3* | Decreased | -2.155 | 0.0002678 |
| *TEKT1* | Decreased | -2.777 | 0.00026959 |
| *RSPO4* | Increased | 6.311 | 0.0003113 |
| *CNTNAP2* | Increased | 3.394 | 0.00036541 |
| *MXRA5* | Increased | 2.918 | 0.00045964 |
| *BMP1* | Increased | 2.014 | 0.0005029 |
| *SCGB3A2* | Decreased | -3.131 | 0.00057163 |
| *KIF20A* | Increased | 2.236 | 0.00057713 |
| *PAX6* | Increased | 4.456 | 0.0006413 |
| *CDHR2* | Increased | 8.482 | 0.00070357 |
| *COL22A1* | Increased | 6.476 | 0.0008072 |
| *IL24* | Increased | 2.595 | 0.00092294 |
| *HOXD10* | Increased | 5.234 | 0.00095554 |
| *MMP17* | Increased | 4.925 | 0.00103251 |
| *SLC45A1* | Decreased | -3.542 | 0.00111048 |
| *GDF15* | Increased | 3.664 | 0.0011407 |
| *PLVAP* | Decreased | -2.717 | 0.00115274 |
| *CFAP47* | Decreased | -2.703 | 0.00116123 |
| *CDH18* | Increased | 8.461 | 0.00116329 |
| *DLX4* | Increased | 3.492 | 0.00120019 |
| *CD209* | Decreased | -2.388 | 0.00127577 |
| *POU3F1* | Increased | 4.362 | 0.0012807 |
| *JAG2* | Decreased | -2.704 | 0.00143543 |
| *PCDH15* | Increased | 6.257 | 0.00143881 |
| *KIF18B* | Increased | 2.23 | 0.00165451 |
| *IL23R* | Increased | 2.501 | 0.00178153 |
| *POU3F3* | Increased | 8.847 | 0.0018428 |
| *CNTN2* | Increased | 3.928 | 0.00213671 |
| *DNAL4* | Decreased | -1.459 | 0.00220723 |
| *PCDH8* | Increased | 6.457 | 0.00258589 |
| *WNT6* | Increased | 6.436 | 0.00267812 |
| *KLRK1* | Decreased | -2.012 | 0.0027162 |
| *IL22* | Increased | 6.075 | 0.00274767 |
| *DNAH5* | Decreased | -2.75 | 0.00327775 |
| *DLX3* | Increased | 3.51 | 0.00345244 |
| *PAX3* | Increased | 7.165 | 0.00356607 |
| *CYP2B6* | Decreased | -3.087 | 0.00423988 |
| *CYP1A1* | Decreased | -3.976 | 0.00441268 |
| *CCL3* | Increased | 2.472 | 0.00629054 |
| *CFAP43* | Decreased | -2.175 | 0.00670925 |
| *WNT11* | Decreased | -2.302 | 0.00748132 |
| *COL23A1* | Increased | 2.386 | 0.00841791 |
| *COL19A1* | Increased | 3.558 | 0.0090981 |
| *CFAP299* | Decreased | -2.282 | 0.01066218 |
| *HOXA10* | Increased | 7.939 | 0.01114039 |
| *IL1A* | Increased | 2.154 | 0.01247701 |
| *CFAP52* | Decreased | -2.013 | 0.01304591 |
| *BARX1* | Increased | 4.465 | 0.01305502 |
| *HOXC8* | Increased | 6.993 | 0.01340792 |
| *POU3F2* | Increased | 5.722 | 0.01486684 |

| | | | |
|---|---|---|---|
| *IL26* | Increased | 3.076 | 0.01598864 |
| *HOXA11* | Increased | 7.463 | 0.01691149 |
| *CDH9* | Unchanged | 6.921 | 0.0187118 |
| *DLX5* | Increased | 4.785 | 0.02327351 |
| *DLX1* | Increased | 7.714 | 0.0278347 |
| *WNT10A* | Increased | 3.609 | 0.02796113 |
| *HOXD13* | Increased | 7.568 | 0.0280044 |
